# Higher-Order Thalamus is Pivotal in Schizophrenia-Associated Pathophysiology

**DOI:** 10.64898/2026.01.24.699491

**Authors:** Jeffrey Stedehouder, Katerina Panti, Yangfan Peng, Charlotte J. Stagg, Andrew Sharott

## Abstract

Synaptic dysfunction has been proposed as cellular pathophysiology underlying schizophrenia, yet the brain-wide distribution of dysfunctional circuits at single-neuron resolution has remained unknown. Here, we perform comprehensive multi-probe electrophysiological investigations *in vivo* in the *Grin2a*^+/-^ preclinical model for schizophrenia and control animals, recording across ∼45 brain regions spanning cortex, striatum, hippocampus, and thalamus. Mutants displayed distributed and graded alterations across regions, with prominent activity reductions in higher-order thalamus and cross-parameter alterations across prefrontal cortices, striatum, and hippocampus. Restoration of higher-order thalamic activity in mutants was sufficient to normalize alterations in connected prefrontal cortices and striatum and unexpectedly cascaded to hippocampus and sensory cortices. Thus, higher-order thalamus plays a pivotal role in schizophrenia pathophysiology and restoration of a single informed locus could present a potent therapeutic strategy.

## Main Text

Schizophrenia is a major debilitating psychiatric disorder whose clinical presentation encompasses positive, negative, and cognitive symptoms.^1^ The neurobiological underpinnings of the disease have remained incompletely understood and novel treatment discovery slow. Convergent genetic evidence has suggested synaptic dysfunction as an overarching cellular pathophysiological mechanisms.^2,3^ However, patient neuroimaging studies have limited spatiotemporal resolution to isolate neural circuits arising from such cellular dysfunction and *in vivo* electrophysiological investigations in preclinical models are generally restricted to a limited number of brain regions. Indeed, preclinical studies of schizophrenia models have independently displayed neural circuit alterations in prefrontal cortex^4,5^, hippocampus^4^, striatum^5^, or sensory thalamus^6^, but comparative brain-wide neural investigations with single-neuron resolution have not yet been performed. As a result, the exact topography of neural alterations throughout the brain in models for schizophrenia has remained elusive.

A major technical advance has been the development of high-density silicon probes capable of recording hundreds of neurons spread across the living brain simultaneously with single-neuron and action-potential spatiotemporal resolution.^7^ Such recordings have provided unique insight into brain-wide population encoding of movement, sensation, and reward in animal models^8–10^, and into multi-layer encoding of speech sounds in human cortex.^11^ Additionally, these methods could provide a powerful approach to comprehensively interrogate dysfunctional circuits *in vivo* in mutant rodent models for neurological or psychiatric disorders, such as schizophrenia.

A well-supported model for synaptic dysfunction in schizophrenia centers on n-methyl-d-aspartate (NMDA) receptor hypofunction. First, NMDA receptor antagonists mimic aspects of schizophrenia symptomatology, such as cognitive dysfunction^12^, and anti-NMDA receptor autoimmune encephalitis mimics several clinical symptoms of schizophrenia.^13^ Furthermore, common and rare high-risk genetic variants in *GRIN2A* encoding the NMDA receptor GluR2A subunit have been consistently associated with schizophrenia.^2,14,15^ Indeed, the *Grin2a*^+/-^ heterozygous loss-of-function mouse model provides a high validity preclinical model for schizophrenia in displaying striatal, hippocampal and dopaminergic transcriptional alterations.^16^ Thus, it poses an excellent model to interrogate how genetic synaptic dysfunction alters brain-distributed cellular and circuit physiology, resulting in schizophrenia susceptibility. Here, we perform comparative and comprehensive electrophysiological recordings using multiple simultaneous Neuropixels probes in adult, awake, head-fixed *Grin2a*^+/-^ heterozygous mutant mice and matched wildtype littermates. We report distributed and graded changes throughout the brain across various measures of neural activity, with prominent alterations in higher-order thalamus and connected prefrontal cortices. Acute restoration of higher-order thalamus activity in mutant mice is sufficient to restore brain-distributed electrophysiological phenotypes, suggesting a pivotal role for this structure in schizophrenia pathophysiology and its potential treatment.

## Results

### Comprehensive neural recordings in the *Grin2a*^+/-^ preclinical model for schizophrenia

We performed large-scale electrophysiological recordings in the *Grin2a*^*+/-*^ heterozygous mouse model using simultaneous Neuropixels probes in a head-fixed, acute setting (**Figs. S1-3; Materials and Methods**), targeted to brain regions including cortex, striatum, hippocampus and thalamus (**Fig. 1a-c**). We recorded adult (∼10-20 weeks), male, *Grin2a*^+/-^ mice and *Grin2a*^+/+^ wildtype littermate controls (*n* = 8/genotype) using ∼2-4 simultaneous probes (**Fig. 1d-g,j**). Care was taken to minimize experimental variation: Pairs of mutants and controls were handled, trained, and recorded at similar times of day with matched region targeting (**Table S1**). During training and recording, mice performed a self-paced rewarded sensorimotor task with both genotypes performing similar numbers of trials (**Fig. S1)**, but with minor differences between reward rate and movement kinematics (**Fig. S1**). Following spike sorting, recorded neurons with non-somatic waveforms were removed followed by additional quality control, with similar numbers of discarded neurons across genotypes, animals or brain areas (**Fig. S2**). Probe reconstruction and channel assignment were performed utilizing previously validated electrophysiological landmarks (**Fig. S3, Table S2**).

**FIGURE 1.**
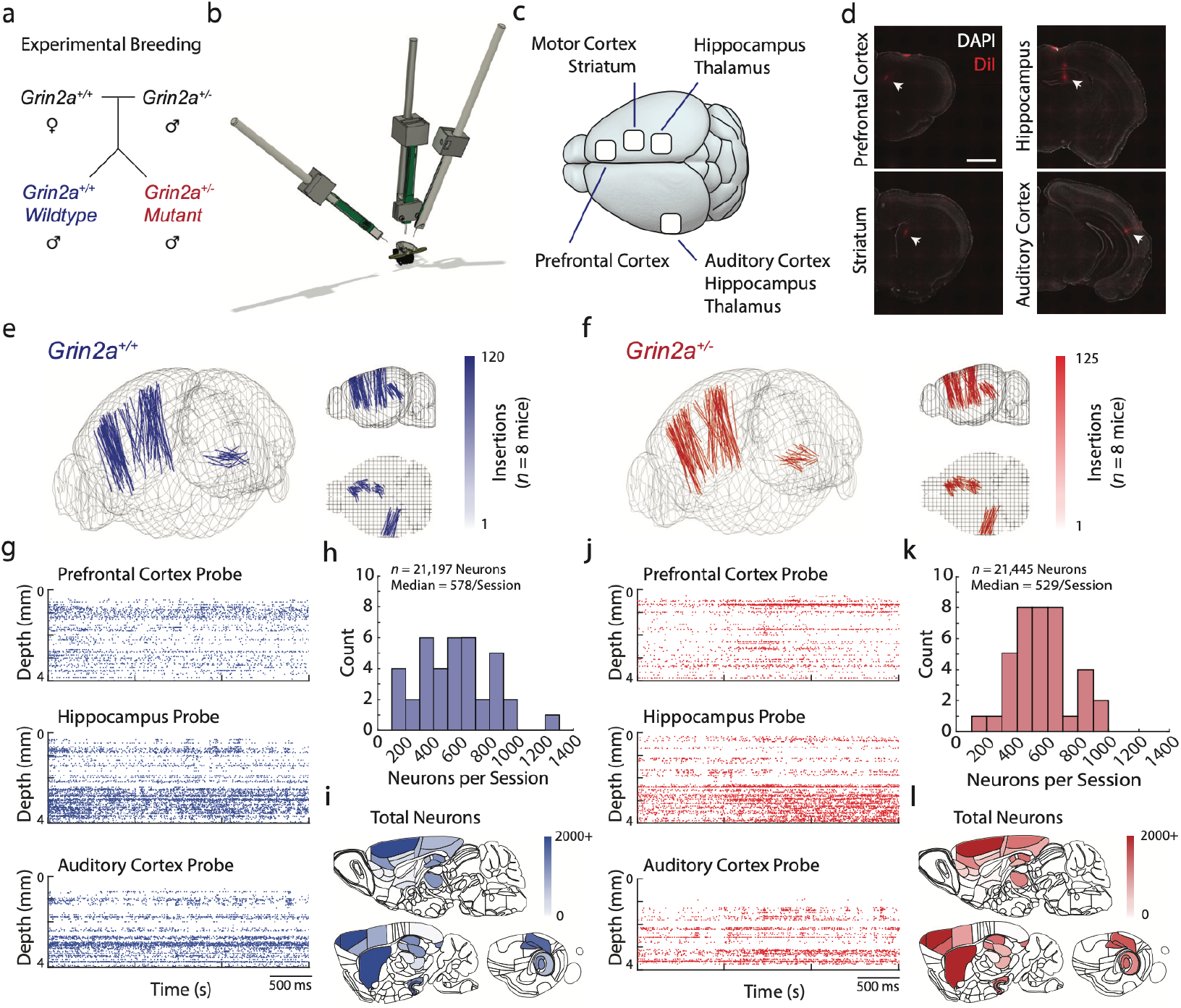
Comparative and comprehensive single-neuron electrophysiological recordings in the *Grin2a*^+/-^ mutant model for schizophrenia. **a)** Experimental breeding scheme. Wildtype *Grin2a*^+/+^ were crossed with mutant *Grin2a*^+/-^ to generate experimental cohorts of littermates consisting of male wildtype *Grin2a*^+/+^ and mutant *Grin2a*^+/-^ mice. **b)** Experimental schematic. Brain-distributed, high-density electrophysiological recordings in *Grin2a*^+/-^ mutant mice and *Grin2a*^+/+^ matched wildtype littermates using up to four simultaneous Neuropixels probes. **C)** Dorsal view of a mouse brain schematic. Craniotomy locations are targeted to prefrontal regions, motor cortices and striatum, hippocampus, auditory cortex, and thalamus. **d)**Exemplar coronal epifluorescent microscope images of probe tracks for prefrontal cortex, striatum, hippocampus, and auditory cortex using the fluorescent marker DiI (red, arrow). Nuclear marker DAPI in white. Additional examples in **Fig. S3, S7**. Scale bar, 1 mm. **e-f)** 3d schematic of brains showing reconstructed probe trajectories for *Grin2a*^+/+^ wildtype (e, blue) and *Grin2a*^+/-^ mutant mice (f, red). **g**,**j)** Example spike rasters across the depth of 3 individual probes from a single *Grin2a*^+/+^ (g, blue) and *Grin2a*^+/-^ mouse (j, red) for several seconds. Each dot represents a single spike. **h**,**k)** Histograms depicting the total number of neurons per recording session for *Grin2a*^+/+^ (h, blue) and *Grin2a*^+/-^ mice (k, red). **i**,**l)** Sagittal schematic view of mouse brain depicting the total number of neurons per brain region for *Grin2a*^+/+^ (i, blue) and *Grin2a*^+/-^ mice (j, red). *n* = 42,643 neurons/16 mice.

In total, we retained 21,445 neurons in mutant *Grin2a*^+/-^ mice (*n* = 125 insertions), and 21,197 neurons in wildtype *Grin2a*^+/+^ littermates (*n* = 120 insertions), with median 578 and 529 neurons per recording session, respectively (**Fig. 1h,k**). Total number of neurons, total number of probe channels, and yield (neurons per channel) were comparable across genotypes (**Fig. 1i,l**; **Table S1)**. In wildtype animals, electrophysiological characteristics of neurons in broadly defined areas (cortex, hippocampus, striatum, thalamus, other) showed expected differences between areas (**Fig. S4**), such as higher firing rate in thalamus and higher burst firing ratios in hippocampus^7,10,17^, indicating valid targeting to these areas.

Together, these data therefore provided a comprehensive, well-controlled cohort of high-density electrophysiological recordings in the mutant *Grin2a*^+/-^ genetic mouse model and matched littermate controls, without overt differences in behavior, recording locations, quality control, neuron number, or yield.

### *Grin2a*^*+/-*^ mutant mice display distributed and graded neural alterations with prominence of higher-order thalamus

We first examined electrophysiological alterations over a range of parameters across broadly defined areas between genotypes for the entire recording period (**Fig. 2a-c**). We found that thalamus, cortex, and hippocampus displayed reduced spontaneous firing rates in mutant *Grin2a*^+/-^ mice compared to controls, with thalamus showing most prominent rate reductions (-16.6%). Analogous effects were observed when exclusively analyzing intertrial recording epochs devoid of bar presses, stimuli, or rewards (**Fig. S5**), and were stable over time on a minute-to-minute timescale and across different task phases (**Fig. S5**). To ensure rate differences were not confounded by our pre-processing approach, we re-analyzed putative firing rate differences in the dataset preceding quality control (*n* = 62,642 neurons), which yielded similar effects (**Fig. S5**). In contrast to rate changes, single-channel waveform comparisons between genotypes showed minor effects (<10%) (**Fig. S6**), suggesting overt activity changes but minimal changes in waveform parameters in mutant *Grin2a*^+/-^ mice.

**FIGURE 2.**
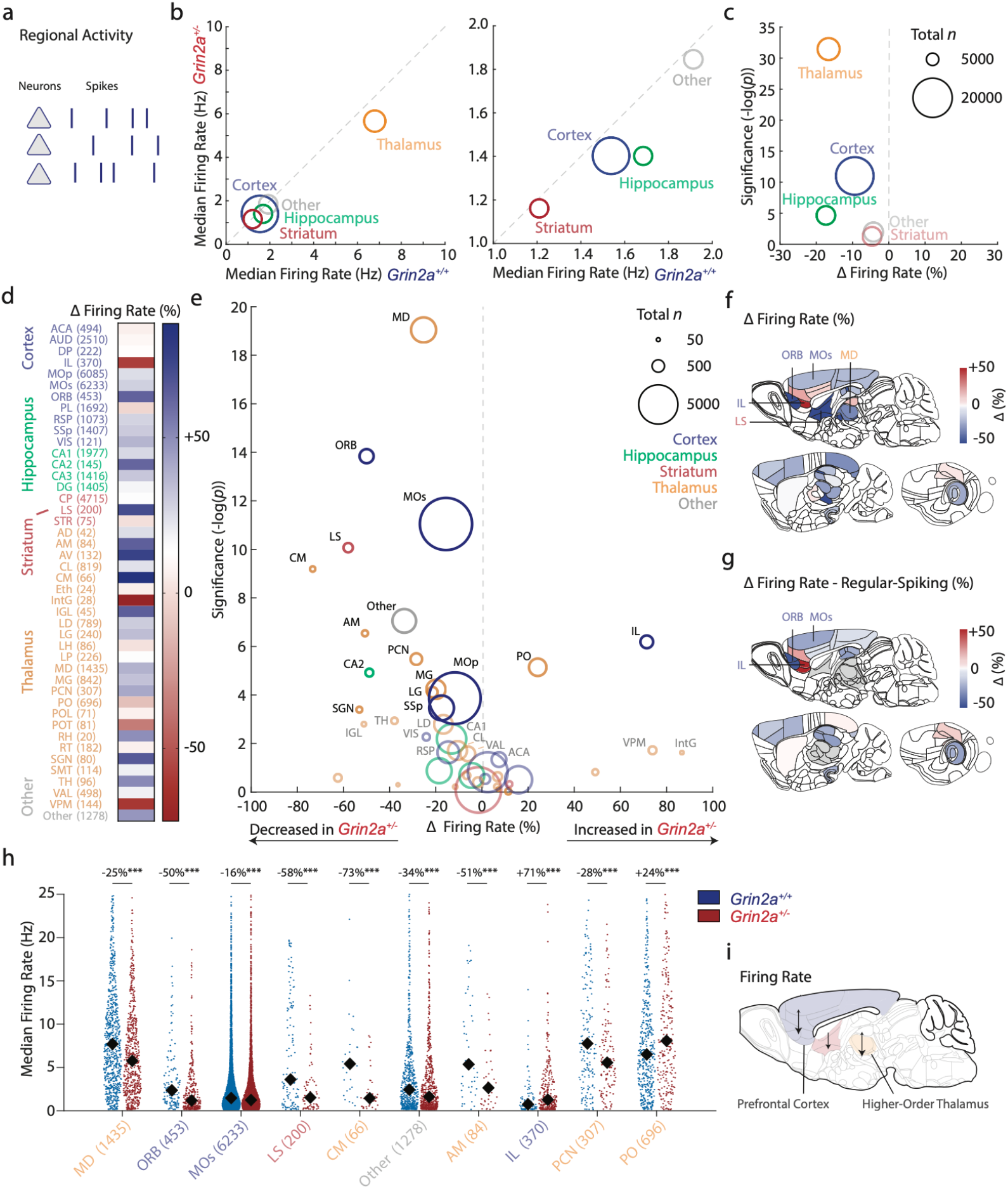
*Grin2a*^*+/-*^ mutant mice display distributed and graded alterations in neuronal activity. **a)** Schematic depiction of analysis of firing rate per region. **b)** Scatter plot depicting median firing rates for *Grin2a*^+/+^ and *Grin2a*^+/-^ mice per area. Circle sizes indicate total recorded neurons. Colors indicate grouping per area. **C)** Vulcano plot depicting effect size per region in spontaneous firing rate set out against significance of pair-wise genotype comparison per area. Circle sizes indicate total recorded neurons. Colors indicate grouping per higher-order area. *p* < 0.001, genotype*area two-way ANOVA. **d)** Heatmap showing effect sizes for firing rate between *Grin2a*^+/+^ and *Grin2a*^+/-^ mice per regions, grouped per larger area. **e)** Vulcano plot depicting effect size per region in spontaneous firing rate set out against significance. Circle sizes indicate total recorded neurons. Colors indicate grouping per higher-order area. Most prominent rate alterations occurred in higher-order thalamus (MD, CM, AM, PCN), and prefrontal cortices (ORB, MOs, IL). Greyed out regions indicate levels of corrected significance. *p* < 0.001, genotype*region two-way ANOVA. **f)** Sagittal brain schematics depicting genotype difference in firing rate per region, regardless of significance. Regions with highest pair-wise genotype significance indicated. **g)** Sagittal brain schematics depicting genotype difference in firing rate per region for regular-spiking neurons only, regardless of significance. Regions with highest pair-wise genotype significance indicated. **h)** Swarm plots with median firing rates depicting individual neurons for the regions attaining highest pair-wise genotype significance in (e). Black diamonds indicate medians, neuron numbers in brackets, and colors indicate grouping. Y-axes capped at 25 Hz. **i)** Summary graph depicting prominent rate differences in higher-order thalamus and mono-synaptically connected prefrontal regions in *Grin2a*^+/-^ mutant mice. \*\*\**p* < 0.001. Two-way ANOVA followed by *post hoc* non-parametric comparisons. Region abbreviations in **Table S3**. Neuron numbers in **Table S4**. Statistics in **Table S7**.

To further examine the robust activity changes, we next subdivided recordings of broad-defined areas into ∼45 individual brain regions (**Table S3** for detailed neuron number; **Fig. S7**), yielding a combined number of 4 to 3385 neurons per region per genotype (*Grin2a*^+/+^: mean ∼416 neurons; *Grin2a*^+/-^: mean ∼420 neurons). Here, genotype comparisons showed ∼35% of regions showed altered spontaneous firing rates compared to controls (*p* < 0.001, two-way ANOVA region*genotype), for which the majority showed activity decreases in mutant animals (∼86% of regions decreased; ∼14% increased; **Fig. 2d-f**)). Alterations were graded (genotype difference range -70% to +70%) and brain-distributed, but were most prominent in thalamus and cortex: Regions with the most prominent activity decreases included the mediodorsal nucleus of the thalamus (MD), orbitofrontal cortex (ORB) and secondary motor cortex (MOs), while infralimbic area of the prefrontal cortex (IL) and posterior nucleus of the thalamus (PO; **Fig. 2f,h,i**) showed rate increases. Combined, thalamic regions with prominent rate alterations consisted of higher-order (prefrontal, associative) thalamic nuclei (**Fig. S8)**, featured a multi-areal matrix projection pattern^18^ (**Fig. S8)**, and were reported to display monosynaptic connections to most prominently affected prefrontal cortices (**Fig. S8**), indicating overarching effects on a connected thalamocortical circuit. Furthermore, medial prefrontal cortical regions showed a bidirectional gradient in rate alterations along its anatomical dorsoventral axis (**Fig. S9**), suggesting a structure to activity alterations across the medial prefrontal cortex as a whole. Analogous subdivision of hippocampus into dorsal and ventral hippocampus^19^ revealed an additional dorsoventral axis of rate effects between genotypes, with dorsal hippocampus displaying hyperactivity and ventral hippocampus hypoactivity in mutants, driven predominantly by changes in CA3 firing rates (**Fig. S10**). Firing rate decreases across the brain were accompanied by modest increases in firing rate variability (CV, CV2), predominantly in thalamus and cortical regions, respectively (**Fig. S11**).

Together, these data indicate *Grin2a*^+/-^ mutants display a distributed, graded pattern of electrophysiological alterations across many regions with most prominent changes in higher-order thalamic nuclei and synaptically connected prefrontal cortices.

### *Grin2a*^*+/-*^ mutant mice display distributed and graded burst firing ratio increases across cortical regions

Activity of individual neurons can occur in bursts – quick successions of action potentials – commonly dependent on intact NMDA receptors.^20^ We next examined whether activity reductions in mutant *Grin2a*^+/-^ mice were related to alterations in neuronal burst firing, defined as spikes with low (<6 ms) inter-spike intervals (**Materials and Methods**). Spikes in bursts often occurred as doublets (∼77%), or triplets, a pattern globally consistent across genotypes (**Fig. S12**).

Regional analyses restricted to single spikes without bursting revealed similar distributed and graded effects as all spikes combined (*p* < 0.001, two-way ANOVA region*genotype; **Fig. S12)**, suggesting minimal contribution of burst spikes to overall firing rate reductions. However, burst firing frequency or burst firing ratio, defined as the number of bursts to the total number of spikes, showed a different pattern: Here, ∼34% of regions showed significant burst firing ratio changes, and this predominantly featured *increases* (∼65% of regions) in burst firing ratios, occurring most prominently in prefrontal and midline cortical regions (RSP, ACA, MOs; **Fig. S12**). A partial and inverse overlap between firing rate alterations in mutant mice and burst firing ratio alterations in mutant mice was observed at the regional level (**Fig. S13**), suggesting partially independent mechanisms for rate and burst ratio alterations at the level of individual regions.

Together, these data suggest firing rate alterations were observed across burst and single spiking populations in *Grin2a*^*+/-*^ mice compared to wildtype littermates, but indicate distributed and graded alterations in neuronal burst firing ratios. Regions with burst ratio alterations partially overlapped with regions containing firing rate alterations and were most prominently observed in prefrontal and midline cortical regions.

### *Grin2a*^*+/-*^ mutant mice display balanced alterations across regular-spiking and fast-spiking neurons

GABAergic inhibitory interneuron dysfunction is a proposed key cellular pathophysiological mechanism for schizophrenia, particularly for fast-spiking, parvalbumin-positive interneurons^21^, and NMDA hypofunction in fast-spiking neurons is a widely supported cellular model for the disease.^22^ Therefore, we next investigated whether rate alterations across cortex, hippocampus, and striatum in *Grin2a*^+/-^ animals were driven by dysfunction of the subpopulation of fast-spiking interneurons.

We differentiated cortical, hippocampal, and striatal neurons into two populations based on repolarization latency of the averaged single-channel waveform (trough-to-peak latency; **Figs. S14-16**), which in wildtype mice differed from one another across other waveform and firing metrics (**Figs. S14**). This classification yielded proportions of regular-spiking and fast-spiking neurons that were highly comparable in both genotypes across grouped regions (**Fig. S15**), suggesting no global reductions of fast-spiking neuron numbers in mutants. Comparing genotypes, we observed neural activity and waveform alterations between *Grin2a*^+/-^ mice and controls across both regular-spiking and fast-spiking neurons over grouped areas (**Fig. S16**): Firing rates were altered in both classes across areas, with concomitant alterations in rate pattern and variation, as well as minor alterations in waveform metrics (**Fig. S16**). Region-specific analyses demonstrated that firing rate alterations for regular-spiking neurons were highly similar to those observed when classes were combined (**Fig. 2**; **Fig. S15**), with greatest differences between genotypes in prefrontal cortical regions (MOs, ORB, IL). Similarly, mutant *Grin2a*^+/-^ mice displayed distributed alterations in fast-spiking neuronal firing across individual regions (**Fig. S16**) and included prefrontal cortical regions (ORB, MOs), suggesting overlapping effects across both subclasses, predominantly in prefrontal cortex. Bursting alterations were similarly observed across both regular-spiking and fast-spiking neurons (**Fig. S13**). Thus, alterations across various metrics were observed across both regular-spiking and fast-spiking neuron populations.

As observed for neuronal classes combined, regular-spiking and fast-spiking neurons in medial prefrontal cortex regions each showed a bidirectional gradient in activity alterations along the dorsoventral axis (**Fig. S9**), inverse to one another: Regions showing reduced regular-spiking activity in *Grin2a*^*+/-*^ mice compared to controls showed increased fast-spiking neuron activity, and vice versa. A broader direct comparison of rate effect sizes of both classes revealed that such an inverse correlation extended throughout the brain (**Fig. S16**). These findings suggest that instead of uniform alterations of fast-spiking neuron functioning across the brain, regular-spiking and fast-spiking neurons in mutant mice present alterations across both populations and a balanced gradient of rate alterations across regions.

Together, in contrast to previous work, we did not find evidence for uniform reductions in fast-spiking neuron activity across the brain nor for alterations exclusive to fast-spiking neurons. Instead, these data indicate that cross-parameter neural alterations in *Grin2a*^*+/-*^ mice were class-independent and occurred in balanced, non-uniform fashion across both regular-spiking and fast-spiking putative interneurons.

### *Grin2a*^*+/-*^ mutant mice display widespread co-activity increases with prominence of prefrontal cortical regions

How do distributed and graded region- and cell class-independent neural alterations correspond to circuit dysfunction? Burst firing and interneuron functioning are intricately linked to moment-to-moment local network activity^23,24^, and thus neural alterations could lead to altered coordination of such neuronal co-activation in mutant animals.

To examine whether mutant *Grin2a*^+/-^ mice displayed neural co-activity alterations, we next calculated short-term (25-ms) normalized rate correlations between pairs of neurons within individual brain regions across genotypes (**Fig. 3a-c**). This analysis revealed varying non-zero mean correlations per region across genotypes (**Fig. S17**) that disappeared under conditions of random or circular shuffling of individual neuron spiking time bins (**Fig. S17**). Mean correlation values across the brain showed a rostrocaudal gradient in both genotypes, with prefrontal regions showing overall comparatively lower mean neural co-activity values (**Fig. S17**).

**FIGURE 3.**
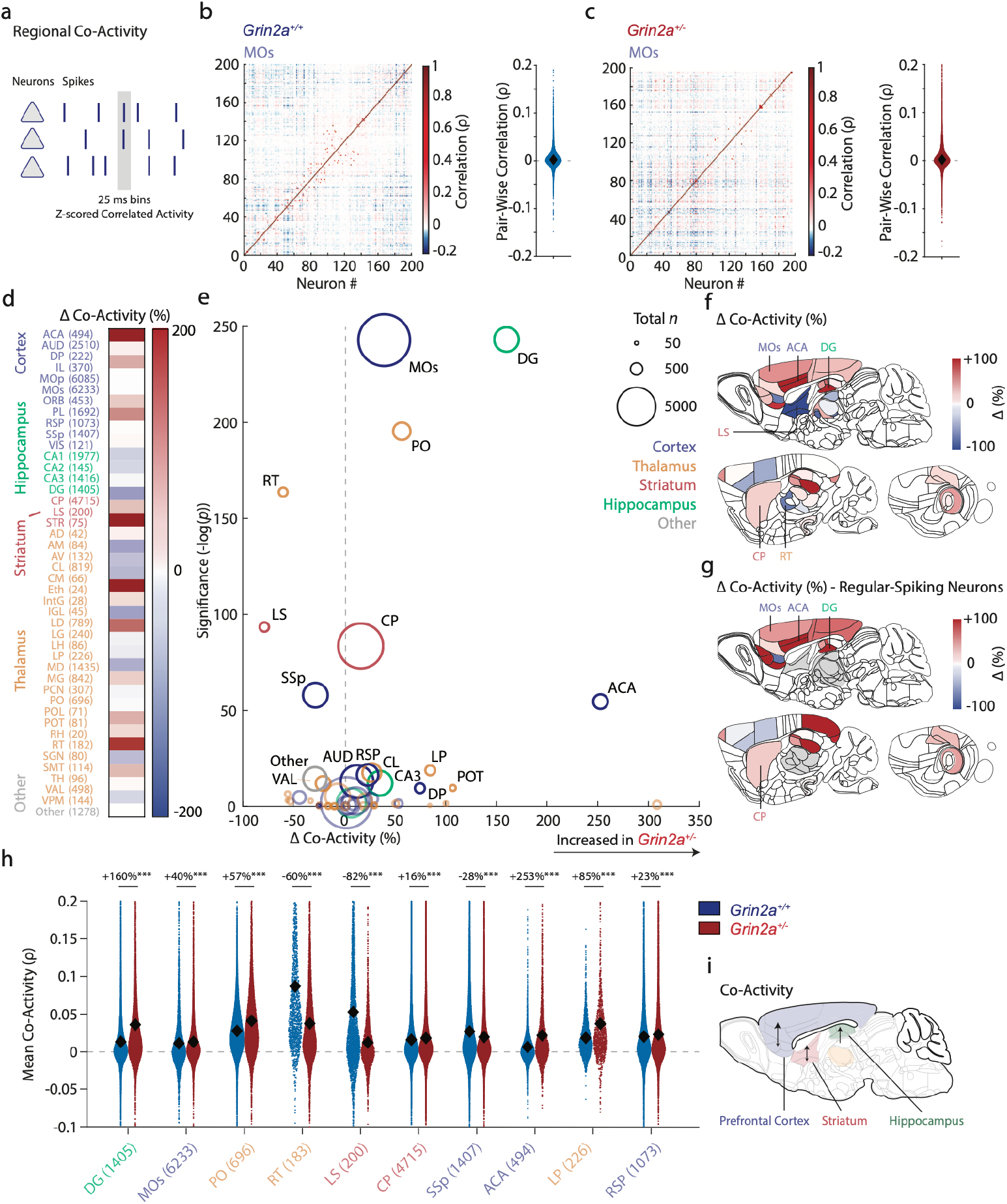
*Grin2a*^*+/-*^ mutant mice display graded and distributed increased neuronal co-activity. **a)** Schematic depiction of analysis. Pair-wise neuronal activity per region was analysed over 25-ms time windows in both genotypes. **b-c)** *Left*: Within-region connectivity for an exemplar session from secondary motor cortex (MOs) calculated through pair-wise correlations for a single recording for a wildtype *Grin2a*^*+/+*^ mouse (b) and a littermate *Grin2a*^*+/-*^ mouse (c). *Right*: Correlation values (Spearman’s *ρ*) for all pairs in the same session. Each dot represents a pair-wise Spearman comparison of a single recording session of neurons within secondary motor cortex (MOs). **d)** Heatmap depicting region-specific co-activity alterations between genotypes, grouped and color-coded per higher order area. **e)** Vulcano plot depicting co-activity effect size between genotypes set out against regional pair-wise significance for regular-spiking neurons and fast-spiking neurons combined. *Grin2a*^+/-^ show increased co-activity across a large number of regions. Circle sizes indicate total neuron numbers, and colours indicate area grouping. Greyed out regions indicate levels of corrected significance. *p* < 0.001, genotype*region two-way ANOVA. **f)** Sagittal schematic view depicting regional alterations regardless of significance, for regular-spiking neurons and fast-spiking neurons combined. Regions with highest pair-wise genotype significance indicated. **g)** Sagittal schematic view depicting regional alterations regardless of significance, restricted to regular-spiking neurons. Regions with highest pair-wise genotype significance indicated. **h)**Swarm plots with individual neuron for the regions attaining highest genotype significance in (e) for co-activity. Black diamonds indicate means, neuron numbers in brackets, and colors indicate grouping. Y-axis capped at -0.1 and 0.2 for visibility. **i)**Summary graph depicting prominent network co-activity increases in striatum, prefrontal cortical regions, and hippocampal DG and CA3 in *Grin2a*^+/-^ mutant mice. Region abbreviations in **Table S3**. Neuron numbers in **Table S4**. Statistics in **Table S7**. \*\*\**p* < 0.001. n.s. non-significant. Two-way ANOVA followed by *post hoc* non-parametric comparisons.

Genotype comparisons per region for all neurons combined revealed brain-distributed alterations in local neuronal co-activity between genotypes (**Fig. 3d,e**; *p* < 0.001, two-way ANOVA region*genotype). Pair-wise regional comparisons revealed ∼71% of regions showed significant alterations in co-activity, with most regions displaying co-activity increases (∼70% increased). Higher neural co-activity was particularly notable in prefrontal cortical regions (ACA, MOs; **Fig. 3d-f,h,i**), striatum (CP), and hippocampus (DG, CA3), and prominent lower co-activity was observed in lateral septum (LS) and thalamic reticular nucleus (RT). Analogous to firing rate, we observed a bidirectional pattern of effects in co-activity between genotypes across dorsoventral axis of the medial prefrontal cortex (**Fig. S9**).

Co-activity alterations between genotypes were robustly observed across a broad range of time windows (10-500 ms), suggesting invariance to time interval (**Fig. S17**). We analogously observed alterations in co-activity analysis in cortex, hippocampus and striatum when including both regular-spiking and fast-spiking neurons (**Fig. 3d-f**) or restricted to regular-spiking (**Fig. 3g**; **Fig. S17-S18**) or fast-spiking neurons (**Fig. S17-S18**), suggesting that neural coordination alterations were independently observed across both neuron classes. Co-activity across thalamic nuclei showed a bidirectional pattern of changes between genotypes (**Fig. S19**). Overall, regions that showed co-activity alterations only partially overlapped with those that displayed firing rate or bursting alterations, suggesting partially independent mechanisms at regional level (**Fig. S19**).

Together, these data suggest distributed and graded alterations in spontaneous local network co-activity in *Grin2a*^*+/-*^ mice across regions and spiking subclasses, with prominent co-activity increases in prefrontal cortical regions, striatum, and hippocampus.

### Absence of overt relation to wildtype Grin2a gene expression

We hypothesized that cross-parameter alterations could potentially correlate to region-by-region Grin2a gene expression in wildtype mice. However, spontaneous rate activity, burst firing, or co-activity effect sizes between genotypes across broad-defined regions compared to data obtained from three transcriptomic databases did not reveal a consistent relation (**Fig. S20**), especially not for thalamus. This suggests neural alteration metrics between genotypes did not overtly correlate to wildtype Grin2a gene expression.

### Restoration of higher-order thalamus activity is sufficient to correct brain-distributed cross-parameter alterations

The above findings demonstrate prominent activity reductions in higher-order thalamus and cross-parameter alterations across prefrontal cortex, regions with extensive reciprocal connections (**Fig. S8**)^18,25^. Furthermore, indicative of this connectivity, previous work in wildtype animals suggests that higher-order thalamus could present a causal modulator of prefrontal cortical activity, network organization and burst firing.^26,27^ Thus, we hypothesized that reestablishing firing rate in higher-order thalamic nuclei in *Grin2a*^+/-^ mutant animals to wildtype levels would causally restore normative cross-parameter neural changes across connected prefrontal cortices.

To test this hypothesis, we performed stereotactic bilateral adeno-associated viral (AAV) injections of activating DREADDs (hM3Dq-mCherry) or control fluorescent protein (mCherry) in adult *Grin2a*^+/-^ mutants (*n* = 3-4/intervention), in affected higher-order thalamic nuclei (**Fig. 4a**; **Fig. S21**; **Materials and Methods**). Four weeks later, animals were injected with clozapine-n-oxide (CNO, 1 mg/kg i.p.) ∼45 mins before initiating large-scale recordings (**Fig. 4b-d**). Following quality control and region assignment (**Fig. S21-S22)**, we retained 11,616 neurons in *Grin2a*^+/-^ mice with hM3Dq-mCherry (*n* = 80 insertions) and 9,521 neurons in *Grin2a*^+/-^ mice with mCherry (*n* = 77 insertions). Targeted recordings in higher-order thalamus showed potent enhancement of firing rate under CNO application in hM3Dq-mCherry mice (+50%) compared to untreated mutants, normalizing to wildtype levels, but not in mCherry controls with CNO (**Fig. 4d**). Higher-order thalamic normalized activity was stable across the full recording window and CNO application in mCherry controls had minimal effect (**Fig. S21**). No effects of acute CNO application were observed on animal performance on the sensorimotor task (**Table S6**). Thus, acute chemogenetic activation in adult mutant mice restored higher-order thalamic nuclei firing rates back to wildtype levels.

**FIGURE 4.**
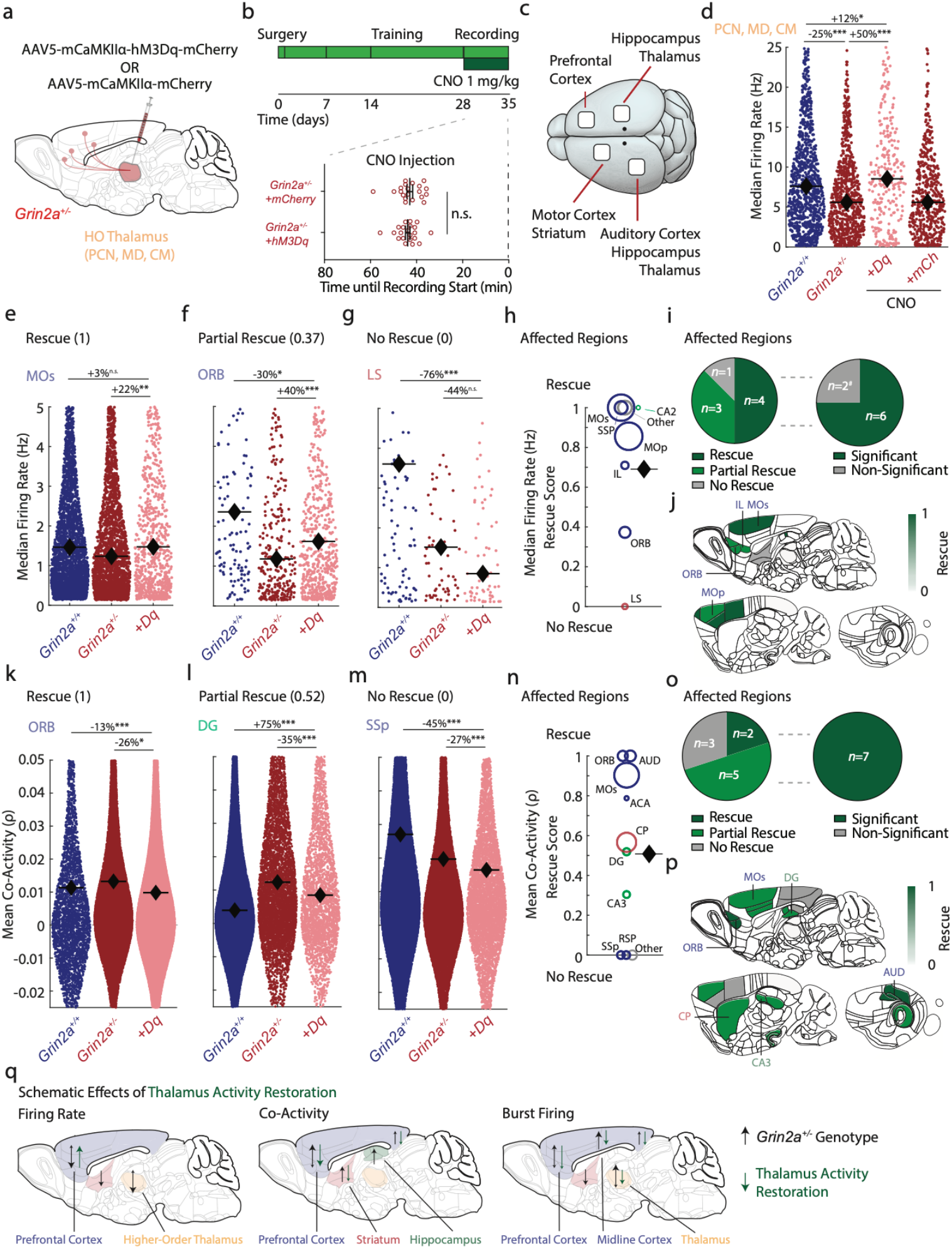
Acute restoration of higher-order thalamus activity corrects distributed neural alterations in *Grin2a*^*+/-*^ mutant mice. **a)** Schematic sagittal view of manipulation approach. Mutant *Grin2a*^+/-^ mice received stereotactic bilateral injection of AAV5 containing hM3Dq-mCherry or mCherry control in higher-order thalamus targeting MD, CM, and PCN, followed by large-scale recordings. **b)** *Top*: Injection schematic. After training and task learning, mice received i.p. CNO 1 mg/kg immediately before head-fixation and subsequent recording. *Bottom*: Time from injection until recording start. No difference in time from CNO injection to recording start between hM3Dq-mCherry or mCherry control mice. **C)** Top schematic view of craniotomy and recording locations. **d)** Median firing rate for *Grin2a*^+/+^ (blue), *Grin2a*^+/-^ (dark red), *Grin2a*^+/-^ hM3Dq-mCherry (light red), and *Grin2a*^+/-^ mCherry (light red) for higher-order thalamus (MD, CM, PCN) showing normalized firing rates under CNO application, but not in mCherry controls. Dots indicate individual neurons, black diamonds indicate median values. Y-axis capped at 25 Hz for visibility. **e**,**f**,**g)** Median firing rate for *Grin2a*^+/+^ (blue), *Grin2a*^+/-^ (dark red), and *Grin2a*^+/-^ hM3Dq-mCherry under CNO (light red) for three regions with different rescue scores: MOs (e), ORB (f), LS (g). Each dot represents an individual neuron, black diamonds indicate medians. Y-axis capped at 5 Hz for visibility. **h)** Scatter plot depicting rescue scores across the 8 significantly affected non-thalamic regions. Each circle indicates a region, circle size indicates neuron number. The majority of regions show full rescue of spontaneous firing rates. Black diamond indicates the mean rescue score. **i)** Pie chart showing the fraction of regions showing no, partial, or full rescue (left), along with the fraction of partial and fully rescue regions attaining single-region significance for firing rates (right). ^#^MOp attained marginal significance for manipulation comparison of firing rate (*p* = 0.07). **J)** Schematic sagittal view depicting rescue scores for firing rates across affected regions. **k**,**l**,**m**) Mean neuronal co-activity values for *Grin2a*^+/+^ (blue), *Grin2a*^+/-^ (dark red), and *Grin2a*^+/-^ hM3Dq-mCherry under CNO (light red) for three regions with different rescue scores: ORB (k), DG (l), SSp (m). Each dot represents an individual neuronal pair, black diamonds indicate means. Y-axis capped at -0.025 and 0.05 π for visibility. **n)** As (h), but for the 10 significantly affected non-thalamic regions for neural co-activity. Black diamonds indicate the median rescue scores. **o)** Pie chart showing the fraction of regions showing no, partial, or full rescue (left), along with the fraction of partial and fully rescue regions attaining single-region significance for co-activity (right). **p)** Schematic sagittal view depicting rescue scores for network co-activity across affected regions. **q)**Schematic depiction of overall findings of the current study. *Grin2a*^+/-^ mutant mice show widespread alterations across a range of metrics (black arrows), most pronounced in higher-order thalamus, and targeted thalamus enhancement instantiates widespread circuit corrections back to wildtype levels (green arrows). Region abbreviations in **Table S3**. Neuron numbers in **Table S4** and **Table S6**. Statistics in **Table S7**.\*\*\**p* < 0.001, \*\**p* < 0.01, \**p* < 0.05. n.s. non-significant. Unpaired two-tailed student’s *t*-test in (b). One-way ANOVA followed by *post hoc* comparisons in (d-g) and (k-m).

Targeted restoration of higher-order thalamus activity in mutant mice altered spontaneous firing rates across brain regions compared to untreated mutant mice (*p* < 0.001; two-way ANOVA). To examine effects of thalamic restoration on brain-distributed neural parameters, we computed a ‘rescue score’ (**Materials and Methods**) to standardize regional effects in treated mutants on a 0-1 scale between untreated mutants and untreated wildtype animals: A score of 0 indicated no effect or exacerbation and 1 indicating complete rescue or normalization beyond wildtype levels (**Fig. 4e-g**). For spontaneous firing rate, of eight non-thalamic regions altered in the discovery cohort, five regions attained correction back to wildtype rates (∼62.5%), two showed a partial rescue (∼25%), and one region showed no effect (∼12.5%). Of seven regions that showed a full or partial rescue (rescue score >0), four regions attained significant increased or decreased rate differences from untreated controls, with a mean rescue score of 0.70, indicating an average ∼70% correction toward wildtype levels (**Fig. 4h-j**). Normalization of spontaneous firing rates in cortex were observed across both regular-spiking and fast-spiking neurons (**Fig. S23**).

Targeted restoration of higher-order thalamus activity in mutant mice further altered neural co-activity across brain regions compared to control mice (*p* < 0.001; two-way ANOVA). Here, of ten non-thalamic regions altered in the discovery cohort, two regions attained normalization to wildtype levels (∼20%), five showed partial rescue (∼50%), and three regions showed no effect (∼30%; **Fig. 4k-m**). Of seven regions that showed a full or partial rescue, seven attained significant increases or decreases in co-activity from control levels (100%), with a mean rescue score of 0.50 (∼50% correction to wildtype levels; **Fig. 4n-p**). For burst firing ratios, of eleven non-thalamic regions from the discovery cohort, six regions attained partial or full rescue (∼55%), and four regions showed no effect (∼30%; one region was not recorded). Of six regions that showed a rescue, three attained significant increases or decreases in differences in burst firing from controls (∼50%; **Fig. S25**), with a mean rescue score of 0.40 (∼40% correction to wildtype levels). Rate and burst alterations showed minor differential trends across dorsal and ventral hippocampus (**Fig. S24**). Thus, restoration of higher-order thalamus activity partially or fully rescued cross-parameter neural abnormalities distributed across affected non-thalamic regions in *Grin2a*^+/-^ animals (**Fig. 4q**).

While we hypothesized correction of prefrontal regions, regions with partial or full rescue unexpectedly included non-prefrontal regions such as striatum, dorsal hippocampus and primary motor and sensory regions (**Fig. 4h,n**), suggestive of widespread correction from acute thalamic activity enhancement alone. Striatum receives collateral inputs directly from higher-order thalamus^28,29^ (**Fig. S8**), and thus monosynaptic inputs could be hypothesized sufficient to drive restorative striatal effects. However, we note that to date no monosynaptic connections have been reported from targeted higher-order thalamus to hippocampus or primary sensory cortices^18,25^, supported by absence of mCherry+ axons in these regions in our experiments (**Fig. S26**), suggestive of a trans-synaptic origin of restorative effects. Furthermore, correcting effects across regions were bidirectional, with metrics showing increases and others decreases, reversed towards wildtype levels. Thus, unidirectional activity enhancement of higher-order thalamus activity led to bidirectional corrective effects distributed throughout the brain.

Together, these findings suggest that acute restoration of higher-order thalamus activity in adult *Grin2a*^+/-^ mice was sufficient to correct cross-parameter, bidirectional neural alterations across prefrontal cortices, and propagating trans-synaptically to distal regions, instantiating distributed neural restorations.

## Discussion

A major limitation to circuit dissection in psychiatric illness such as schizophrenia has been the lack of spatiotemporal resolution for single-neuron and single-action potential interrogation in the living brain. As a result, despite convergent genetics suggesting neuronal synaptic dysfunction as cellular pathophysiological mechanism^2,3^, the brain-wide topography of this dysfunction in schizophrenia has remained uncertain. Here, we employ comprehensive high-density recordings *in vivo* to reveal graded and distributed neural alterations across the brain arising from a genetic NMDA receptor hypofunction in *Grin2a*^+/-^ mutant mice. Notably, higher-order thalamus and connected prefrontal cortices presented markedly altered firing rates in mutant animals, and partially overlapping regions displayed large alterations in burst firing ratios and co-activity. Restoring activity of higher-order thalamic neurons of mutant animals to the level of wild-type animals alone was sufficient to bidirectionally restore normative functioning in synaptically connected frontal cortices. Remarkably, this intervention also restored normative activity, bursting, and coactivity in several areas not known to be monosynaptically coupled to higher-order thalamus. Thus, these findings indicate that global neural alterations caused by genetic synaptic dysfunction can be reversed by restoring normative activity at a single locus. Together, these data suggest that higher-order thalamus is a pivotal region in schizophrenia pathophysiology and may provide a potent site of intervention for circuit-informed therapeutic strategies.

Detailed analysis of our multi-region recordings demonstrated that *Grin2a*^+/-^ mutants have widespread and partially overlapping alterations of rate, firing pattern, and burst firing across regions. This, analogous to transcriptomic findings^16^, indicates that genetic NMDA receptor hypofunction leads to broad neural alterations beyond individual regions, with adjacent regions showing a multitude of effects (e.g. gradients, bidirectionality). Further complexity resided in the observation that some individual regions displayed genotype differences on one parameter (e.g. ACA: burst firing) but not on another (e.g. ACA: firing rate). Such complex region-superseding variation urges caution in conclusions based on studies of single regions and encourages within-animal multi-region recording in models for schizophrenia or other pathologies.

Prominent alterations were observed in interconnected higher-order thalamic nuclei and prefrontal cortices. While prefrontal cortices have been robustly linked to schizophrenia^30^, higher-order thalamic nuclei have comparatively remained less examined. However, a growing number of findings have started to implicate higher-order thalamus in schizophrenia pathophysiology. In addition to prefrontal cortices, higher-order thalamus is widely connected to striatum, amygdala, and hippocampus through entorhinal cortex^18,28^, all implicated in domains of schizophrenia pathology. Indeed, higher-order thalamus dysfunction has been implicated in patients through human imaging^31^ and lesion^32^ studies, and targeted modulation of thalamus in wildtype animals recapitulates schizophrenia-linked symptomatology such as cognitive dysfunction.^33^ In addition, the mediodorsal thalamus in particular has been implicated in supporting social interaction,^34^ and routing of corollary discharge signals^35^, further domains thought affected in patients. Thus, higher-order thalamus resides at a crucial nexus between subcortical and prefrontal regions and its dysfunction could play a particularly crucial role in disease pathogenesis. Whether higher-order thalamic dysfunction generalizes across other models for schizophrenia beyond *Grin2a* and whether thalamic alterations occur primary (upstream) or secondary (downstream) in the pathophysiological hierarchy remains to be elucidated.

Changes in neural activity metrics across the brain appeared minimally correlated to bulk Grin2a expression levels as reported by various transcriptomic atlases. Mammalian expression patterns for genes implicated in brain disorders are often employed to guide explorations in molecular, cellular, or circuit endeavours^36^, with regions presenting high levels of a particular gene or protein assumed to suffer from greater impact of given mutations. The present expression-effect misalignment for *Grin2a* suggests a reassessment of this assumption and indicates alternative factors could modulate topographical circuit impact of genetic variation. For example, the ∼6-fold higher firing rate in thalamus than cortex^7,10,17^ could render it particularly vulnerable to genetic mutations in synaptic components, more so than baseline molecular expression. Nonetheless, these findings further underscore the importance of unbiased, brain-wide electrophysiological investigation in pathological models.

For cortical and hippocampal dysfunction, GABAergic interneuron alterations have been broadly supported as a key cellular dysfunction in schizophrenia.^21^ However, we did not observe uniform widespread activity reductions in fast-spiking putative interneurons in *Grin2a*^*+/-*^ mice. Instead, we report a non-uniform pattern of neural differences in both regular-spiking and fast-spiking classes, which overall presented comparable alterations across regions: Alterations in activity, bursting, or waveform changes were observed across both classes. While the possibility remains that fast-spiking effects could be masked by compensatory changes, these data suggest that either fast-spiking neuron dysfunction is not affected uniformly throughout the brain, or dysfunction resides not in neural activity but perhaps in aberrant integration of afferent inputs or efferent transmission to postsynaptic targets. Here, too, we argue that unbiased examination of fast-spiking neuron alterations across the brain in models of disorders are paramount.

Neural co-activity indicates the structured coordination of spiking of neurons, and is thought to mediate communication within and across brain regions.^37^ We report widespread increases in regional neuronal co-activity in *Grin2a*^*+/-*^ mice, most prominently observed across prefrontal corticex, and appear independent of time window length, task phase, bursting, and firing rate. These effects are in line with studies showing increased neural co-activity in medial prefrontal cortex^38^ and visual cortex^39^ in other preclinical models of schizophrenia, and align with the suggestion that antipsychotic medications could function through decorrelating (ie. reducing co-activity of) longer-range connections.^40^ Partially overlapping with rate and bursting, co-activity alterations were observed in prefrontal regions synaptically connected to altered higher-order thalamus, further supporting cross-parameter implication of the prefrontal-thalamocortical circuit. Increases in neural co-activity across prefrontal cortical regions has been hypothesized to present the circuit alteration to underlie predictive coding abnormalities in schizophrenia.^41^ Following from this, as enhancement of higher-order thalamus activity sufficed to normalize cortical co-activity, a subsequent hypothesis arises that primary higher-order thalamus dysfunction could be hypothesized to causally underlie predictive coding abnormalities in schizophrenia. Regardless, exactly how widespread co-activity changes result in the broad diversity of schizophrenia symptoms remains to be elucidated.

The recurrent alterations across a prefrontal thalamocortical circuit present higher-order thalamus as a pivotal target for intervention. Indeed, restoration of thalamic activity normalized cross-parameter alterations in prefrontal regions in *Grin2a*^*+/-*^ mice, in line with similar targeted modulations in wildtype.^26^ However, the present corrections included restoration of multiple prefrontal cortical regions, reflecting the strongly divergent contacts of higher-order thalamocortical projections.^42^ Unexpectedly, restorative effects of normalizing thalamic firing rate to wild-type levels propagated into primary motor and sensory cortices, as well as hippocampus, regions generally not reported to receive monosynaptic connections from targeted higher-order thalamic nuclei. Notably, restorative experiments here were performed in post-developmental animals (age ∼12-20 weeks), using acute (CNO injections before recording), and thus challenge hypotheses^4^ that physiological rescue is restricted to a particular critical period. The findings align with others^43,44^ suggesting that post-developmental, adult reversal of pathophysiology and symptoms for schizophrenia is plausible, which is particularly relevant for clinical translation. The small size of higher-order thalamic

nuclei, homogeneity of neuron type, lack of recurrent connections, and divergent cortical connectivity pattern position them as an attractive therapeutic for schizophrenia, for example through transcranial ultrasound approaches or deep-brain stimulation.^45^ While the exact circuit mechanisms of how thalamus restoration results in widespread corrections remain to be elucidated, it could route in part via enhancement of cortical fast-spiking neurons, which here showed uniform rate enhancement across cortical regions after manipulation (∼30% increase overall).

In general, the development of treatments for psychiatric illnesses is impaired by the lack of identified core pathological circuits. Patient neuroimaging contains limited spatiotemporal resolution and encephalography lacks depth resolution. Human transcriptomic approaches on post-mortem tissue are generally restricted to a small number of brain regions, and human induced-pluripotent stem cell-derived approaches are often similarly restricted to specific glutamatergic, GABAergic, or dopaminergic neurons. While these approaches valuably contribute to biological understanding of disease and could assist therapeutic discovery, they remain limited for dissection of complex circuits. The current findings argue that next-generation electrophysiological models could provide a missing investigational link: Identification of a restricted number of pivotal circuits from such experiments could provide much-needed focus for subsequent targeted imaging, transcriptomic, and cell-culture approaches.

Together, the current findings present the first comprehensive *in vivo* electrophysiological examination with single-neuron precision of a high-validity model for schizophrenia. These findings indicate that detailed, unbiased electrophysiological preclinical investigations can identify novel pivotal pathophysiological mechanisms with strong potential for clinical translation.

## Author contributions

Conceptualization: JS

Methodology: JS, KP, YP

Investigation: JS, KP, YP

Visualization: JS, KP

Funding acquisition: JS, CS, AS

Supervision: CS, AS

Writing – original draft: JS

Writing – review & editing: JS, KP, YP, CS, AS

## Acknowledgements

This research was supported by the Wellcome Trust (224129/Z/21/Z), John Fell Fund Oxford (0013599), and Win Seed Grant to JS, a Medical Research Council PhD Scholarship (MR/W006731/1) and Onassis Foundation Scholarship (F ZU 056-1/ 2024-2025) to KP, a Wellcome Trust award to CJS (224430/Z/21/Z), as well as a Medical Research Council UK award to AS (MC_UU_00003/6). The authors would like to thank Jane Westcott and Rae Dolman for animal technical assistance and Ben Micklem for technical support.

## Conflicts of interest

The authors declare no competing financial interests associated with this study.

## Data and materials availability

All data supporting the figures are available in the manuscript or supplementary materials. The *Grin2a* mouse line was obtained from Riken BRC (#RBRC02256) deposited by Dr. Motoya Katsuki, under a material transfer agreement. Processed large-scale recording data, Matlab and Python analysis codes, histological images and probe reconstruction locations, 3D printable objects, and related data will be made available through the Medical Research Council (MRC) Brain Network Dynamics Unit (BNDU) data sharing platform upon publication (https://data.mrc.ox.ac.uk/mrcbndu/data-sets/search).

## Notes

**Conflict of interest:** The authors declare no competing financial interests associated with this study.

### Competing Interest Statement

The authors have declared no competing interest.

