## Supplementary Materials for "Higher-Order Thalamus is Pivotal in Schizophrenia-Associated Pathophysiology"

Jeffrey Stedehouder *et al*.

**The PDF file includes:**

Materials and Methods
Figs. S1 to S26
Tables S1 to S7
References

### Materials and Methods

#### Ethics

Experimental procedures were conducted according to the UK Animal Scientific Procedures Act (1986) and under personal and project licenses released by the Home Office following appropriate ethics review.

#### Mice

Homozygous mutant *Grin2a*$^{-/-}$ were obtained from BRC Riken (#RBRC02256), deposited by Dr. Motoya Katsuki.[46] Homozygous male *Grin2a*$^{-/-}$ were backcrossed with female wildtype C57BL/6J to obtain *Grin2a*$^{+/-}$ mice. Experimental cohorts were obtained from male mutant *Grin2a*$^{+/-}$ crossed with female wildtype C57BL/6J mice. *Grin2a*$^{+/-}$ and *Grin2a*$^{+/+}$ were born in near-Mendelian probabilities (~58%:42%), in equal male:female ratios (~50%:50%), and *Grin2a*$^{+/-}$ mice were healthy, viable, and displayed no overt growth or behavioral abnormalities. We did not observe any seizures in *Grin2a*$^{+/-}$ mice either phenotypically or in any electrophysiological measure.

We have used a total of 24 adult male mice in this study. For head-fixed recordings, adult male *Grin2a*$^{+/-}$ and *Grin2a*$^{+/+}$ mice were used between 8 and 20 weeks of age, weighing between ~25-35 g. All mice were group-housed pre-surgery and single-housed post-surgery throughout habituation, training and recording. Mice were kept in a 12-hour light/dark cycle (light on 07:00-19:00) in individually ventilated cages with constant temperature (~22°C) and humidity (40-70%). All experiments were performed during the light phase of the cycle. Food pellets were provided *ad libitum*. Mice had controlled access to water during training and recording.

Unless indicated otherwise, all experiments were performed in one-by-one pairs of heterozygous mutant *Grin2a*$^{+/-}$ and wildtype *Grin2a*$^{+/+}$ littermates, with consecutive surgery, habituation, training, recording, and perfusion, with balanced order of either *Grin2a*$^{+/-}$ - *Grin2a*$^{+/+}$ or *Grin2a*$^{+/+}$ - *Grin2a*$^{+/-}$ experiments. Manipulation experiments were performed in one-by-one *Grin2a*$^{+/-}$ hM3Dq-mCherry - *Grin2a*$^{+/+}$ mCherry pairs.

#### Surgery

Headpost surgery was performed as described previously.[47] Briefly, mice were anaesthetized under isoflurane (~4-5%) followed by maintenance isoflurane (~1.5-2%) at a flow of ~1.5 L/min $O_2$. Analgesia was provided using Vetergesic (0.08 mg/kg in sterile saline) administered subcutaneously after induction of anesthesia. The skin and periosteum on the dorsal surface of the skull was removed, and the surrounding skin secured to the skull using cyanoacrylate (Vetbond, 3M, Minnesota, USA). The skull was prepared using 3% $H_2O_2$ and etched using a bone scraper. A custom-made titanium headpost was implanted on each animal (Get It Made Ltd, London; 0.7 g, internal diameter 9 mm), aligned to lambda and the midline and secured using dental cement (Super Bond C&B, Parkell). A reference screw (Precision Technology, East Grinstead) was wrapped in silver wire and positioned into a craniotomy over right cerebellum and fixed in place using dental cement (Jet Denture Repair Powder: Lang Dental, Illinois, USA; Meadway Repair Liquid: MR. Dental, Surrey, UK). A custom-designed, 3D-printed shield was secured to the top of the headplate. Mice were allowed to recover for 7 days, followed by habituation to head-fixation and behavioral training (see below).

Twenty-eight days after headplate surgery, anesthetized recovery surgery was performed to create small craniotomies (~0.5-1 mm) above each proposed recording site, leaving the dura intact. Coordinates and planned craniotomy sites were selected based on previous work.[47] Following surgery, the craniotomies were covered with DuraGel (Cambridge Neurotech, UK) and overlaid with silicon (Body Double, Smooth-On).

In chemogenetic experiments, adult male *Grin2a*$^{+/-}$ mutant animals (aged ~12-20 weeks) received during headpost surgeries intracranial transduction of AAVs containing activating DREADDs (hM3Dq-mCherry), as described previously.[48] AAVs were diluted to a titre of $5.0 \times 10^{11}$ in sterile saline, and 300 nl was infused per site using a Hamilton 2.5 µl Syringe (7632-01, Hamilton Europe, Romania) with a 33-gauge, 20 mm, 4-12° needle (7803-05, Hamilton Europe, Romania). Needles were lowered at a speed of 1 mm/min and left in place for 5 minutes before start of infusion. AAVs were infused with a speed of 100 nl/min, followed by another 5 minutes of leaving the needle in place, before raising the needle at a speed of 1 mm/min. The following coordinates were used to target bilateral higher-order thalamus (from bregma (in mm)): AP: -1.20; ML: ±0.35; DV: -2.9.

**Adeno-associated constructs**

The following adeno-associated viral (AAV) constructs were obtained from Viral Vector Facility (Zürich, Switzerland):

#v96-5: AAV2/5-mCaMKIIα-hM3D(Gq)_mCherry; titre 5.6x$10^{12}$ vg/ml
#v199-5: AAV2/5-mCaMKIIα-mCherry; titre 6.0x$10^{12}$ vg/ml

**Behavioral task**

The bar pressing task was adapted from a previous study.[49] Mice were habituated to head-fixation for ~7 days in incremental durations before onset of behavioral training. Controlled access to fluids was used to motivate mice during training and performance of the task, with mice held at ~88-89% of pre-headplate starting body weight during behavioral experiments, with no difference between genotypes (**Table S1, Table S5**). Mice were placed in a tube with custom printed, cork-lined interior lining with forepaws resting on two metal bars under head-fixation. A moveable waterspout was placed in ventral and anterior to the mouse. A PyControl system was used to control dispensation of sucrose (5% (wt/vl) sucrose in water, ~5 µl) droplets through the spout. Room illumination was set to ~30-50 lux.

Mice were trained to push the left metal bar self-paced forward over a distance of ~5-10 mm and return it to starting location. Each bar push was coupled to a pure tone (2-16 kHz, 100 ms, ~75 dB) on 100% of the trials at bar push onset. The bar had to be returned to starting position between 250 and 1000 ms for it to be considered a correct trial and be rewarded (‘success’). Further details on sound presentation and behavioral performance can be found in a parallel manuscript.

Mice were trained on ~20-35 min sessions per day for two weeks (mean: 11.75 sessions; range 11 – 13 sessions). Over consecutive training sessions, mice performed a total of ~5000 trials (mean: 5065 trials; range 2572-7933 trials) before continuing with craniotomy, a number no different between genotypes (*Grin2a*$^{+/+}$: 5067 trials; *Grin2a*$^{+/-}$: 5063 trials; $t_{14} = 0.006$, $p = 0.996$, unpaired *t*-test). We note that hM3Dq-mCherry *Grin2a*$^{+/-}$ animals performed marginally significant lower number of trials compared to mCherry *Grin2a*$^{+/-}$ controls during *training*, in absence of CNO (*Grin2a*$^{+/-}$ hM3Dq-mCherry: 2756 trials; *Grin2a*$^{+/-}$ mCherry: 4575 trials; $t_5 = 2.520$, $p = 0.0453$, unpaired *t*-test, **Table S5**).

During recordings, mice were exposed to the same bar pushing paradigm, with some small changes and divided into three stages: In stage (1), mice performed 30 standard trials similar as during training. In stage (2), mice performed up to 500 trials or trials for 30 mins, whichever was attained earlier. During this stage, bar presses could elicit a tone of 1 octave different from the training tone on a subset (15%) of trials. In another 15% of trials, bar presses did not elicit any tone. In addition, every 20-40 seconds, the target tone was played passively in absent of any bar press. All remaining trials were as in stage (1). In stage (3), the bar was locked in place and mice were passively exposed to a series of 30 tones across 4 frequencies with an interval of ~2 seconds (total 120 tones).

**Probe trajectory planning**

Probe trajectories were planned using the Allen Institute Common Coordinate Framework (CCFv3)[50] GUI tool and PinPoint[51], with the following estimated coordinates (mm from Bregma):

Frontal cortex: +2.00 AP, +0.50-1 ML, 5-15º ML tilt, 0-15º AP tilt
Striatum: +1.00 AP, ±1.00-2.00 ML, 5-15º ML tilt, 0º AP tilt
Hippocampus: -1.00-1.75 AP, +1.00-1.50 ML, 5-15º ML tilt, 0º AP tilt

Auditory cortex: -2.50 AP, -4.70-4.20 ML, 40-65º ML tilt, -10-15º AP tilt

**Electrophysiological recordings**

Up to four Neuropixels probes (Neuropixels 1.0 single-shank, IMEC, Belgium) were used simultaneously to record acutely from multiple brain areas during rest and during performance on the bar pressing task. Custom 3D printed probe holders and headstage holders (10k resin, Formlabs) were used to support the probes and headstages, respectively. Probes were lowered using individual automated single-axis motors (700-105140-IVM, Scientifica, United Kingdom) mounted on a custom-designed frame with a Kopf micromanipulator (DKI 1760-61-SB) a speed of 3-4 µm/s till a depth of ~3500 µm, generally leaving top probe channels outside of the brain for realignment. In a subset of experiments, deeper probes were lowered with a speed of 5-6 µm/s for the first ~2000 µm, followed by a speed of 3-4 µm/s till recording depth. Upon target location, probes were retracted by 100 µm at a speed of 3 µm/s and allowed to settle in place for 5 mins before recording. Probes were grounded to a cerebellar reference screw and were referenced to the probe tip. Sterile saline was applied to the craniotomy during probe lowering, recording and retraction.

Electrophysiological recordings were made using OpenEphys software[52] at 30 kHz. Each recording session lasted ~35 minutes (range 28 - 39 mins), with an average total head-fixation duration of <120 mins. Recording sessions occurred once per day per mouse for ~5 days (range 4 – 6 days). PyControl[53] was used to acquire event times.

Neuropixels probes were sharpened using a repurposed hard drive disk. The ground and reference pads of the probes were shortened and soldered to a wire that connected to the animal and ground. Before each recording, a fluorescent lipophilic membrane stain 1,1'-dioctadecyl-3,3,3',3' -tetramethylcarbocyanine perchlorate (DiI; Invitrogen, California, USA; 1-2 mg/ml in isopropanol) was applied to the probes ~15 mins before brain insertion. In a subset of experiments, we employed DiO (V22886, Invitrogen; California, USA; 1-2 mg/ml in $H_2O$) instead. After recording, probes were retracted at a speed of 5-6 µm/s. At the end of a recording session, the craniotomy was again covered with DuraGel and overlaid with silicone. After recording, probes were cleaned in 1% tergazyme, de-ionised water, and isopropanol.

For chemogenetic experiments, *Grin2a*$^{+/-}$ animals were habituated to additional fixation for intraperitoneal injection during the second week of training for ~7 days, ~5 mins before onset of head-fixation. On recording days, clozapine n-oxide (CNO, 6329, Tocris Bioscience, Bristol, United Kingdom) was made fresh in sterile saline (0.5 mg/ml) and injected daily (1 mg/kg) intraperitoneally to both hM3Dq-mCherry and mCherry *Grin2a*$^{+/-}$ animals ~5 mins before head-fixation onset, and ~40-45 mins before recording start.

**Histology**

Brains were processed as previously described.[54] One day after the last recording, mice were anesthetized using a terminal pentobarbital intraperitoneal injection (200 mg/kg). Each mouse was transcardially perfused with ~5-10 ml 0.1 M phosphate buffer (PB, pH 7.4) followed by ~50 ml of 4% paraformaldehyde solution (PFA) in PB. Brains were removed and post-fixed for 2 hours at 4ºC in 4% PFA before washing with PB and placed overnight in 10% sucrose/0.1 M PB. The following day, brains were embedding in 12% gelatin/10% sucrose using custom, 3d-printed embedding wells, post-fixed in 10% PFA/30% sucrose for 2 hours and subsequently stored in 30% sucrose until cutting.

Whole brains except the cerebellum were coronally sectioned into 50 µm slices using a sliding microtome (Expredia HM450*,* Fisher Scientific*,* UK), with slices sequentially collected in 0.1 M PB. Slices were immersed in 1:5000 DAPI nuclear marker for 5 mins, mounted on glass slides, and coverslipped using Vectashield mounting medium (H-1000).

Slices across the full rostrocaudal axis of the brain except the cerebellum were acquired using an Axio Imager M2 epifluorescent microscope (Zeiss), equipped with Plan-Achromat objective lenses, a Hamatsu Flash 4.0 LT camera (C10600) and Colibri LEDs. ZEN Blue software was used to acquire tiled single plane images of each entire slice with a 10x objective. Slide overviews were obtained at 12 x 5 tiles overview at 2.5x, followed by 9 x 5 tiles per section at 10x (10% tile overlap), 2000x2000 pixels at 26 µm/pixel. DAPI, DiO, DiI, and mCherry were imaged using Colibri 5/7 LEDs at wavelength 385, 475, 555, and 555, respectively.

Probe tracks were reconstructed using the Allen Institute Common Coordinate Framework (CCFv3) GUI MatLab tool, SHARP-Track.[55] Briefly, coronal images including the entire brain except the cerebellum were processed and aligned to the matching coordinates on the reference atlas, with probe tracks manually traced (~20 probe points per track) across every 2$^{nd}$ image at ~100 µm anterioposterior intervals. Surface analysis did not reveal any overt changes in brain anatomy (*data not shown*).

Probe tracks could be equally recovered from mutant and wildtype mice, with similar numbers of employed probe points between genotypes or manipulations (**Table S1, Table S5**), and probe tracks were assigned to recording days based on a combination of insertion location within the craniotomy, insertion depth, and insertion angle. Probe track data were utilized to map brain areas to channels along the length of the Neuropixels probe.

hM3Dq-mCherry and mCherry virus reconstruction were performed and visualized using a custom version of SHARP-Track.[55] In brief, coronal images including the full rostrocaudal extent of virus expression were processed and aligned to the matching coordinates on the reference atlas, with areas containing mCherry-positive somata traced across every 4$^{th}$ image at ~200 µm anterioposterior intervals.

**Regional alignment**

To align channels from reconstructed tracks to individual brain areas, we utilized single neuron and local field potential (LFP) information for manual alignment of individual probe channels with electrophysiology using a custom-written GUI. We employed between ~1-6 previously reported electrophysiological landmarks (**Fig. S3, Table S2**) per probe to align landmarks to regional boundaries. We opted to employ previously validated landmarks in the form of neuron numbers, spike rate, waveform peak amplitude, LFP power, LFP power change and LFP cross-correlation across channels. Briefly, raw LFP across all channels was corrected through common-average referencing: Subtracting each channel's median to remove baseline offsets, then subtracting the median across all channels at each time point to remove artefacts. LFP cross-correlation was calculated by obtaining a random 4-min window of corrected LFP, filtering the signal for the 1-80 Hz range (Butterworth, 2$^{nd}$ order), and calculating correlations across all channels across the time window.

Similar numbers of landmarks were employed between mutant mice and controls per probe or between manipulations (average landmark number: *Grin2a*$^{+/-}$: 2.6 ± 0.1, *Grin2a*$^{+/+}$: 2.6 ± 0.1; *Grin2a*$^{+/-}$ *(hM3Dq-mCherry)*: 2.2 ± 0.1, *Grin2a*$^{+/-}$*(mCherry)*: 2.4 ± 0.1). We took particular care that alignment in wildtype or mutant probes was not dependent on any singular landmark to avoid biasing due to to-be-discovered electrophysiological alterations in *Grin2a*$^{+/-}$ mutant mice. Updated brain regions were adjusted and loaded back into the CellExplorer[56] environment.

In a subset of follow-up analyses, we split hippocampal regions (CA1, CA2, CA3, and DG) into dorsal versus ventral hippocampus[19] by splitting based on recording probe location: Dorsal hippocampus was defined as hippocampus targeted by perpendicular probes travering along the dorsoventral gradient through motor cortex, somatosensory cortex and retrosplenial cortex (probe insertion location: AP -0.5 - -2.0, ML 0.5-2.0, from bregma (mm)), while ventral hippocampus was defined as hippocampus targeted by oblique angled probes traversing auditory cortex or ventral visual cortex (probe insertion location: AP -2.0 -3.5, ML 3.5-4.5, from bregma (mm)).

**Anatomical nomenclature**

We used a combination of anatomical atlases throughout this study. Franklin and Paxinos[57] was utilized for initial planning of craniotomies and headpost location, for planning of viral injection experiments, and used to create first version of our sagittal brain view schematics used here. Sagittal schematic depiction of effect sizes throughout this study were designed based on[57] at three mediolateral locations (ML: 0.48, 2.18, and 3.72 mm) using custom Matlab code. Final probe reconstruction, region nomenclature, and clustering were based on Allen Brain Institute CCFv3[50] (**Table S2**). Neurons assigned to white matter regions were excluded from further analysis.

In one analysis (**Fig. S8**), thalamic and cortical regions were re-clustered based on anatomical thalamocortical connection patterns as per[18].

**Session Curation**

Twelve sessions were removed as animals received a pharmacological intervention. One session was removed due to technical problems during recording. Twenty sessions (one hM3Dq-mCherry animal) were removed as we failed to recover viral expression in consecutive histology. All sessions are removed before final analysis described throughout the results.

**Data analysis**

Electrophysiological data were spike-sorted using Kilosort 3.0[58] using default parameters. Neurons labelled as ‘good’ by Kilosort 3.0 were selected for downstream analyses. Non-somatic spikes[59] were removed with custom-written Matlab scripts, removing single-channel waveforms that either contained a first peak with an amplitude larger than the main waveform trough, or when the second peak following the trough was larger in amplitude than the main waveform trough. No difference in non-somatic spike removal was observed between genotypes (~1.7%) or manipulations (~1.2%). Further quality control was performed utilizing a combination of previously described[7,9,60] spiking and waveform metrics including neurons with presence ratio (>0.7), total spike count (>300), and average waveform amplitude (>20 µV), filtered using BombCell[61] and custom-written Matlab scripts. To minimize any bias, we did not perform any manual neuron curation.

In total, 106,972 neurons (~71.9%) were removed after Kilosort output for comparative experiments, an equivalent number between genotypes ($Grin2a^{+/-}$: 53,443 neurons (~72.1%), $Grin2a^{+/+}$: 53,529 neurons (~72.6%)) and 14,003 neurons ( ~72.2%) were removed from manipulative experiments ($Grin2a^{+/}$ (hM3Dq-mCherry): 17,368 neurons (~71.3%), $Grin2a^{+/-}$ (mCherry): 13,635 neurons (~73.2%)). Neurons were imported into CellExplorer [56] environment and combined with the region location (see above), as well as event-related data acquired from PyControl [53] to acquire region-relevant neural output during and outside of the task.

Unless otherwise indicated, all electrophysiological analyses were performed across the full recording period. Spontaneous firing rate was calculated by the total number of spikes across the full duration of the recording. CV was calculated as standard deviation of the ISIs divided by the mean of the ISIs per neuron. CV2 was calculated as $2 \times (ISI_{n+1} - ISI_n)/ISI_{n+1} - ISI_n)$ per neuron. Bursting behavior was calculated as described previously.[62] Two or more identified spikes with a consecutive inter-spike interval (ISI) lower than 6 ms were classified as bursts. Spikes with ISIs of >6 ms were considered ‘single spikes’. The burst index was calculated by dividing the number of bursts by the total number of spikes (ratio, %), across the full recording duration. Neurons were considered as ‘bursting’ if they fired at least one burst across the full recording period.

Neurons were assigned to the probe channel where its mean waveform amplitude was largest. Average single-channel waveform characteristics were calculated from waveforms across the full recording duration and included the following characteristics: Amplitude was calculated as the peak negative deflection from baseline. Trough-to-peak latencies (in ms) were calculated from refined waveforms (see below) by subtracting the trough negative deflection latency from the peak positive deflection latency following the trough. Shoulder asymmetry was calculated as the ratio between the positive peak preceding the trough and the positive peak following the trough. Peak-trough ratio was calculated as the ratio between the amplitude of the first peak preceding the trough and the amplitude of the trough negative deflection following the first peak.

To examine the effects of thalamus rate enhancement on brain-wide cross-parameter electrophysiology, we computed the ‘rescue score’ per region, calculated as

$$Rescue\ score = \frac{X(Grin2a\ het) - X(Grin2a\ het\ Dq)}{X(Grin2a\ het) - X(Grin2a\ wt)}$$

with $X$ denoting the mean or median value for all neurons for any given region for firing rate, burst firing, or co-activity for a given genotype or manipulation. Rescue scores were capped at 0 and 1, and interpreted as 1 (rescue), 0 (no rescue), or $0<x<1$ (partial rescue).

**Cell type classification**

Cell-type classification was processed as previously described.[63] In brief, all waveforms consisting of 48 data points from averaged single-channel waveforms at their peak channel for individual neurons were increased by a factor of 100 to 4800 data points using quadratic interpolation in Matlab. Trough-to-peak latencies (in ms) were calculated from these refined waveforms by subtracting the trough negative deflection latency from the peak positive deflection latency following the trough, and trough-to-peak latencies showed minimal differences across genotypes (**Figs. S4, 6**). The latencies were fitted with a 1-dimensional, 2-component Gaussian Mixture Model per grouped region for each genotype and the classification threshold was set at the intersect of the two Gaussian components. For striatum and hippocampus, a 3-component Gaussian Mixture Model was used instead which showed a superior fit ($p < 0.001$).

Segregation into regular-spiking neurons and fast-spiking neurons was performed for cortex, hippocampus, and striatum (excluding lateral septum). Neurons with a trough-to-peak latency below the intersect were classified as fast-spiking neurons, while neurons with a trough-to-peak latency above the intersect were classified as regular-spiking neurons. Regular-spiking and fast-spiking neurons differed from one another across a range of electrophysiological and waveform characteristics (**Figs. S13-S15**).

**Co-activity analysis**

Co-activity analysis on neurons were performed as described previously.[38] In brief, neuronal activity across the full recording period (unless otherwise indicated) was binned and z-scored in 25-ms bins. In a subset of analyses, spikes occurring in bursts (interspike interval < 6 ms) where removed from the spike trains before binning and z-scoring. In another subset of analyses, neurons were split between regular-spiking and fast-spiking neurons, before binning and z-scoring. For the shuffle analysis, the binned and z-scored neuronal activity (25-ms) was randomly shuffled across the time bins for each individual neuron. This approach maintained average firing rates of individual neurons while breaking any temporal association within bins across neurons in the same brain region. An additional circular shuffling analysis was performed by maintaining the temporal order of time bins for each individual neuron (e.g. oscillations, state) but randomly shuffling all time bins forward or backward. This approach maintained average firing rates of individual neurons *and* temporal firing patterns or neuronal states, while breaking the temporal association within bins across neurons in the same brain region. To examine effect of time window, neuronal spiking activity was binned across various windows, including 10-ms, 25-ms, 50-ms, 100-ms, 200-ms, and 500-ms.

Pair-wise Spearman $\rho$ correlation values were calculated on this neural activity of putative pairs of neurons from the same animal, recording session, and brain region, and correlational values were consecutively averaged (mean) across all pairs per brain region. Autocorrelations were removed, and each pair was included once in calculation of each mean.

With these criteria, we examined a total of 7,177,469 putative pairs in *Grin2a*$^{+/+}$ mice ($n$ = 8) and 6,734,845 putative pairs in *Grin2a*$^{+/-}$ mice ($n$ = 8) including all neurons across all regions, and a total of 4,814,214 putative pairs in *Grin2a*$^{+/+}$ mice and 4,570,665 putative pairs in *Grin2a*$^{+/-}$ mice when only including regular-spiking neurons across cortex, striatum, and hippocampus. For manipulation experiments we examined a total of 3,175,478 putative pairs in *Grin2a*$^{+/-}$ mice (hM3Dq-mCherry, $n$ = 4) and 2,655,614 putative pairs in *Grin2a*$^{+/-}$ mice (mCherry, $n$ = 4) including all neurons across cortex, striatum, and hippocampus.

**MouseLight Neurons**

Reconstructed single thalamocortical neurons were obtained from the Janelia MouseLight browser (mouselight.janelia.org).[42] We obtained neurons with their somata residing in the following regions: MD ($n$ = 15 neurons): AA1624, AA1620, AA1612, AA1433, AA1423, AA1181, AA1168, AA0371, AA0370, AA0368, AA0363, AA0353, AA0095, AA0055, AA0054. PCN ($n$ = 14) neurons: AA1443, AA1437, AA1151, AA0598, AA0595, AA0372, AA364, AA0337, AA0335, AA0301, AA0298, AA0050, AA0047. CM ($n$ = 1) neurons: AA0074.

**Transcriptomic Data**

Expression data for *GRIN2A* across rodent brain were obtained from Allen Institute Database[64], (mouse.brain-map.org), experiment 71307394 ('Allen v1'), and 75081000 ('Allen v2') and Human Protein Atlas.[65] For both, hypothalamus and midbrain expression were combined into a new group 'Other', analogous to electrophysiology data.

**Statistics**

Sample sizes were not calculated but based on a previous study using large-scale recordings.[47] Experiments were not randomised and investigators were not blinded to allocation during experiments and outcome assessment, although all experiments between genotypes were performed in $Grin2a^{+/-}$ - $Grin2a^{+/+}$ pairs or $Grin2a^{+/-}$ (hM3Dq-mCherry) - $Grin2a^{+/-}$ (mCherry) pairs on similar times of day, and data was processed using standardized experimental and analysis pipelines. Statistical analysis was performed using GraphPad Prism v7 and Matlab. Where appropriate, data sets were analysed using Shapiro-Wilk test for normality. No outlier data were identified or removed unless indicated. Data sets with normal distributions (e.g. waveform metrics) were analysed for significance using unpaired Student's two-tailed $t$-test or analysis of variance (ANOVA) measures followed by *post hoc* comparisons. Data sets with non-normal distributions (e.g. firing rate, burst firing) were analysed using Mann-Whitney $U$ test, with multiple comparison thresholds indicated in Figures and Figure legends. Data distributions (e.g. bar press duration) were analysed with Kolmogorov-Smirnov test. Non-normal distributions are accompanied by median values, while normal distributions are accompanied by means. Correlations were examined using Pearson's (quality control metrics) or Spearman rank test (neuronal co-activity). Unless indicated otherwise, effects between genotypes or between manipulations were calculated as

$$\Delta = \frac{X(Grin2a\ het) - X(Grin2a\ wt)}{X(Grin2a\ wt)} * 100$$

with $X$ denoting the mean or median value for all neurons for any given region for a particular metric. Genotype effects were replicated across 4 cohorts of 4 mice, and manipulation data was replicated across 2 cohorts of 4 mice. Data are expressed as mean ± standard error throughout. $n$ indicate number of neurons. Exact $P$-values values are provided throughout the text, except when $P < 0.001$. Significance threshold was set at $P < 0.05$.

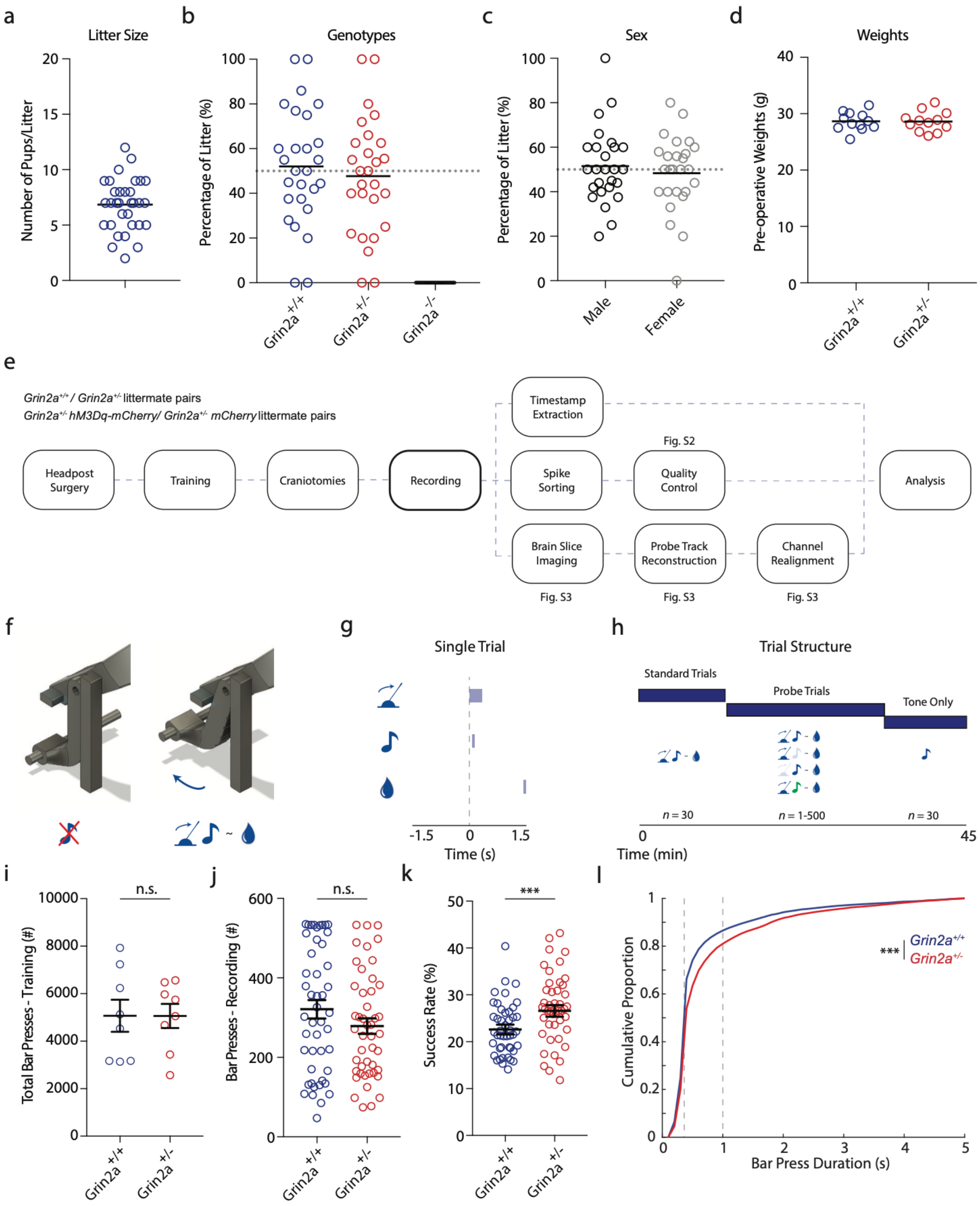

**Fig. S1. Standardized pipeline for Grin2a mutant and matched wildtype littermate control recordings**

**a)** Litter sizes of C57Bl/6J wildtype x *Grin2a*$^{+/-}$ heterozygote mutant breedings.

**b)** Genotype ratios of C57Bl/6J wildtype x *Grin2a*$^{+/-}$ heterozygote mutant breedings.

**c)** Sex ratios of C57Bl/6J wildtype x *Grin2a*$^{+/-}$ heterozygote mutant breedings.

**d)** Pre-surgery weights of *Grin2a*$^{+/-}$ and *Grin2a*$^{+/+}$ experimental mice. Bars denote group means.

**e)** Experimental pipeline. Pairs of *Grin2a*$^{+/-}$ mutant and *Grin2a*$^{+/+}$ wildtype littermates, or pairs of *Grin2a*$^{+/-}$ mutant littermates with either transduced hM3Dq-mCherry or mCherry fluorophore control, underwent surgery, habituation, training, and recording, followed by a standardized reconstruction and analysis pipeline.

**f-g)** Behavioral task setup. Adult, head-fixed mice were trained to press a bar self-paced for delayed reward. Bar press onset was accompanied by an auditory cue (100 ms, 2-16 kHz, tone frequency was set for each animal).

**h)** Recording trial structure. During recording, mice were allowed to press 30 ‘standard’ trials, followed by a period where the accompanying tone was absent, or replaced by a tone of an unexpected frequency, or a random tone was played in absence of the bar push. This was followed by a period where the bar was fixed in placed, and tones of varying frequencies (100 ms, 2-16 kHz) were played in random intervals. See **Materials and Methods**

**i-j)** *Grin2a*$^{+/-}$ mutant mice showed no change in total number of bar presses during trials (total, i), or during recording (each value is one recording day, j).

**k)** *Grin2a*$^{+/-}$ mutant mice showed an increase in the success rate of bar presses, ie. the number of trials the bar is returned to its starting position within 1 s.

**l)** *Grin2a*$^{+/-}$ mutant mice showed slower bar press movements compared to *Grin2a*$^{+/+}$ wildtype mice. Grey dotted lines indicate the window during which the bar had to be returned to base position for a reward. Statistics in **Table S7**.

***$p < 0.001$. Unpaired two-tailed *t*-test in (i) through (k). Kolmogorov-Smirnov test in (l). Error bars denote s.e.m.

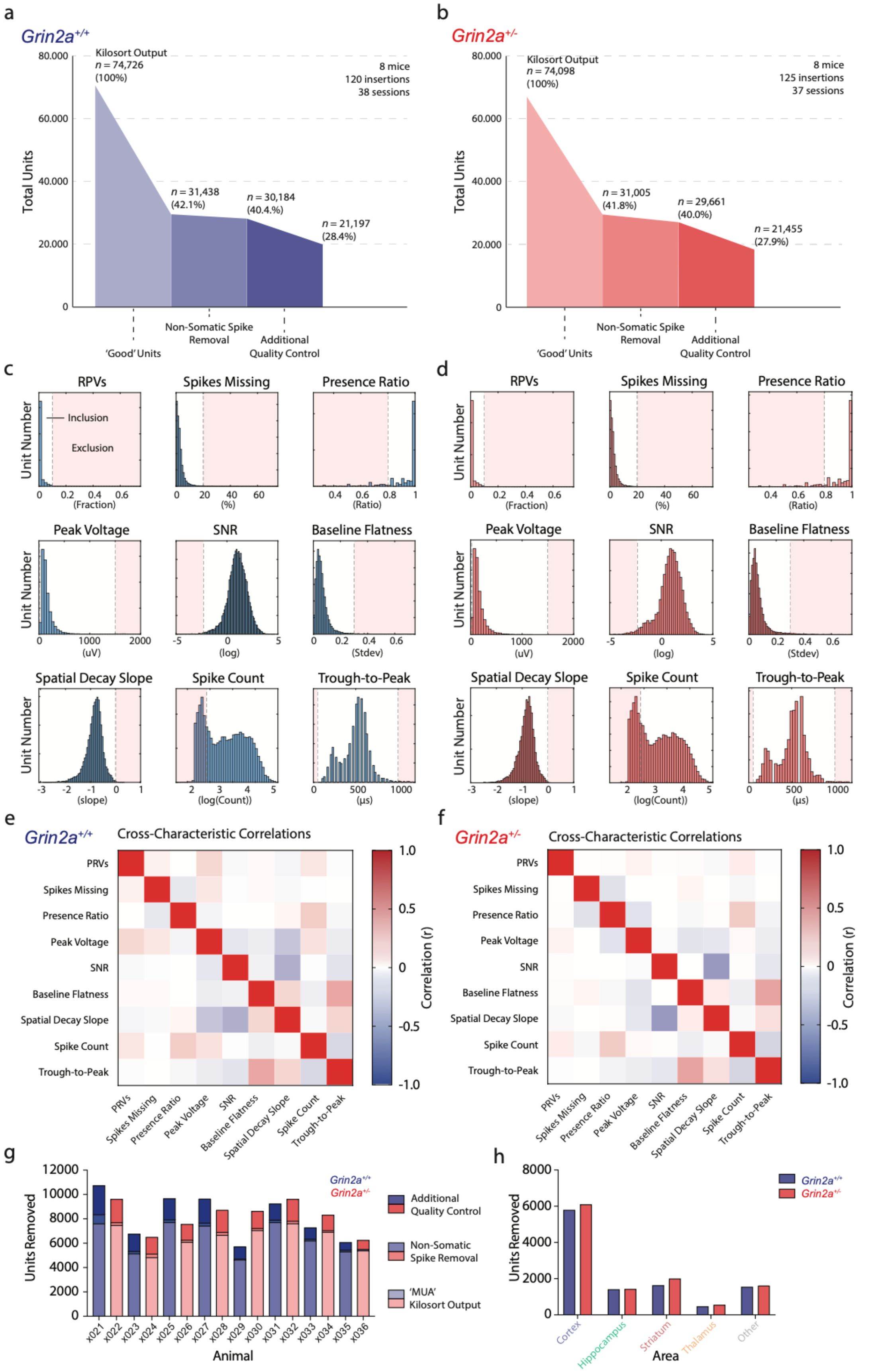
a
Grin2a+/+
Kilosort Output
n = 74,726
(100%)
8 mice
120 insertions
38 sessions
n = 31,438
(42.1%)
n = 30,184
(40.4.%)
n = 21,197
(28.4%)
Total Units
'Good' Units
Non-Somatic Spike Removal
Additional Quality Control
b
Grin2a+/-
Kilosort Output
n = 74,098
(100%)
8 mice
125 insertions
37 sessions
n = 31,005
(41.8%)
n = 29,661
(40.0%)
n = 21,455
(27.9%)
Total Units
'Good' Units
Non-Somatic Spike Removal
Additional Quality Control
c
RPVs
Spikes Missing
Presence Ratio
Inclusion
Exclusion
Unit Number
(Fraction)
(%)
(Ratio)
Peak Voltage
SNR
Baseline Flatness
(uV)
(log)
(Stdev)
Spatial Decay Slope
Spike Count
Trough-to-Peak
(slope)
(log(Count))
(µs)
d
RPVs
Spikes Missing
Presence Ratio
Unit Number
(Fraction)
(%)
(Ratio)
Peak Voltage
SNR
Baseline Flatness
(uV)
(log)
(Stdev)
Spatial Decay Slope
Spike Count
Trough-to-Peak
(slope)
(log(Count))
(µs)
e
Grin2a+/+
Cross-Characteristic Correlations
PRVs
Spikes Missing
Presence Ratio
Peak Voltage
SNR
Baseline Flatness
Spatial Decay Slope
Spike Count
Trough-to-Peak
Correlation (r)
f
Grin2a+/-
Cross-Characteristic Correlations
PRVs
Spikes Missing
Presence Ratio
Peak Voltage
SNR
Baseline Flatness
Spatial Decay Slope
Spike Count
Trough-to-Peak
Correlation (r)
g
Units Removed
Grin2a+/+
Grin2a+/-
Additional Quality Control
Non-Somatic Spike Removal
'MUA' Kilosort Output
x021 x022 x023 x024 x025 x026 x027 x028 x029 x030 x031 x032 x033 x034 x035 x036
Animal
h
Units Removed
Grin2a+/+
Grin2a+/-
Cortex
Hippocampus
Striatum
Thalamus
Other
Area

**Fig. S2. Spike sorting quality control.**

**a-b)** Flowchart showing quality control steps and neuron exclusion based on successive criteria in both wildtype control *Grin2a*$^{+/+}$ (a, blue) and *Grin2a*$^{+/-}$ (b, red) mutant mice.

**c-d)** Histograms of several quality control characteristics in control *Grin2a*$^{+/+}$ (c) and *Grin2a*$^{+/-}$ (d) mutant mice *before* the additional quality control step ($n$ = 31,438/8 mice; $n$ = 31,005/8 mice), along with exclusion criteria (vertical grey dashed lines) and to be further excluded neurons (red shaded areas).

**e-f)** Correlations for quality control metrics showing moderate correlations between several metrics but minimal differences between genotypes.

**g)** Neurons removed for various stages of QC, per mouse and genotype.

**h)** Neurons removed for various stages of QC, per area and genotype.

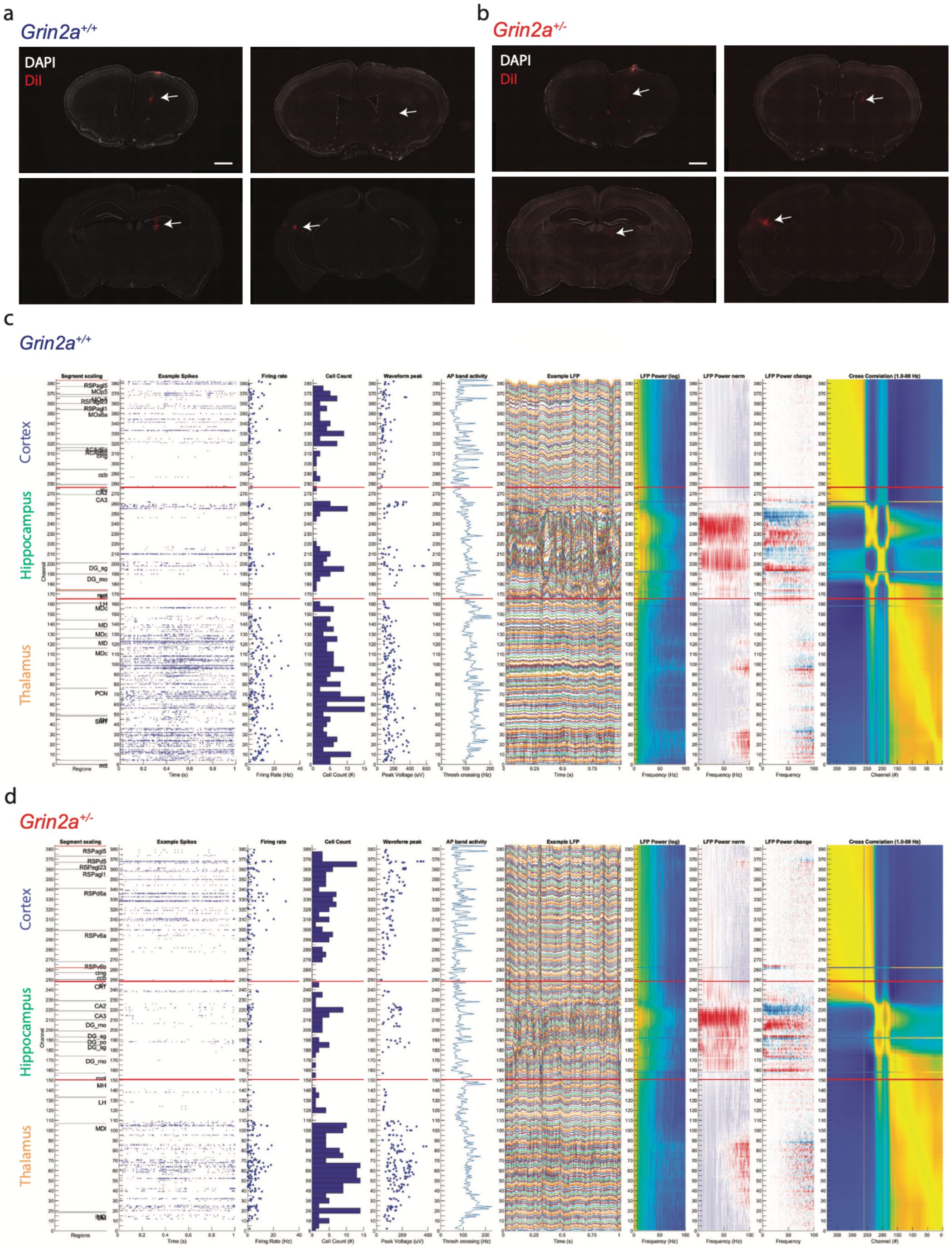

**Fig. S3. Histological reconstruction of probe tracks and channel realignment along the probe axis.** **a-b)** Low magnification epifluorescent microscope images showing coronal brain sections labeled with nuclear marker DAPI (white) with example DiI fluorescence (red) for *Grin2a*$^{+/+}$ (a) and *Grin2a*$^{+/-}$ (b) mice. Scale bar, 1 mm.

**c)** Screenshot of our custom region alignment GUI depicting region, spiking, and LFP information across channel depth for a single probe track from a *Grin2a*$^{+/+}$ mouse through cortex and hippocampus. Red lines indicate landmarks for rearranging channels for brain region borders. Cortex, hippocampus, and thalamus region indicated for visual aid.

**d)** As (c), but for a *Grin2a*$^{+/-}$ animal.

Region abbreviations in **Table S3**.

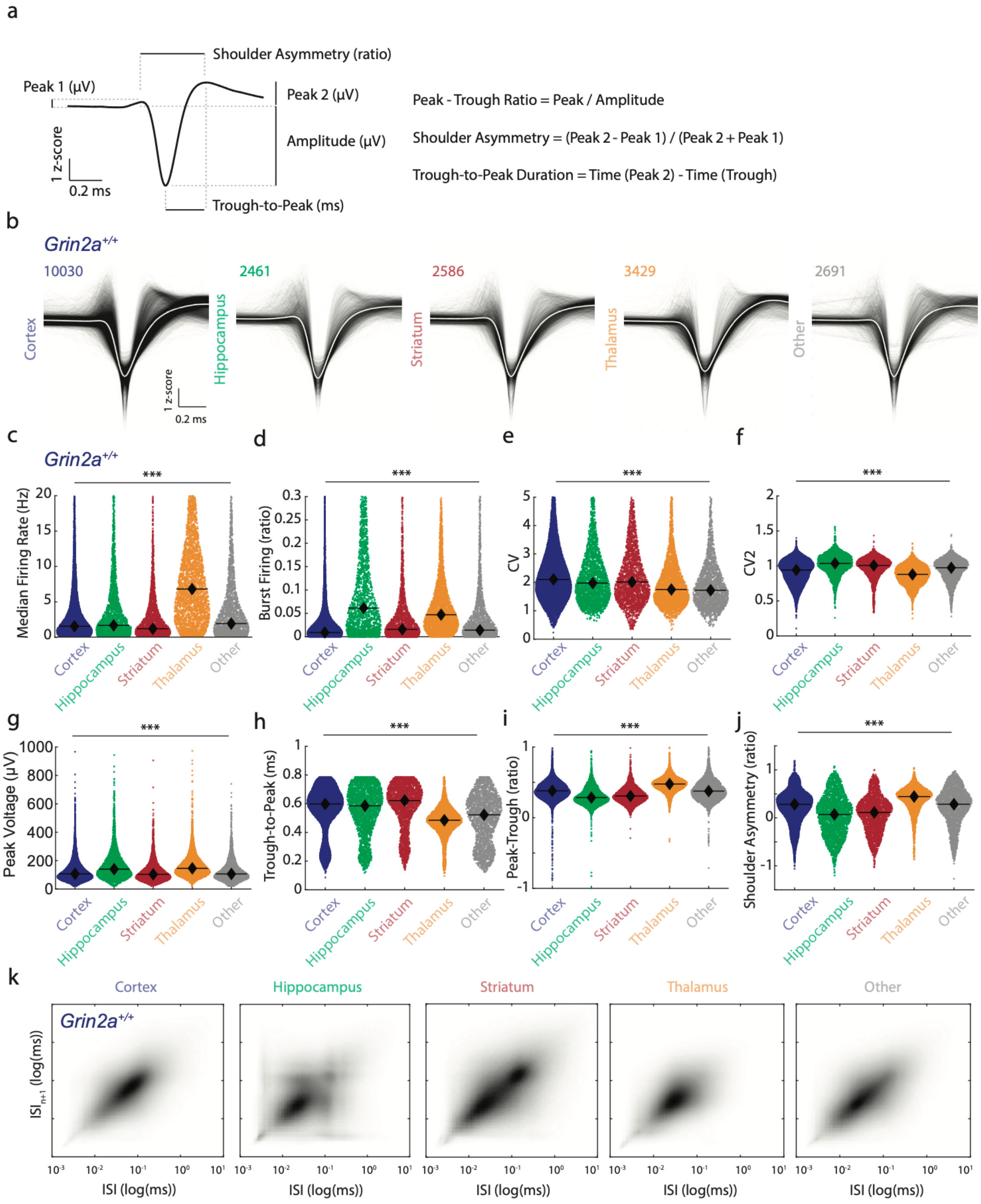

**Fig. S4. Broadly-defined area electrophysiological characteristics in wildtype mice.**

**a)** Visual depiction of calculations of waveform characteristics.

**b)** Average single-channel waveforms for all neurons from *Grin2a*$^{+/+}$ mice across cortex, hippocampus, striatum, thalamus and remaining regions, along with their neuron numbers. Scale bar, 0.2 ms, 1 Z-score.

**c-j)** Swarm plots for electrophysiological characteristics show differences per area, regardless of genotype, in firing rate (c), burst firing (d), CV (e), CV2 (f), peak voltage (g), trough-to-peak latency (h), peak-to-trough ratio (i), shoulder asymmetry (j). Black diamonds indicate median (c), (d), (g) or mean (e), (f), (h), (i), (j) values.

**k)** ISI return plots across various areas.
***$p < 0.001$. One-way ANOVA in (c) through (j). Statistics in **Table S7**.

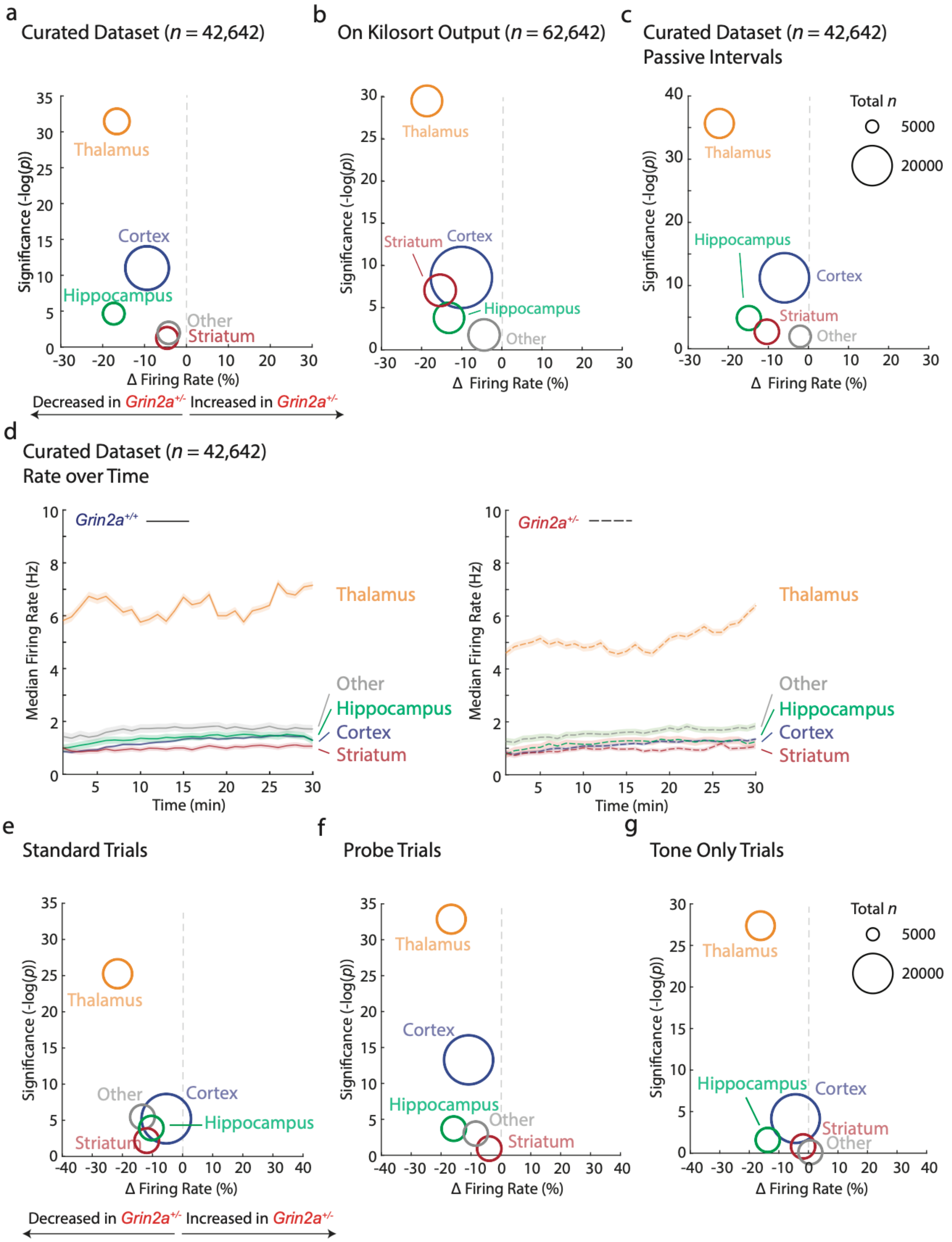

**Fig. S5. Electrophysiological characteristics without quality control, in passive intervals, and across different task phases.**

**a)** Original dataset depicting firing rate difference set out against pair-wise genotype significance per grouped area. Areas color-coded and circle size indicates total neuron number.

**b)** As (a), but on the unfiltered dataset of neurons directly obtained from Kilosort 3.0 without additional quality control. $n$ = 62,642 neurons/16 mice.
**c)** As (a), but only extracting firing rate from passive, intertrial intervals in absence of bar movement, sensory stimulus, or reward.
**d)** Depiction of median firing rate per area over time in 30-second bins. Median firing rate difference between genotypes remains stable over the recording period. *Grin2a*$^{+/+}$ (*left*; solid) and *Grin2a*$^{+/-}$ (*right*; dashed). Shaded areas indicated s.e.m.
**e-g)** As (a), but segregated per task phase: standard trials (e), probe trials (f), and tone-only trials (g; see **Materials and Methods**).
Statistics in **Table S7**.

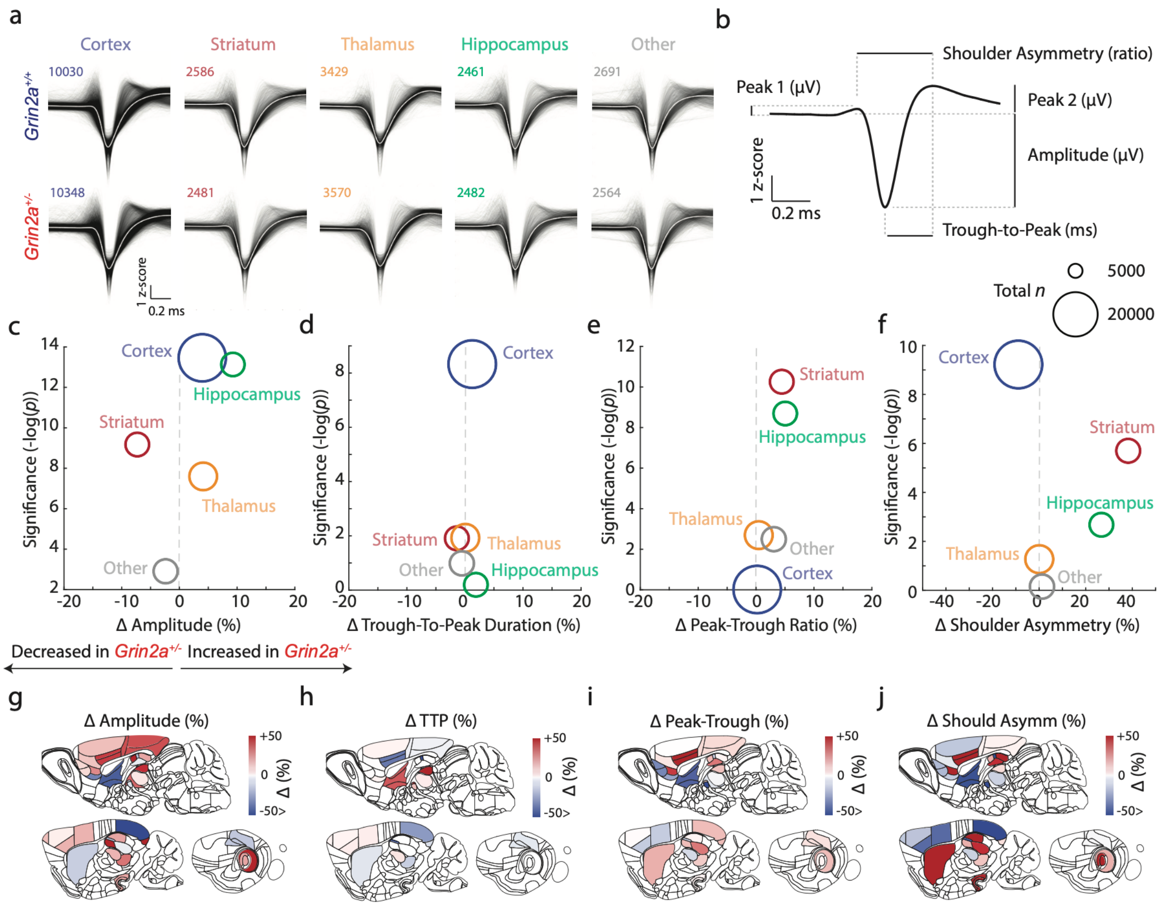

**Fig. S6.** ***Grin2a***$^{+/-}$ **mice display minor alterations in single-channel waveform characteristics.**
**a)** Average spike waveforms clustered per area for both *Grin2a*$^{+/+}$ (top) and *Grin2a*$^{+/-}$ (bottom) mice, along with the final recorded neuron number post-quality control. Scale bar, 1 z-score, 0.2 ms.
**b)** Schematic depiction of calculation of various waveform components.
**c-f)** Vulcano plots depicting waveform characteristic differences between genotypes per area for neurons single-channel amplitude (c), trough-to-peak duration (d), peak-trough ratio (e), and shoulder asymmetry (f). Areas color-coded and circle size indicates total neuron number.
**g-j)** Sagittal schematic brain region view depicting effect sizes between genotypes for waveform amplitude (g), trough-to-peak duration (h), peak-trough ratio (i), and shoulder asymmetry (j). Region abbreviations in **Table S3**. Statistics in **Table S7**.

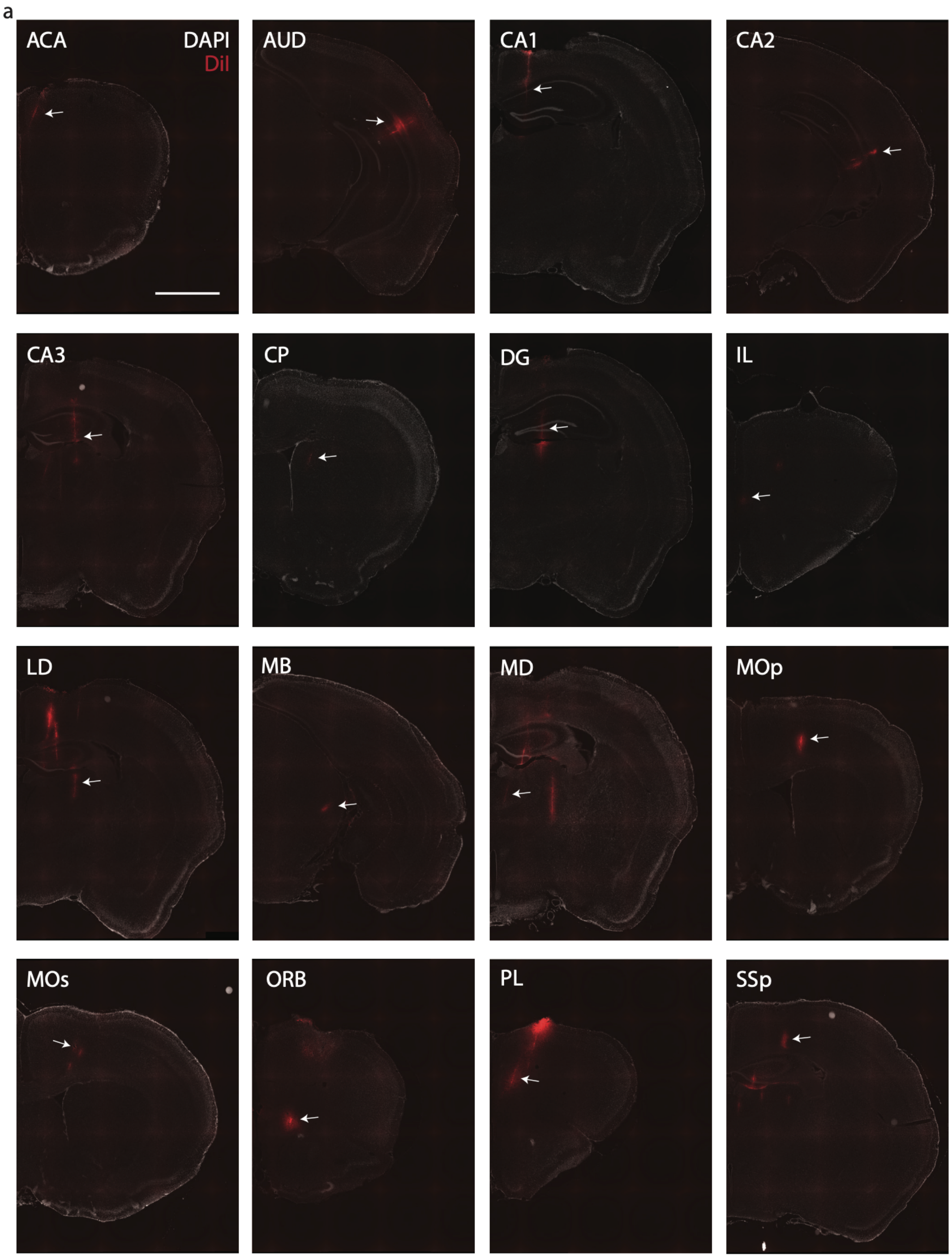

**Fig. S7. Exemplar histological confirmation of region-specific probe tracks.**
**a)** Low magnification epifluorescent microscope images showing coronal brain section labeled with nuclear marker DAPI (grey) with DiI fluorescence (red) for exemplar anterior cingulate area (ACA), auditory cortex (AUD), hippocampal CA1, CA2, CA3, caudate-putamen (CP), dentate gyrus (DG), infralimbic area (IL), laterodorsal thalamus (LD), midbrain (MB), mediodorsal thalamus (MD), primary motor cortex (MOp), secondary motor cortex (MOs), orbitofrontal cortex (ORB), prelimbic area, (PL), and somatosensory cortex (SSp), as indicated by the white arrow. Region abbreviations in **Table S3**. Scale bar, 2 mm.

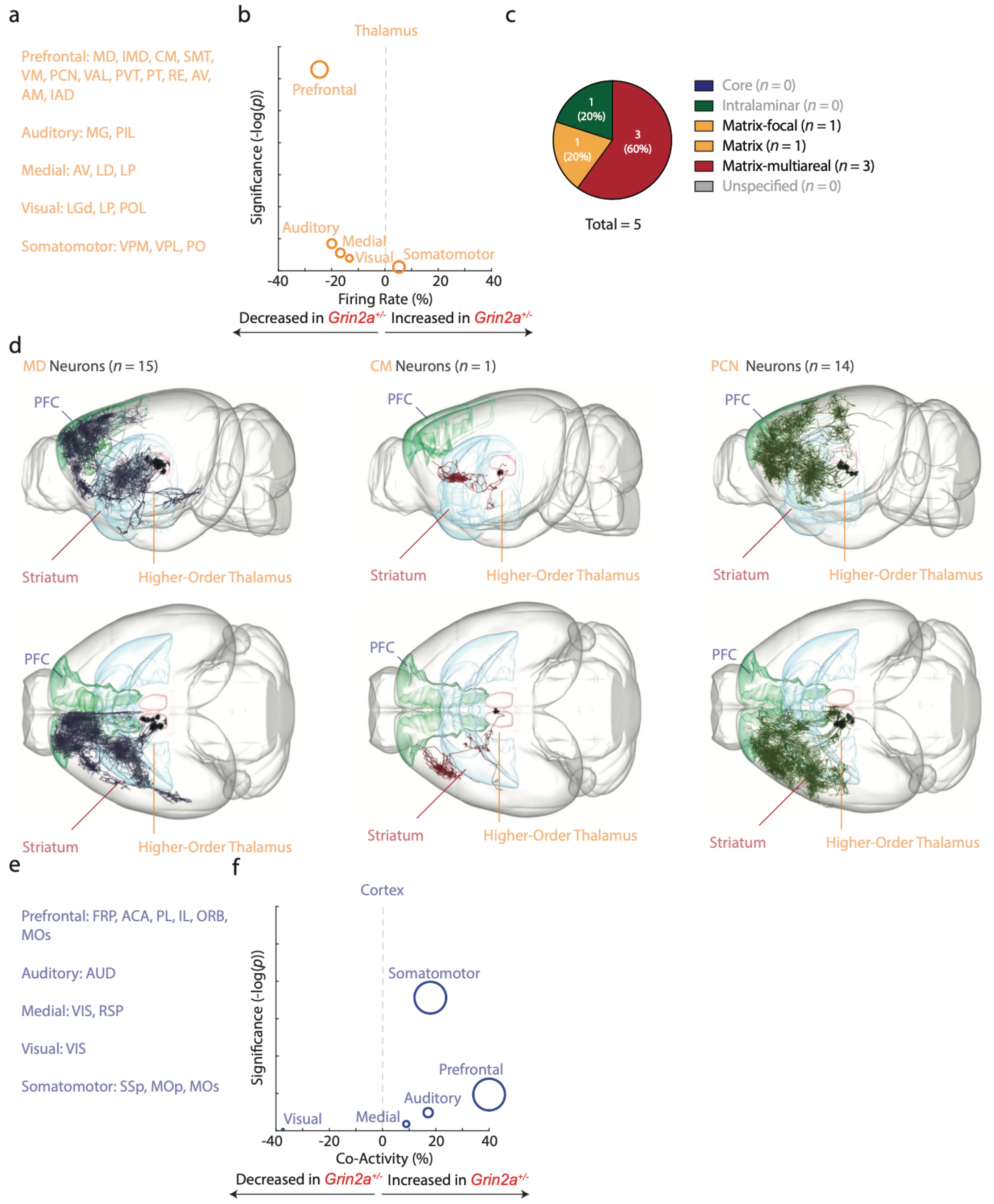

**Fig. S8. Affected higher-order thalamic and prefrontal nuclei in *Grin2a*$^{+/-}$ mice form a monosynaptic circuit.**

**a)** Alternative grouping of thalamic nuclei based on cortical projection pattern, as per [18].
**b)** Vulcano plot depicting the genotype spontaneous firing rate effects between *Grin2a*$^{+/-}$ and control mice for clustered thalamic regions suggests prominent reductions in ‘prefrontal’, higher-order thalamus.
**c)** Segregation of most prominently altered thalamic nuclei based on their axonal projection pattern, as per [18]. The majority of affected thalamic regions are matrix nuclei.
**d)** Single-neuron reconstructions for neurons in higher-order thalamic nuclei, as obtained from the MouseLight browser [42], for MD neurons (left), a CM neuron (mid), and PCN neurons (right). Prefrontal

cortex (PL, IL, MOs) is indicated in green, higher-order thalamus (MD) indicated in pink, striatum indicated in blue (CP). Higher-order thalamic nuclei broadly target striatum and prefrontal cortex.
**e)** Alternative grouping of cortical regions depending on their thalamic synaptic contacts, as per [18].
**f)** Vulcano plot depicting the neuronal co-activity effects between *Grin2a*$^{+/-}$ and control mice for clustered cortical regions suggests prominent alterations in prefrontal and somatomotor cortices.
Region abbreviations in **Table S3**. Statistics in **Table S7**.

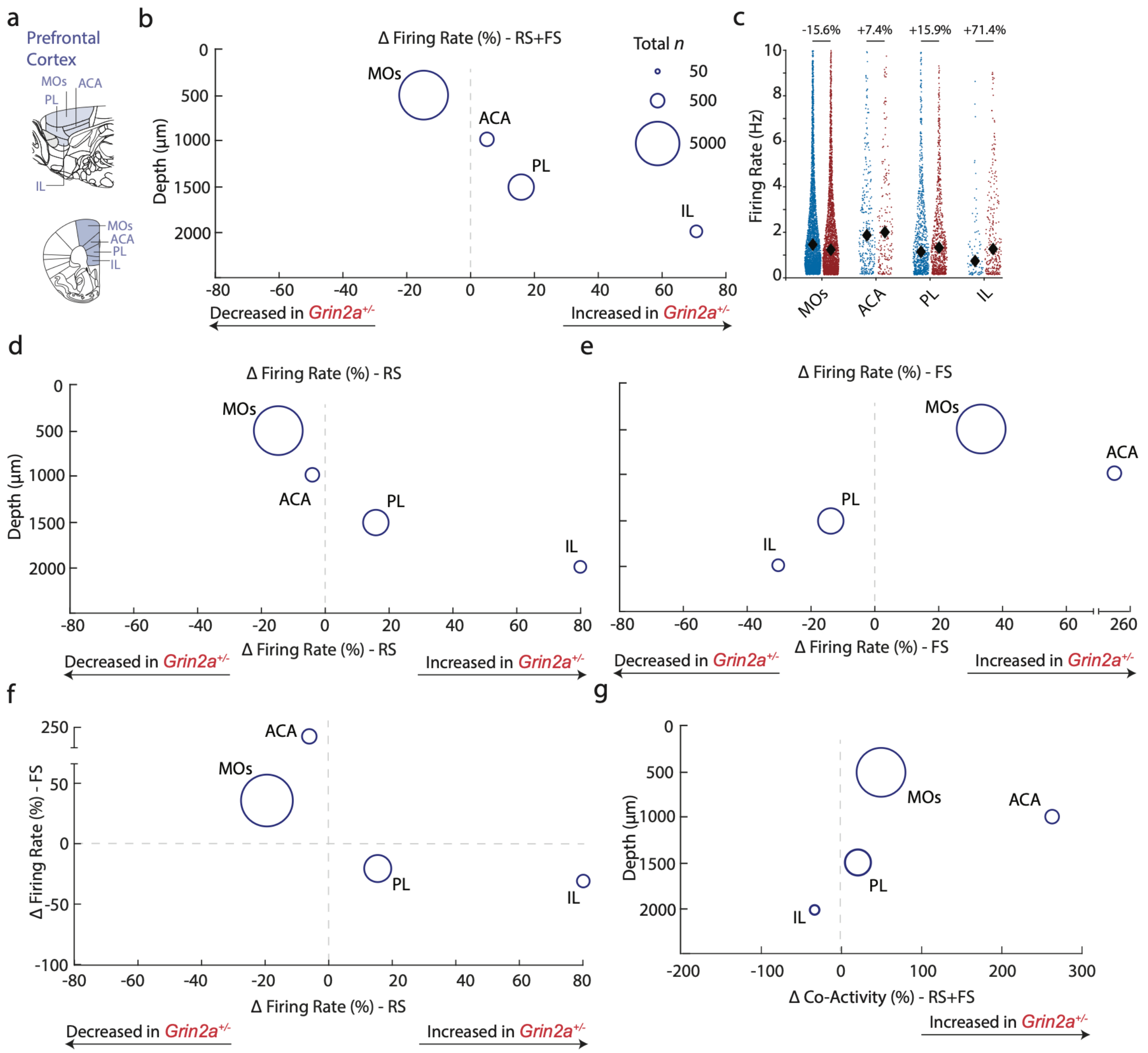

**Fig. S9. *Grin2a*$^{+/-}$ mice display a bidirectional, dorsoventral gradient in firing rate and co-activity across medial prefrontal cortex.**

**a)** Schematic depiction of mouse brain showing medial prefrontal cortex areas (blue) in sagittal view (top), and coronal view (bottom).

**b)** Firing rate alterations between genotypes show a non-uniform dorsoventral gradient in *Grin2a*$^{+/-}$ mutant animals compared to controls, with more ventral regions showing firing rate increases. Circle size indicates combined neuron number.

**c)** Swarm plots for firing rates of medial prefrontal cortical regions in (a)-(b) for *Grin2a*$^{+/+}$ wildtypes (blue) and *Grin2a*$^{+/-}$ mutant animals (red). Black diamonds indicate median values. Y-axis capped at 10 Hz for visibility.

**d)** As (b), but restricted to regular-spiking neurons.

**e)** As (b), but restricted to fast-spiking neurons.

**f)** As (b), but individually for regular-spiking and fast-spiking neurons. Regular-spiking and fast-spiking neurons show an inverse pattern of rate effects in medial prefrontal cortex.

**g)** Co-activity alterations including regular-spiking and fast-spiking neurons show a bidirectional effect between genotypes across the dorsoventral axis of medial prefrontal cortex**.** Circle size indicates total neuron number across both genotypes.

Region abbreviations in **Table S3**.

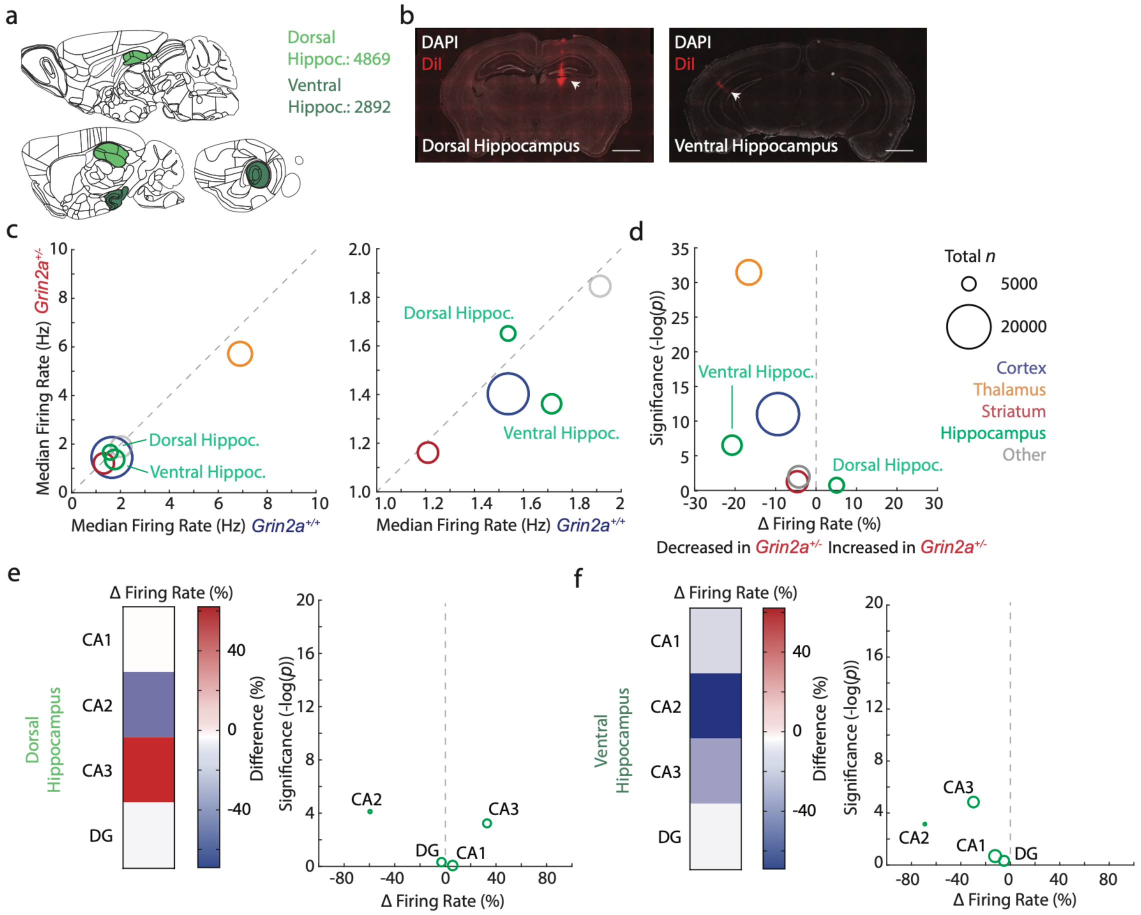

**Fig. S10.** ***Grin2a***$^{+/-}$ **mice display bidirectional dorsal-ventral hippocampal alterations.**

**a)** Schematic sagittal view of mouse brain indicating dorsal hippocampus (light green) and ventral hippocampus (dark green).

**b)** Exemplar epifluorescent microscopy images of coronal brain sections showing example probe tracks (DiI, red) of dorsal (left) and ventral hippocampal (right) probes (arrowheads). DAPI nuclear marker in white. Scale bar, 1 mm.

**c)** Firing rate for broad-defined areas for *Grin2a*$^{+/-}$ mutant and *Grin2a*$^{+/+}$ control animals with hippocampus split for dorsal and ventral compartments. Dorsal hippocampus shows slightly increased rates while ventral hippocampus showed reduced firing rates in mutant animals. Circle colours indicate grouping into areas and circle sizes indicated combined number of recorded neurons per area. Diagonal dashed grey line indicates no genotype difference.

**d)** Vulcano plot depicting firing rate effect sizes set out against pair-wise genotype significance for areas with hippocampus split for dorsal and ventral compartments. Horizontal dashed line indicated *p*-value threshold.

**e-f)** Heatmap and volcano plot exclusively for hippocampus, divided into dorsal (e) and ventral (f) hippocampus. Hippocampal CA3 shows minor increases in rate in dorsal hippocampus and decreases in ventral hippocampus.

Region abbreviations in **Table S3**. Statistics in **Table S7**.

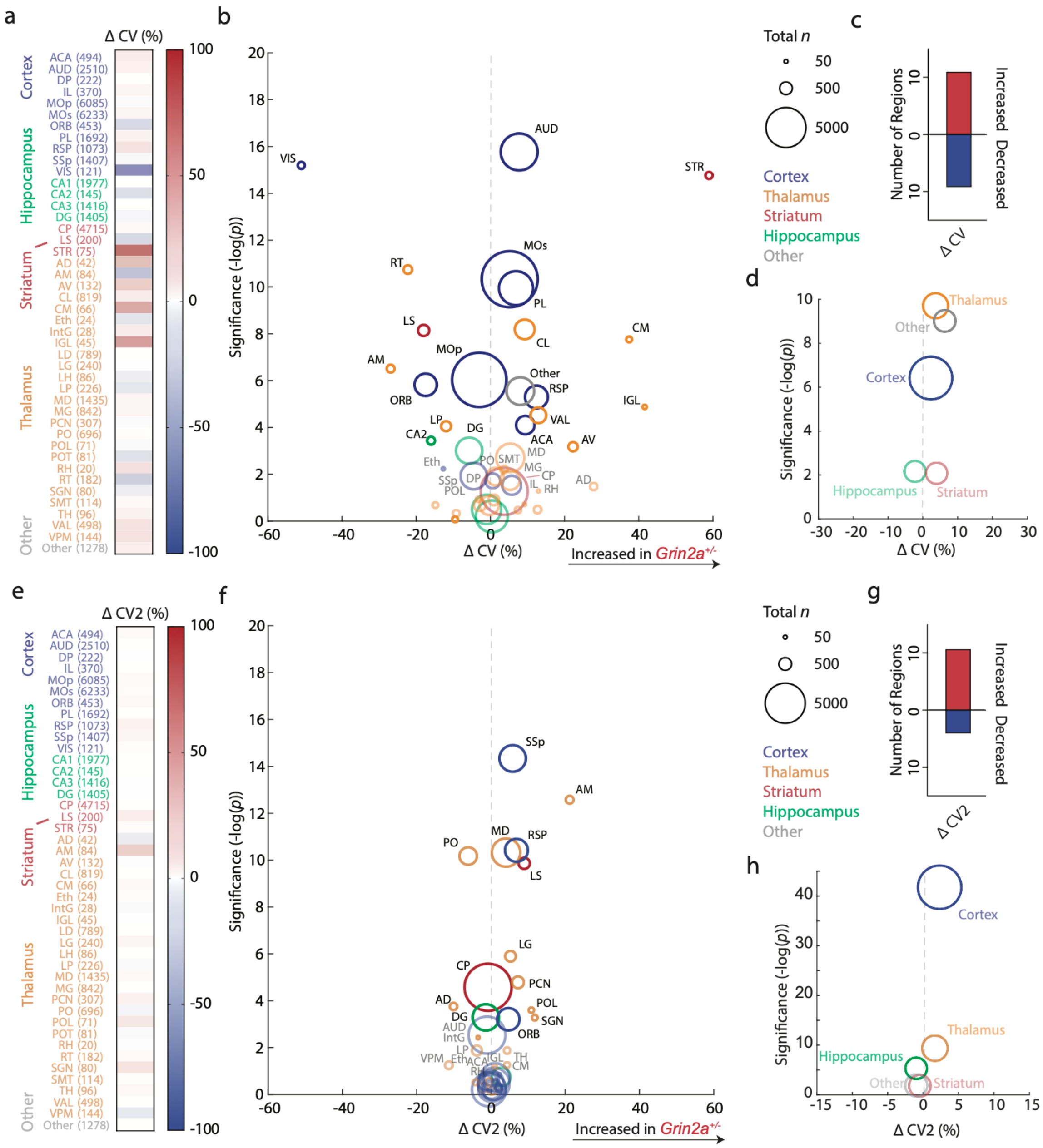

**Fig. S11.** ***Grin2a***$^{+/-}$ **mice display distributed and minor alterations in firing rate variability.**

**a)** Heatmap displaying genotype differences of firing rate variability (CV), ordered by broadly defined areas and alphabetically (left), along with the pair-wise genotype significance.

**b)** Vulcano plot showing effect size set out against pair-wise significance between genotypes per region. Circle sizes indicate total neuron number, colours indicate broadly defined area grouping. Greyed-out regions indicate levels of corrected significance.

**c)** *Grin2a*$^{+/-}$ mice show balanced increases and decreases in CV.

**d)** Grouping into broadly defined areas suggests modest increases in CV in *Grin2a*$^{+/-}$ mice in thalamus, cortex, and other regions.

**e-h)** As (a)-(d), but for CV2.

Region abbreviations in **Table S3**. Statistics in **Table S7**.

565

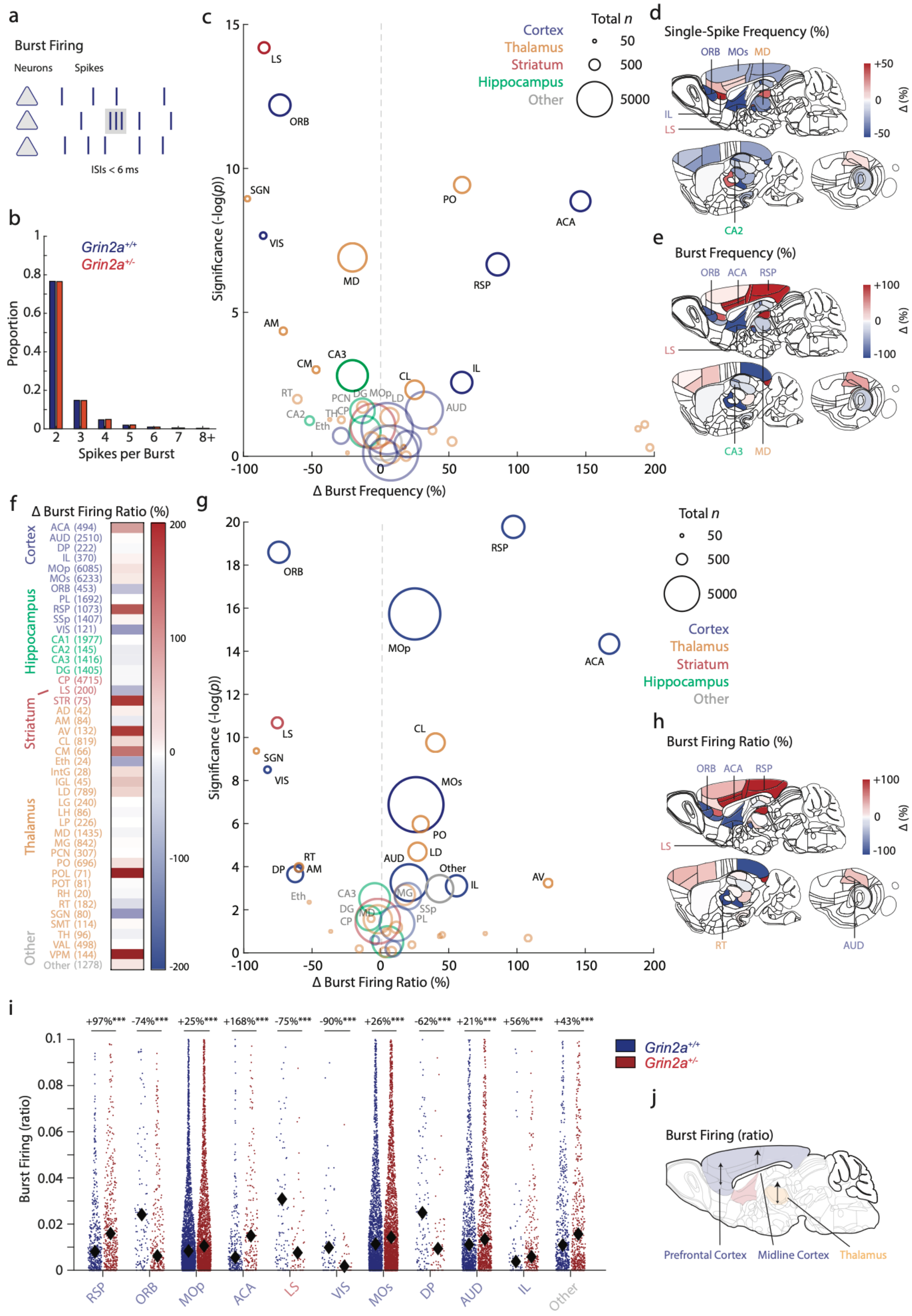

**Fig. S12.** ***Grin2a***$^{+/-}$ **mutant mice display distributed and graded increases in neuronal burst firing ratios.**

**a)** Schematic for burst firing analysis. Inter-spike intervals under 6 ms were characterized as bursts.
**b)** Number of spikes when occurring in a burst for *Grin2a*$^{+/+}$ (blue) and *Grin2a*$^{+/-}$ mice (red). Most bursts consist of doublets (two spikes per burst), or triplets (3 spikes per burst) in both genotypes.
**c)** Vulcano plot depicting burst firing frequency alterations per region, set out against region-compared genotype significance. Colours indicate origin area and circle size indicates neuron number.
**d)** Schematic sagittal views for single spike firing frequencies per brain region for genotype effect size, regardless of significance. Selected regions indicated.
**e)** Schematic sagittal views for burst firing frequencies per brain region for genotype effect size, regardless of significance. Note the ventral-dorsal gradient in mutant mice. Selected regions indicated.
**f)** Heatmap depicting effects sizes for burst firing ratio of individual neurons between genotypes, grouped and colored per broadly defined area and alphabetically.
**g)** Vulcano plot depicting burst firing ratio alterations per region, set out against region-compared genotype significance. Colours indicate origin area and circle size indicates neuron number. A subset of regions shows bursting alterations (17 out of 50 regions; ~34%) in mutant mice, of which the majority (11 out of 17; ~65%) shows an increase in bursting behavior.
**h)** Schematic sagittal view for burst firing ratios per brain region for genotype effect size, regardless of significance.
**i)** Swarm plots with burst firing ratios depicting individual neurons for the non-thalamic regions attaining highest pair-wise genotype significance in (f). Black diamonds indicate medians, neuron numbers in brackets, and colors indicate grouping. Y-axes capped at 0.1.
**j)** Schematic view of burst firing ratio results.
Region abbreviations in **Table S3**. Statistics in **Table S7**. ***$p < 0.001$. Two-way ANOVA followed by *post hoc* comparisons.

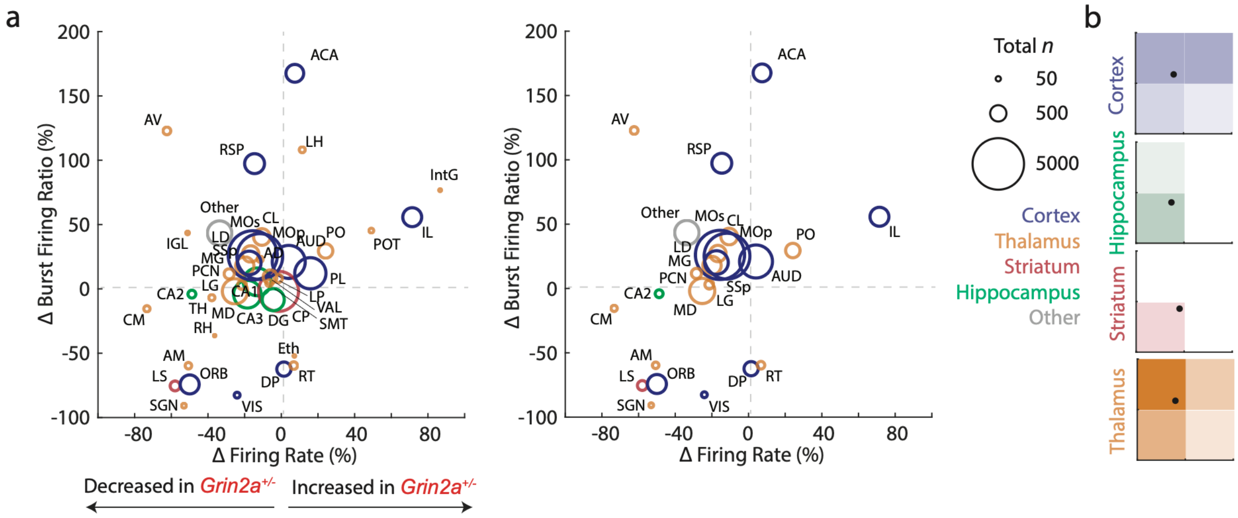

**Fig. S13. Partial overlap between firing rate and burst firing ratio alterations between genotypes.**

**a)** *Left*: Overall firing rate effects set out against burst firing ratio effects for all examined regions, showing a partial overlap between regions. *Right*: As (a), but for top significant affected regions per metric only.

**b)** Relative counts for quadrants shown in (a), with the mean of all regions indicated by black dot, per broadly defined area. Cortex and thalamus show decreased firing rates with increased burst ratios, whereas hippocampus and striatum show decreased firing rates and decreased burst ratios.

Region abbreviations in **Table S3**.

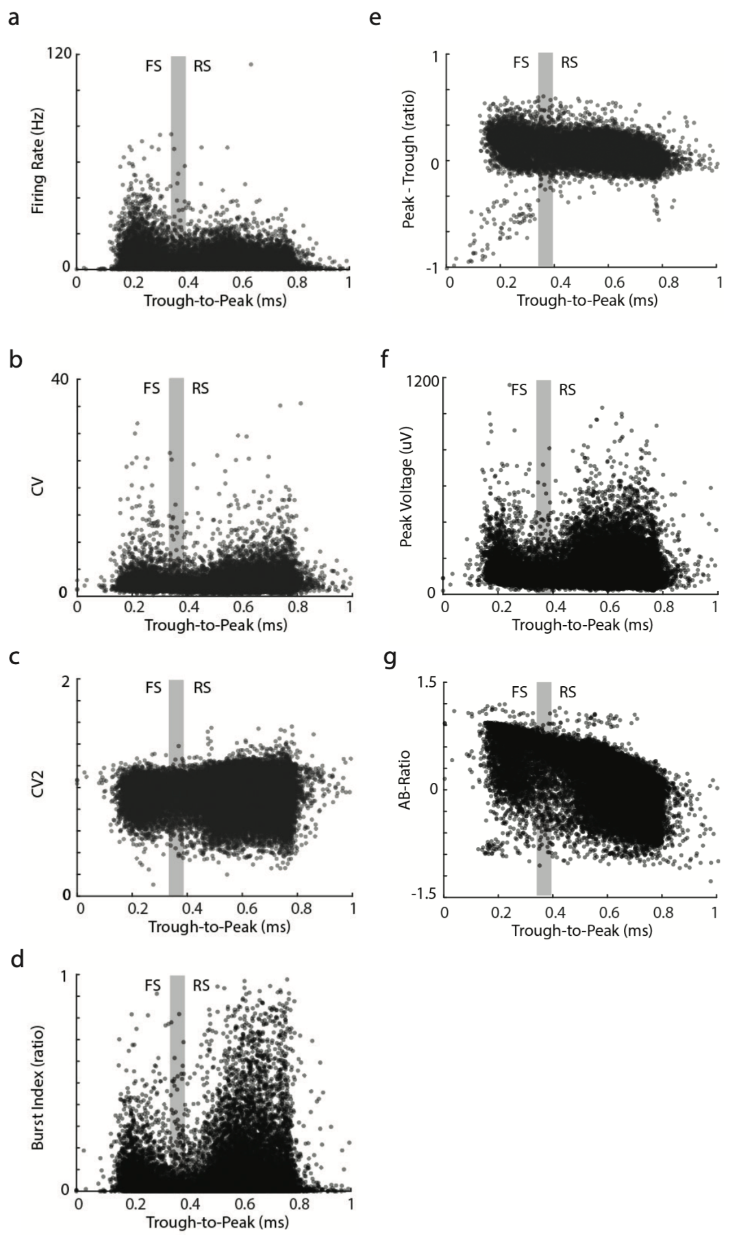

**Fig. S14. Regular-spiking (RS) and fast-spiking (FS) neurons across cortex, hippocampus, and striatum differ across spiking and single-channel waveform characteristics.**

**a)** Scatter plots depicting all individual neurons from cortex, hippocampus and striatum (excluding lateral septum) from both genotypes (*n* = 30,614/16 mice) with their trough-to-peak latency (ms) set out against firing rate (Hz). Gray shaded areas indicate the segregation resulting from bimodal Gaussian fits. RS and FS neurons differ from one another across a range of characteristics.

**b-g)** As (a), but depicting firing rate variability (CV, b; CV2, c), burst firing ratio (d), peak-through ratio (e), peak voltage (f), and shoulder asymmetry (AB-ratio; g).

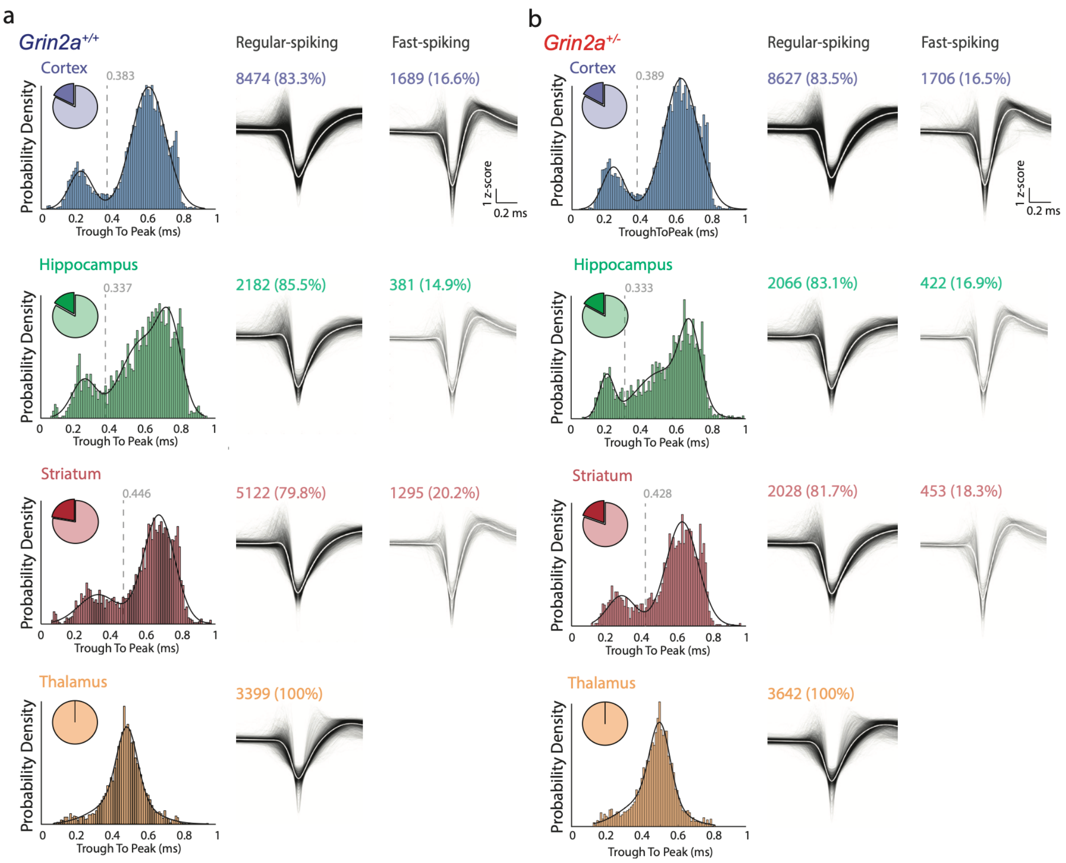

**Fig. S15. Regular-spiking (RS) and fast-spiking (FS) neurons do not differ in overall numbers between both genotypes.**

**a)** *Left*: histogram per broadly defined area showing trough-to-peak latencies obtained from single-channel waveforms for all neurons recorded in that area for *Grin2a*$^{+/+}$ wildtype mice (bars colored as per area) fitted with a bimodal Gaussian (black line), with the valley indicated by the grey dashed line. Insert: pie chart depicting fraction of neurons classified as either regular spiking (RS, light), or fast-spiking (FS, dark). *Right*: Individual average single-channel spike waveforms for RS and FS cells black, mean waveform in white, along with their number and respective percentages within that area. Note that the hippocampus in both genotypes showed a superior fit with a trimodal Gaussian model. Scale bar, 0.2 ms, 1 z-score. Thalamus shows a unimodal distribution of trough-to-peak latencies, in line with a low proportion of fast-spiking putative interneurons.

**b)** As (a), but for *Grin2a*$^{+/-}$ mice.

Statistics in **Table S7**.

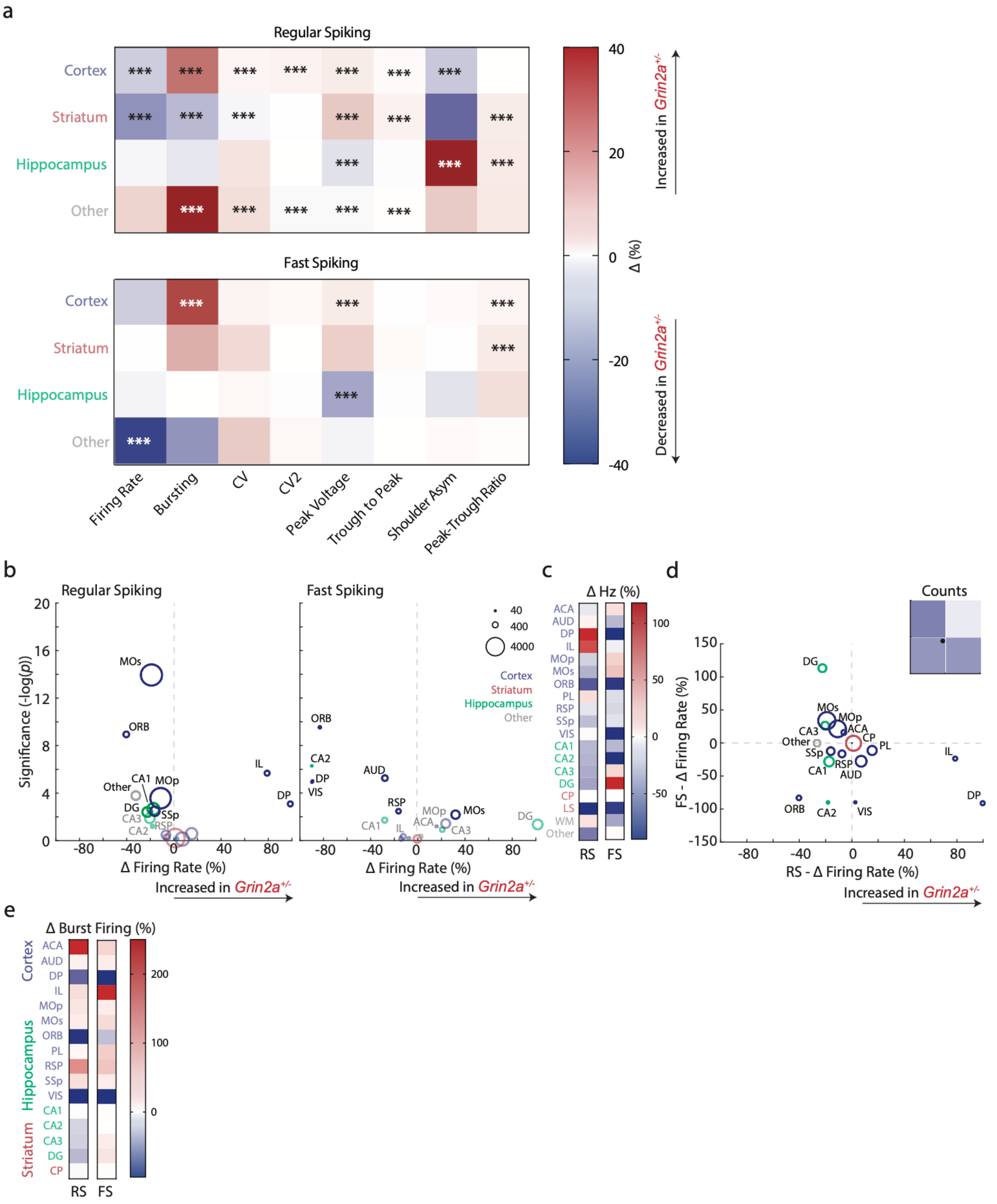

**Fig. S16. *Grin2a*$^{+/-}$ mutant mice display balanced alterations across both fast-spiking and regular-spiking neurons.**

**a)** Heatmap depicting genotype effects for various spiking and single-channel waveform effects between *Grin2a*$^{+/-}$ and *Grin2a*$^{+/+}$ genotypes, segregated between regular-spiking and fast-spiking neurons, over broad-defined areas. Genotype effects for various metrics are observed across both cell subtypes.

**b)** Vulcano plots for regular spiking (left) and fast-spiking neurons (right) depicting firing rate alterations per region between genotypes. Circle sizes indicate total recorded neurons. Colors indicate grouping per higher-order area. Greyed out regions did not attain Bonferroni-corrected threshold for significance.
**c)** Heatmaps depicting the data in (b), ordered per broadly defined area and alphabetically.
**d)** Scatter plot depicting effect sizes for regular-spiking and fast-spiking neurons between genotypes, regardless of significance. An inverse relationship can be observed between effects sizes for regular-spiking and fast-spiking neurons between genotypes. *Insert*: Relative counts per quadrant indicating regions either depict regular-spiking rate reduction, fast-spiking rate reduction, or both.
**e)** Burst firing ratio differences for regular-spiking (RS) and fast-spiking (FS) neurons per region, ordered per broadly defined area and alphabetically. Burst firing ratio alterations are observed across both populations.
*** $p < 0.001$. One-way ANOVA followed by *post hoc* comparison per parameter and cell class. Region abbreviations in **Table S3**. Statistics in **Table S7**.

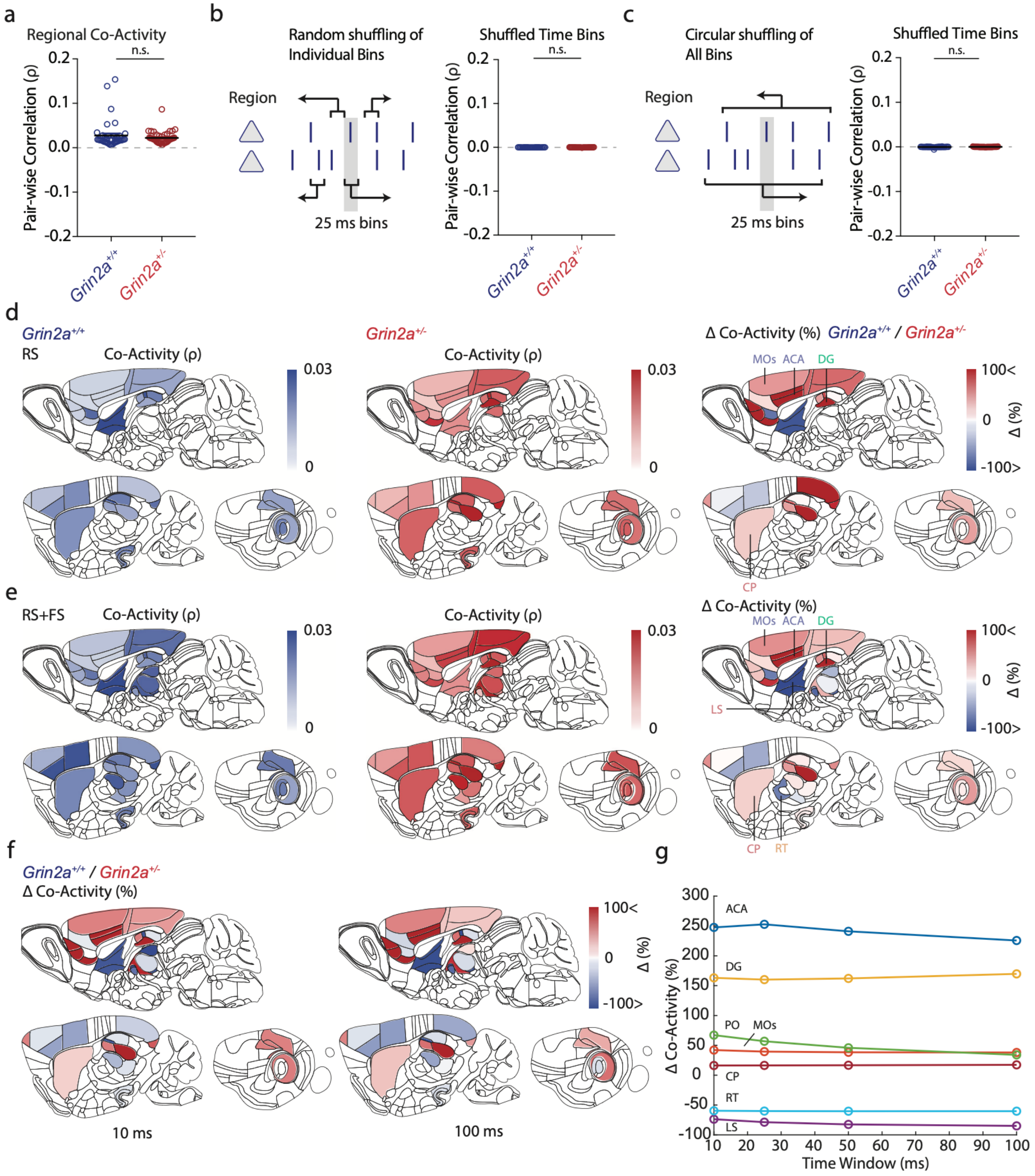

660 **Fig. S17. Co-activity alterations in *Grin2a*$^{+/-}$ mice are invariant to time window or cell-type.**

**a)** Mean regional network co-activity for *Grin2a*$^{+/-}$ and *Grin2a*$^{+/+}$ mice for 25-ms time bins. Each circle indicates one value for all neurons in that region, black lines indicate median across regions.

**b)** Random shuffling of spike bins negates pair-wise correlations within regions between genotypes. black lines indicate median across regions.

665 **c)** Circular shuffling of spike bins negates pair-wise correlations within regions between genotypes. black lines indicate median across regions.

**d)** Sagittal brain views for mean co-activity values across the brain for wildtype *Grin2a*$^{+/+}$ mice (left), mutant *Grin2a*$^{+/-}$ mice (middle), and their difference (right), for regular-spiking neurons. Selected regions indicated.

**e)** As (d), but for all neurons per region.

**f)** Sagittal view of genotype effect sizes for regular-spiking co-activity across two different time windows (10 ms – 100 ms).

**g**) Effect of network co-activity values with various analysis time windows between genotypes, showing minimal effects of time window duration.

n.s. non-significant. Unpaired two-tailed Student’s t-test in (a) through (c).

Region abbreviations in **Table S3**. Statistics in **Table S7**.

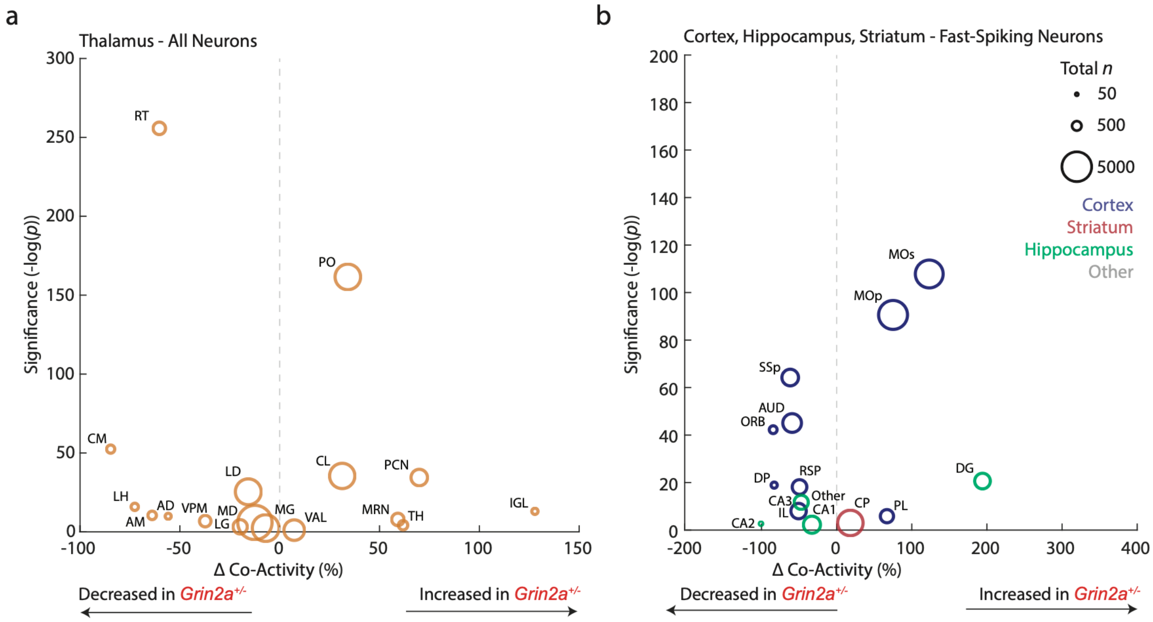

**Fig. S18. Bidirectional co-activity alterations in *Grin2a*$^{+/-}$ mice in thalamus and fast-spiking neurons.**

**a)** Vulcano plot depicting effects size for neuronal co-activity of all neurons between genotypes, restricted to thalamic nuclei, set out against pair-wise genotype significance. Circle size indicates neuron number.
**b)** As (a), but restricted to fast-spiking neurons in cortex, striatum, and hippocampus. Circle colour indicates grouped area and size indicates neuron number.
Region abbreviations in **Table S3**. Statistics in **Table S7**.

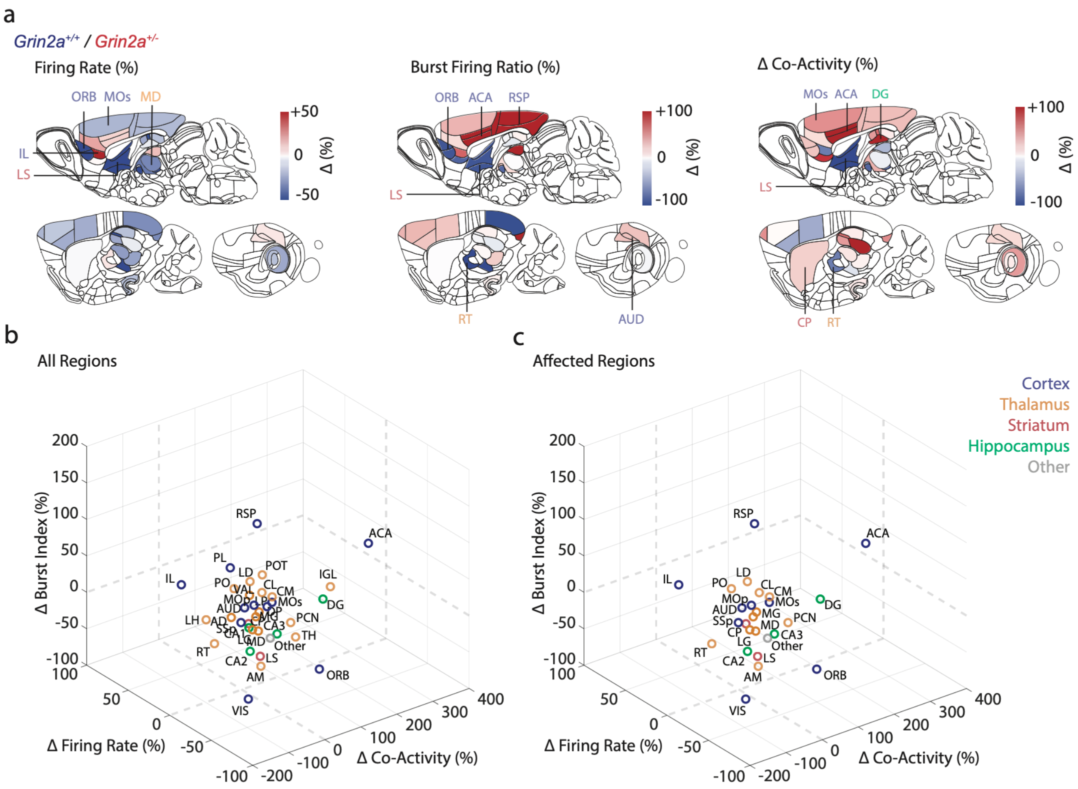

**Fig. S19. Co-activity alterations in *Grin2a*$^{+/-}$ mutants are independent to firing rate or burst firing.**
**a)** Schematic sagittal visualizations of effect sizes for genotype firing rate differences (left), burst firing (middle) differences, and co-activity (right) differences between genotypes.
**b)** 3d scatter plot displaying the partial overlap of rate, bursting, and co-activity of all regions, for the comparison of *Grin2a*$^{+/+}$ and *Grin2a*$^{+/-}$.
**c)** 3d scatter plot displaying the partial overlap of rate, bursting, and co-activity of affected regions, *Grin2a*$^{+/+}$ and *Grin2a*$^{+/-}$ under enhancement of higher-order thalamus.
Region abbreviations in **Table S3**.

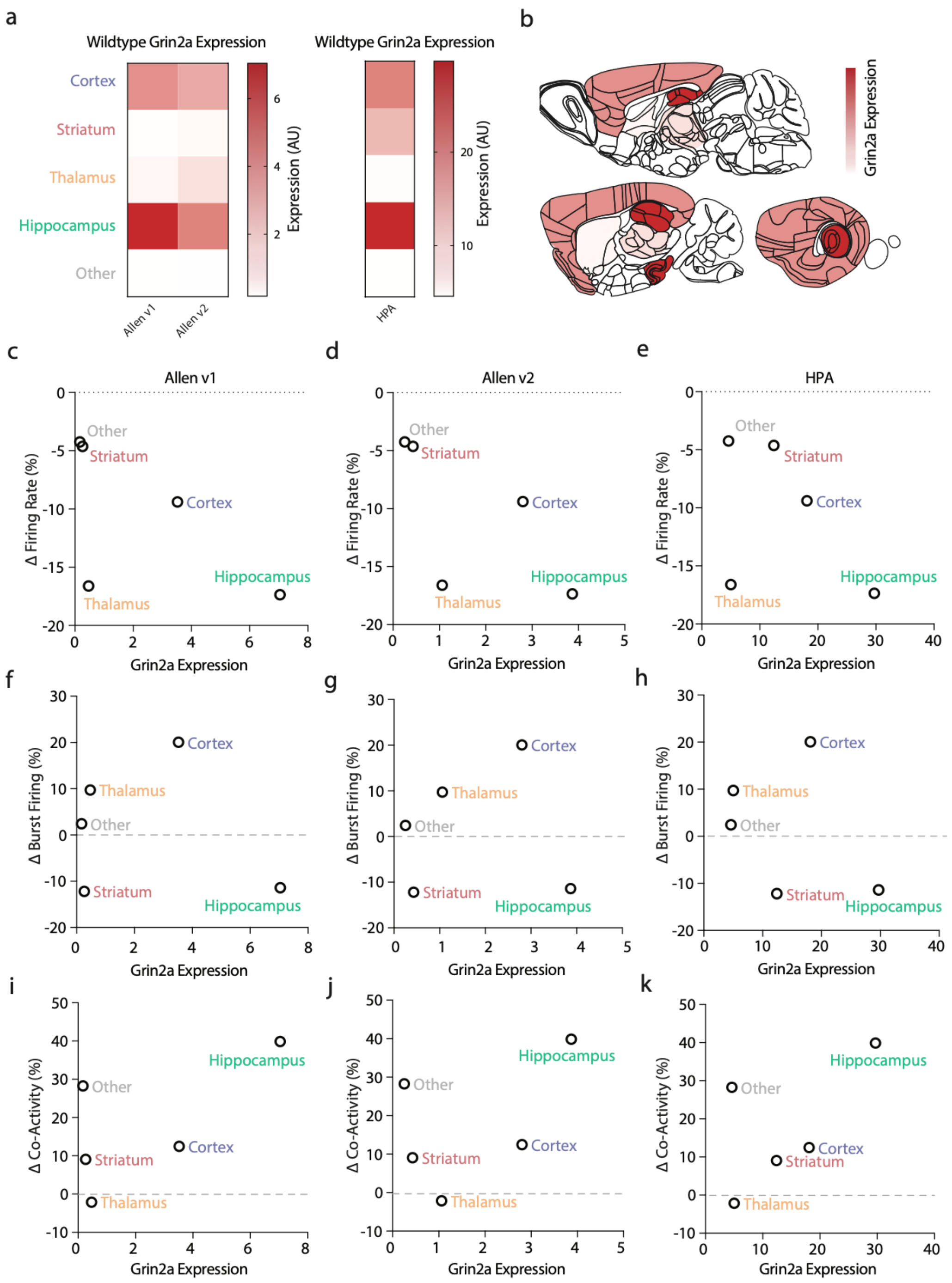

**Fig. S20. Correlations between physiology and bulk *Grin2a* expression in wildtype animals.**
**a)** Bulk RNA expression obtained from (left) two individual experiments from the Allen Brain Atlas and from the Human Protein Atlas (source: **Materials and Methods**) for Grin2a expression in rodent. Expression was consistently highest in hippocampus.
**b)** Schematic sagittal view of expression of Grin2a across grouped areas.
**c-e)** Grin2a expression levels in (a-b) and firing rate effect sizes between genotypes observed in Fig. 2. Thalamus appears low in Grin2a expression across all databases (c-e) but show large firing rate effect sizes.

**f-h)** Grin2a expression levels in (a-b) and burst firing effect sizes between genotypes observed in Fig. S12. Cortex appears intermediate in Grin2a expression across all databases (f-h) but shows large increases in burst firing ratio effect sizes.

**i-k)** Grin2a expression levels in (a-b) and network co-activity effect sizes between genotypes observed in Fig. 3. Hippocampus appears high in Grin2a expression across all databases (i-k) and shows large increases in co-activity.

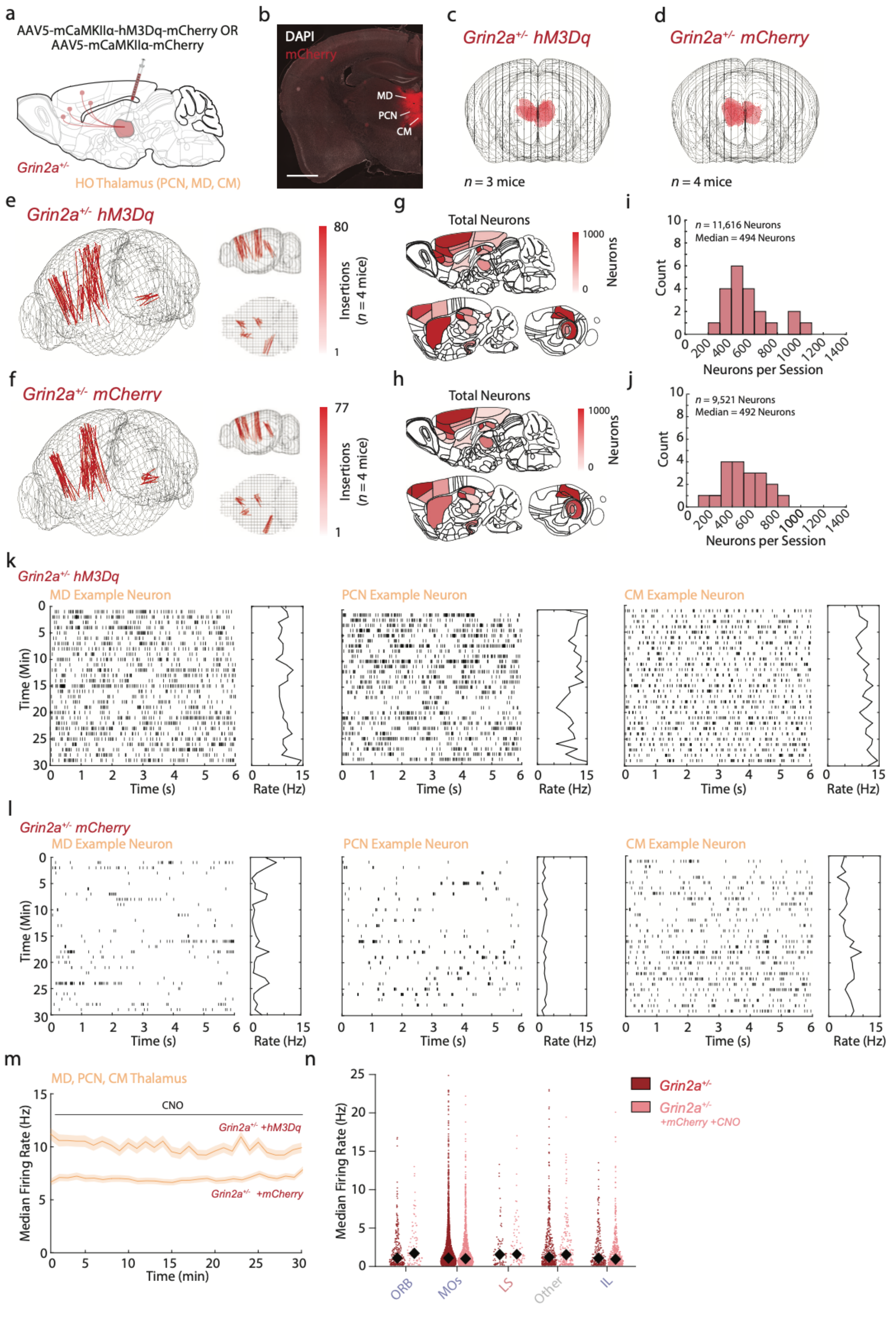

**Fig. S21. Acute and stable chemogenetic enhancement of higher-order thalamus in mutant *Grin2a*$^{+/-}$ mice**

**a)** Schematic sagittal view of viral transduction approach. Adult mutant *Grin2a*$^{+/-}$ mice received stereotactic intracranial injections of AAV5 containing hM3Dq-mCherry or mCherry fluorescent control in bilateral higher-order thalamus targeting MD, CM, and PCN.

**b)** Representative epifluorescent coronal microscopic image of mCherry labeling (red) in higher-order thalamus in a mutant *Grin2a*$^{+/-}$ mice. MD, PCN, MD indicated. DAPI nuclear marker in white. Scale bar, 1 mm.

**c-d)** Front view of 3d schematic of brains depicting reconstructions of viral transduction spread across all animals for *Grin2a*$^{+/-}$ hM3Dq-mCherry (c) and *Grin2a*$^{+/-}$ mCherry (d) animals. Red areas indicate viral transduction. For one *Grin2a*$^{+/-}$ hM3Dq-mCherry we could not recover viral labeling, and this animal was excluded from further analysis.

**e-f)** 3d schematic of brains showing reconstructed probe trajectories for all *Grin2a*$^{+/-}$ hM3Dq-mCherry (e) and *Grin2a*$^{+/-}$ mCherry control (f) animals. All animals and recording tracks shown.

**g-h)** Sagittal view of mouse brain depicting the number of neurons per brain region for both manipulation types across all animals.

**i-j)** Histograms depicting the total number of neurons per session across all animals.

**k)** Exemplar single neuron activity across time along with minute-to-minute mean rate, suggesting stable enhancement of firing rate, for example neurons in MD (left), PCN (middle), and CM (right) nuclei of the thalamus, in *Grin2a*$^{+/-}$ animals with hM3Dq-mCherry after CNO application.

**l)** As (k), but in *Grin2a*$^{+/-}$ mutant control animals with mCherry control after CNO application.

**m)** Median firing rate activity across all neurons over time in higher-order thalamus (MD, PCN, CM; $n$ = 506 neurons) over time for *Grin2a*$^{+/-}$ animals with hM3Dq-mCherry and *Grin2a*$^{+/-}$ animals with mCherry after CNO application (1 mg/kg i.p.). Shaded areas indicate s.e.m.

**n)** Swarmplot depicting spontaneous firing rates between untreated *Grin2a*$^{+/-}$ mutant animals (dark red; Fig 1-3) and *Grin2a*$^{+/-}$ mutant animals (light red) under application of CNO, for the main affected regions. Expression of mCherry fluorescent protein in thalamus and i.p. application of CNO have minimal effect. Dots represent individual neurons, black diamonds indicate median values.

Region abbreviations in **Table S3**. Statistics in **Table S7**.

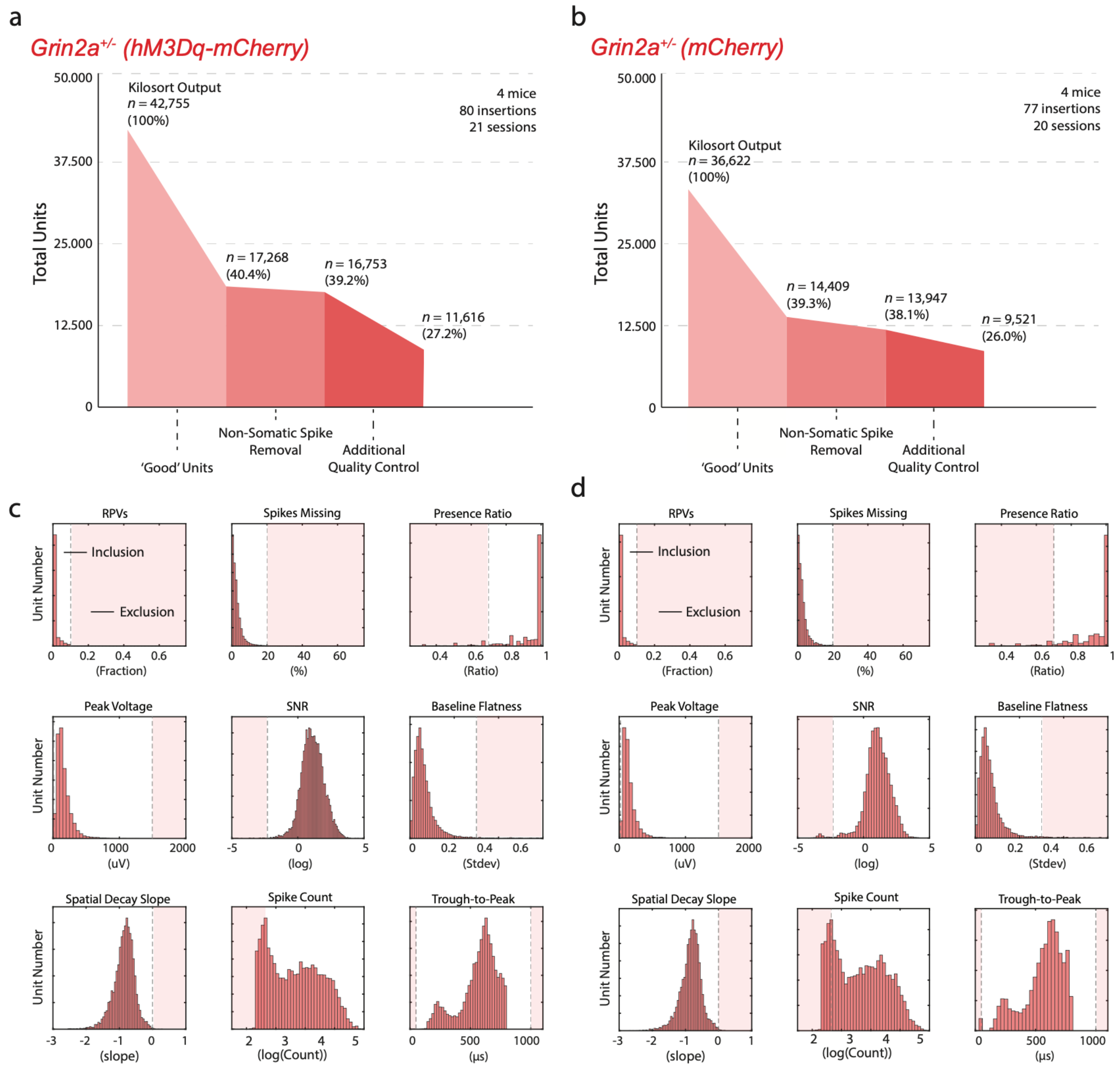

**Fig. S22. Recorded neuron quality control metrics are not affected by mCherry expression or CNO application**

**a-b)** Flowchart showing quality control steps and neuron exclusion based on successive criteria in both control *Grin2a*$^{+/-}$ with hM3Dq-mCherry (a) and *Grin2a*$^{+/-}$ with mCherry control (b) mutant mice, for all animals.

**c-d)** Histograms of several quality control characteristics in control *Grin2a*$^{+/+}$ (c) and *Grin2a*$^{+/-}$ (d) mutant mice before the quality control step (*n* = 17,268; *n* = 14,409, respectively), along with exclusion criteria (vertical grey dashed lines) and to be further excluded neurons (red shaded areas).

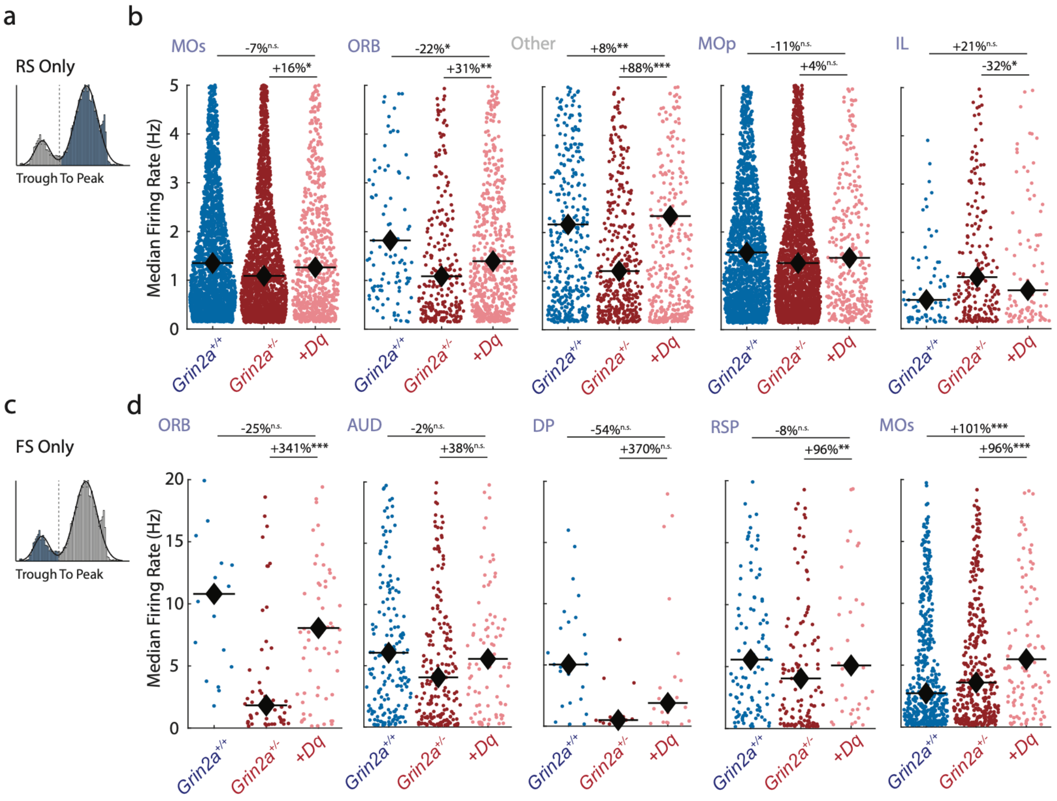

**Fig. S23. Restoration of higher-order thalamus activity increases RS and FS firing rates across affected cortical regions.**
**a,c)** Schematic depiction of through-to-peak latency as a segregation between regular-spiking neurons and fast-spiking neurons.
**b,d)** Swarmplots depicting the effect of thalamus activity enhancement in untreated *Grin2a*$^{+/+}$ animals (blue), *Grin2a*$^{+/-}$ animals (dark red), and *Grin2a*$^{+/-}$ mutant with hM3Dq-mCherry under CNO application (light red) on regular-spiking (RS, b) and fast-spiking neurons (FS, d) in most prominently altered regions. Black diamonds indicate medians. Thalamus activity restoration effects on cortical and other regions show trends in increasing activity across both RS and FS subclasses.
Statistics indicate one-way ANOVA (not shown) followed by *post hoc* comparison between wildtype and *Grin2a*$^{+/-}$ mutant with hM3Dq-mCherry and CNO application, and between untreated *Grin2a*$^{+/-}$ mutants and *Grin2a*$^{+/-}$ mutant with hM3Dq-mCherry under CNO application.
Region abbreviations in **Table S3**. Statistics in **Table S7**.

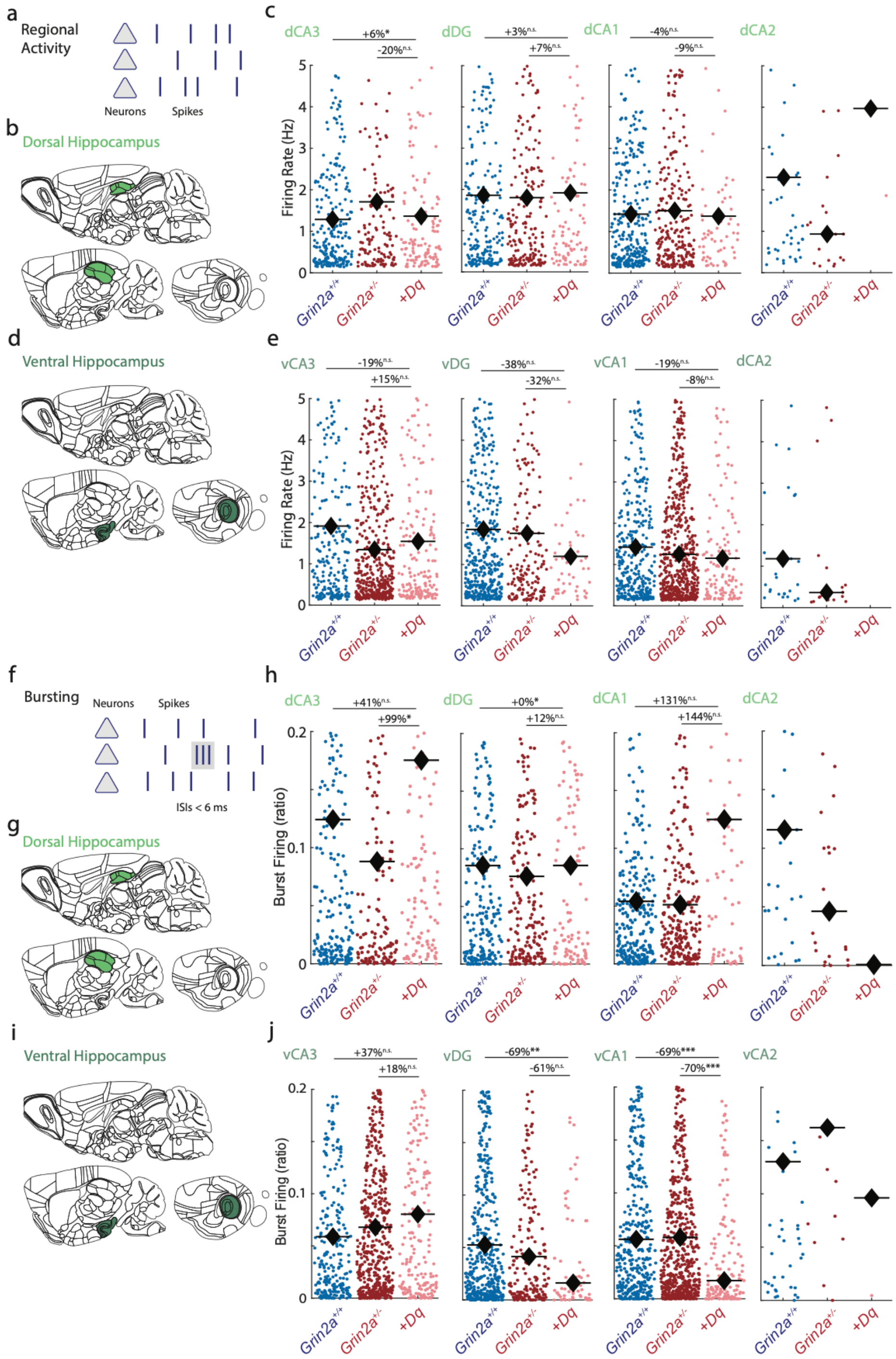

**Fig. S24. Restoration of higher-order thalamus activity shows trends toward correcting activity and burst firing in dorsal, but not ventral hippocampus**

**a-b,d)** Schematic depiction of differentiation of analysis of firing rates (a) between dorsal (b) and ventral (d) hippocampus.

**c,e)** Firing rate alterations for dorsal (c) and ventral (d) hippocampus in untreated wildtype *Grin2a*$^{+/+}$ mice (blue), untreated *Grin2a*+/- mutant mice (dark red), and higher-order thalamus restored *Grin2a*$^{+/-}$ mutant

mice (light red). Black diamonds indicate median values. Thalamus activity restoration shows a trend toward decreasing firing activity in dorsal CA3, but not ventral CA3. Y-axis capped at 5 Hz for visibility.
**f,g,i)** Schematic depiction of differentiation of analysis of burst firing (f) between dorsal (g) and ventral (i) hippocampus.
**h-j)** Burst firing alterations for dorsal (h) and ventral (j) hippocampus in untreated wildtype *Grin2a*$^{+/+}$ mice (blue), untreated *Grin2a*+/- mutant mice (dark red), and higher-order thalamus enhanced *Grin2a*$^{+/-}$ mutant mice (light red). Black diamonds indicate median values. Thalamus activity restoration shows an effect of increasing burst firing in dorsal CA3, but not ventral CA3. Y-axis capped at 0.2 for visibility.
***$p < 0.001$, **$p < 0.01$, **$p < 0.05$. n.s. non-significant. Two-way ANOVA (not shown) followed by *post hoc* comparison between wildtype and *Grin2a*$^{+/-}$ mutant with hM3Dq-mCherry and CNO application, and between untreated *Grin2a*$^{+/-}$ mutants and *Grin2a*$^{+/-}$ mutant with hM3Dq-mCherry under CNO application.
Region abbreviations in **Table S3**. Statistics in **Table S7**.

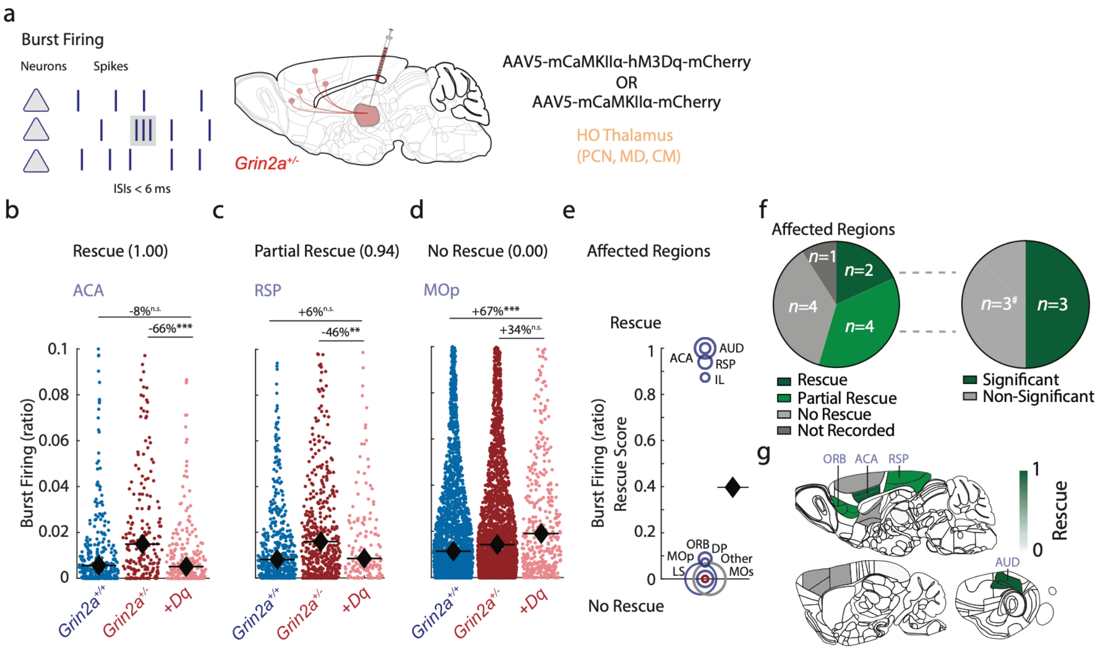

**Fig. S25. Restoration of higher-order thalamus activity normalizes burst firing increases in *Grin2a*$^{+/-}$ mutant mice in prefrontal cortex and primary auditory cortex.**

**a)** Burst firing (ratio) was quantified by the number of action potentials in quick succession (interspike interval <6 ms) divided by the number of total spikes per neuron. Schematic sagittal view of viral transduction approach. Mutant *Grin2a*$^{+/-}$ mice received bilateral stereotactic injections of AAV5 containing hM3Dq-mCherry or mCherry fluorescent control in bilateral higher-order thalamus targeting MD, CM, and PCN.

**b-d)** Burst firing rates for *Grin2a*$^{+/+}$ (blue), *Grin2a*$^{+/-}$ (dark red), and *Grin2a*$^{+/-}$ hM3Dq-mCherry under CNO (light red) for three regions with different rescue scores: ACA (b), RSP (c), MOp (d). Each dot represents an individual neuron, black diamonds indicate medians. Y-axis capped at 0.1 for visibility.

**e)** Scatter plot depicting rescue scores across the 10 significantly affected non-thalamic regions from Fig. S12. Each circle indicates a region, circle size indicates neuron number. Several regions show full rescue of spontaneous burst firing rates. Black diamond indicates the mean rescue score.

**f)** Pie chart showing the fraction of regions showing no, partial, or full rescue (left), along with the fraction of partial and fully rescue regions attaining single-region significance (right).

**g)** Schematic sagittal view depicting rescue scores for burst firing across affected regions.

***$p < 0.001$, n.s. non-significant. One-way ANOVA followed by *post hoc* comparisons in (b) through (d) and (f).

Region abbreviations in **Table S3**. Statistics in **Table S7**.

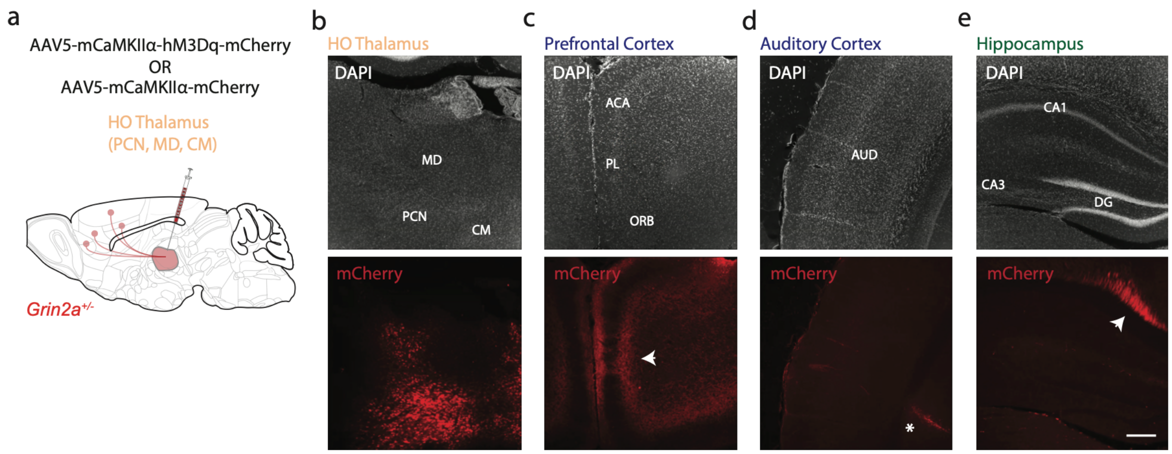

**Fig. S26. Absence of higher-order thalamus derived mCherry+ projections in hippocampus or primary sensory cortex**

**a)** Schematic sagittal view of viral approach. Mutant *Grin2a*$^{+/-}$ mice received bilateral stereotactic injection of AAV5 containing hM3Dq-mCherry or mCherry fluorophore control in bilateral higher-order thalamus targeting MD, CM, and PCN.

**b-e)** Representative epifluorescent coronal images of DAPI nuclear marker (white) and viral mCherry (red) showing transduced neuronal somata in thalamus (b) and mCherry+ axons in prefrontal cortex (arrowhead, c), but not in primary auditory cortex (d), and hippocampus (e). Important region abbreviations indicated. Note the labeled white matter tracts towards anterior cingulate cortex (arrowhead, e), and remnant DiI+ probe tract in (red line in d; asterisk). Scale bar, 200 µm.
Region abbreviations in **Table S3**.

| **Class** | **Subclass** | ***Grin2a***$^{+/+}$ | ***Grin2a***$^{+/-}$ | **Statistic** |
|---|---|---|---|---|
| Overall | Animals | 8 | 8 | NA |
| | Sessions | 38 | 38 | NA |
| | Insertions | 119 | 125 | NA |
| | Insertions/Session | ~3.13 | ~3.29 | NA |
| Behavior | Pre-operative weight (g) | 28.4 ± 0.6 | 29.0 ± 0.7 | t=0.65, df=14, $p$ = 0.526 |
| | Weight at Recordings (%) | 88.7 ± 0.3 | 88.6 ± 0.2 | t=0.4671 df=78 $p$ = 0.641 |
| | Total Training Sessions | 11.75 | 11.75 | t=0.001, df=14, $p$ = 1 |
| | Total Training Trials | 5067 ± 674 | 5063 ± 509 | t=0.006, df=14, $p$ = 0.996 |
| Recording | Head-fixation Duration (mins) | 86.8 ± 0.9 | 88.8 ± 0.8 | t=1.598, df=91, $p$ = 0.114 |
| | Total Recording Duration (s) | 2024 ± 24 | 2082 ± 17 | t=1.901, df=74, $p$ = 0.061 |
| Histology | Probe Points | 26.5 ± 0.8 | 27.2 ± 0.8 | t=0.639, df=277, $p$ = 0.524 |
| | Electrophysiological Landmarks | 2.6 ± 0.1 | 2.6 ± 0.1 | t=0.2074, df=257, $p$ = 0.836 |

**Table S1. Experimental comparison between genotypes**

Values are mean ± s.e.m. Statistics represents two-tailed unpaired Student's *t*-tests.

| Region | Landmarks | Feature | Reference |
|---|---|---|---|
| All | Brain surface<br><br>White matter tract<br><br>Ventricles | No spiking<br>Reduced LFP power<br>Lower spiking<br>Smaller spike amplitude<br>No spiking<br>No neurons<br>Reduced LFP power | [7,9,10,66] |
| Thalamus | Boundary from hippocampus | Higher spiking<br>Larger spike amplitude | [7,9,10,66] |
| Hypothalamus | Boundary from thalamus | Smaller spike amplitude | [7,9,10,66] |
| Hippocampus | CA1 pyramidal cell layer<br>Dentate gyrus | LFP phase reversal<br>High neuron density<br>Peak LFP power | [7,9,10,66] |
| Striatum | Boundary from cortex | Lower spiking<br>Smaller spike amplitude | [7,9,10,66] |

**Table S2. Electrophysiological landmarks**

Previously validated electrophysiological landmarks used for realignment of channels along the long axis of the probe as well as the publication reference.

| **Area** | **Region** | **Included** | **Name** |
|---|---|---|---|
| Cortex | ACA | ACAv, ACAd | Anterior cingulate area |
| Thalamus | AD | AD | Anterodorsal thalamus |
| Thalamus | AM | AMd, AMv | Anteromedial thalamus |
| Cortex | AUD | AUDv, AUDd, AUDp, AUDpo | Auditory cortex |
| Thalamus | AV | AV | Anteroventral thalamus |
| Hippocampus | CA1 | CA1 | Hippocampal CA1 area |
| Hippocampus | CA2 | CA2 | Hippocampal CA2 area |
| Hippocampus | CA3 | CA3 | Hippocampal CA3 area |
| Thalamus | CL | CL | Central lateral nucleus of the thalamus |
| Thalamus | CM | CM | Central median nucleus of the thalamus |
| Striatum | CP | CP | Caudate-putamen |
| Hippocampus | DG | DG_mo, DG_po, DG_sg | Dentate gyrus |
| Cortex | DP | DP | Dorsal peduncle |
| Thalamus | Eth | Eth | Ethmoid nucleus |
| Thalamus | IGL | IGL | Intergeniculate leaflet of the LGN |
| Cortex | IL | ILA | Infralimbic area |
| Thalamus | IntG | IntG | Intermediate geniculate nucleus |
| Thalamus | LD | LD | Laterodorsal thalamus |
| Thalamus | LG | LGd_sh, LGv, LGd_sho | Lateral geniculate thalamus |
| Thalamus | LH | LH | Lateral habenula |
| Thalamus | LP | LP | Lateroposterior nucleus of the thalamus |
| Striatum | LS | LSc, LSr | Lateral septum |
| Thalamus | MD | MD, IMD | Mediodorsal thalamus |
| Thalamus | MG | MGBd, MGBv, MGBd | Medial geniculate thalamus |
| Cortex | MOp | MOp | Primary motor cortex |
| Cortex | MOs | MOs | Secondary motor cortex |
| Cortex | ORB | ORBl, ORBm, ORBvl | Orbitofrontal cortex |
| Thalamus | PCN | PCN | Paracentral nucleus of the thalamus |
| Cortex | PL | PL | Prelimbic area |
| Thalamus | PO | PO | Posterior nucleus of the thalamus |
| Thalamus | POL | POL | Posterior limiting nucleus of thalamus |
| Thalamus | POT | POT | Posterior thalamus |
| Cortex | RSP | RSPd, RSPv, RSPagl | Retrosplenial cortex |
| Thalamus | RH | RH | Rhomboid nucleus of the thalamus |
| Thalamus | RT | RT | Reticular nucleus of the thalamus |
| Thalamus | SGN | SGN | Subgenual nucleus of the thalamus |
| Thalamus | SMT | SMT | Submedial nucleus of the thalamus |
| Cortex | SSp | SSP_ll, SSP_tr | Primary somatosensory cortex |
| Striatum | STR | STR | Striatum, *undefined* |
| Thalamus | TH | TH | Thalamus, *undefined* |
| Thalamus | VAL | VAL | Ventral anteriolateral nucleus of the thalamus |
| Cortex | VIS | VISa | Visual cortex |
| Thalamus | VPM | VPM | Ventral posterior nucleus of the thalamus |
| Other | WM | alv, ar, ccb, ccg, ccs, chpl, cing, fa, fi, fiber_tracts, fp, or, out, root, scwm,sm | *White matter regions* |
| Other | Other | SEZ, MB, MRN, OLF, TTd, SAG, MRN, HY | *Unclustered regions* |

**Table S3. Region names and clustering**

Regions parcellated as per Allen Brain Institute Reference Atlas (CCFv3) [50], sorted alphabetically. All layers were combined for cortical regions (e.g. 1, 23, 4, 5, 6) or hippocampus. Only regions that were recorded from in both genotypes included in this list.

| | | *Grin2a*$^{+/+}$ | | | *Grin2a*$^{+/-}$ | | |
|---|---|---|---|---|---|---|---|
| **Area** | **Region** | **Neurons** | **Sessions** | **Mice** | **Neurons** | **Sessions** | **Mice** |
| Cortex | ACA | 297 | 19 | 8 | 197 | 14 | 7 |
| Thalamus | AD | 16 | 2 | 2 | 26 | 2 | 2 |
| Thalamus | AM | 46 | 1 | 1 | 38 | 4 | 3 |
| Cortex | AUD | 1309 | 27 | 7 | 1201 | 31 | 8 |
| Thalamus | AV | 14 | 3 | 2 | 118 | 7 | 4 |
| Hippocampus | CA1 | 945 | 28 | 8 | 1032 | 28 | 8 |
| Hippocampus | CA2 | 97 | 10 | 6 | 48 | 8 | 7 |
| Hippocampus | CA3 | 607 | 26 | 8 | 809 | 26 | 8 |
| Thalamus | CL | 305 | 9 | 5 | 514 | 9 | 5 |
| Thalamus | CM | 28 | 3 | 3 | 38 | 3 | 2 |
| Striatum | CP | 2455 | 27 | 8 | 2335 | 28 | 8 |
| Hippocampus | DG | 914 | 23 | 8 | 491 | 28 | 8 |
| Cortex | DP | 141 | 4 | 2 | 81 | 6 | 5 |
| Thalamus | Eth | 10 | 2 | 2 | 14 | 1 | 1 |
| Thalamus | IGL | 22 | 2 | 1 | 23 | 2 | 1 |
| Cortex | IL | 84 | 8 | 4 | 286 | 17 | 8 |
| Thalamus | IntG | 15 | 2 | 1 | 13 | 1 | 1 |
| Thalamus | LD | 447 | 12 | 6 | 342 | 12 | 6 |
| Thalamus | LG | 134 | 6 | 2 | 106 | 5 | 1 |
| Thalamus | LH | 54 | 5 | 5 | 32 | 6 | 5 |
| Thalamus | LP | 105 | 6 | 3 | 121 | 7 | 3 |
| Striatum | LS | 130 | 4 | 4 | 70 | 1 | 1 |
| Thalamus | MD | 799 | 11 | 5 | 636 | 9 | 6 |
| Cortex | MG | 259 | 15 | 6 | 583 | 19 | 5 |
| Cortex | MOp | 2767 | 26 | 7 | 3318 | 26 | 7 |
| Cortex | MOs | 3385 | 32 | 8 | 2848 | 29 | 8 |
| Cortex | ORB | 133 | 4 | 1 | 320 | 5 | 3 |
| Thalamus | PCN | 173 | 8 | 5 | 134 | 7 | 5 |
| Cortex | PL | 859 | 21 | 8 | 833 | 17 | 8 |
| Thalamus | PO | 476 | 8 | 5 | 291 | 11 | 5 |
| Thalamus | POL | 20 | 2 | 2 | 51 | 5 | 2 |
| Thalamus | POT | 34 | 3 | 2 | 47 | 5 | 2 |
| Thalamus | RH | 4 | 1 | 1 | 16 | 2 | 2 |
| Thalamus | RSP | 509 | 15 | 8 | 564 | 10 | 5 |
| Thalamus | RT | 57 | 2 | 2 | 125 | 2 | 1 |
| Thalamus | SGN | 72 | 4 | 3 | 8 | 1 | 1 |
| Thalamus | SMT | 62 | 2 | 1 | 52 | 1 | 1 |
| Cortex | SSp | 714 | 13 | 7 | 693 | 21 | 8 |
| Striatum | STR | 35 | 3 | 3 | 40 | 10 | 6 |
| Thalamus | TH | 36 | 8 | 4 | 60 | 12 | 6 |
| Thalamus | VAL | 277 | 8 | 5 | 221 | 4 | 4 |
| Cortex | VIS | 84 | 2 | 2 | 37 | 1 | 1 |
| Thalamus | VPM | 4 | 1 | 1 | 140 | 4 | 3 |
| Other | WM | 1694 | 38 | 8 | 1930 | 38 | 8 |
| Other | Other | 624 | 22 | 8 | 654 | 27 | 8 |

**Table S4. Neurons per region across both genotypes**

Number of neurons per region across both genotypes. Only regions that were recorded from in both genotypes included in this list. Region abbreviations in **Table S3**.

| Class | Subclass | *Grin2a*$^{+/-}$ Dq | *Grin2a*$^{+/-}$ mCh | Statistic |
|---|---|---|---|---|
| Overall | Animals | 4 | 4 | NA |
| | Sessions | 21 | 20 | NA |
| | Insertions | 80 | 77 | NA |
| | Insertions/Session | 3.8 | 3.9 | NA |
| Behavior | Pre-operative weight (g) | 26.3 ± 1.7 | 27.2 ± 1.0 | t=0.436, df=6, $p$ = 0.679 |
| | Weight at Recordings (%) | 89.1 ± 0.6 | 88.7 ± 0.4 | t=0.632 df=44 $p$ = 0.341 |
| | Total Training Sessions | 11.5 | 11.5 | t=0.001, df=6, $p$ = 1 |
| | Total Training Trials | 2756 ± 523 | 4575 ± 497 | t=2.520, df=6, **$p$ = 0.0453*** |
| Recording | Head-fixation Duration (mins) | 90.1 ± 1.3 | 87.5 ± 1.5 | t=1.304, df=39, $p$ = 0.199 |
| | Total Recording Duration (s) | 2162 ± 40 | 2122 ± 41 | t=0.685, df=38, $p$ = 0.497 |
| | CNO Injection Time till Recording (min) | 44.1 ± 1.0 | 43.2 ± 1.2 | t = 0.568, df=39, $p$ = 0.574 |
| Histology | Probe Points | 22.9 ± 0.9 | 22.0 ± 0.7 | t=0.851, df=146, $p$ = 0.397 |
| | Electrophysiological Landmarks | 2.2 ± 0.1 | 2.4 ± 0.1 | t=1.56, df=141, $p$ = 0.121 |

**Table S5. Experimental comparisons between manipulation experiments**

Values indicate means ± s.e.m. Statistics represent two-tailed unpaired Student’s *t*-tests.

| | | *Grin2a*$^{+/-}$ *hM3Dq-mCherry* | | | *Grin2a*$^{+/-}$ *mCherry* | | |
|---|---|---|---|---|---|---|---|
| **Area** | **Region** | **Neurons** | **Sessions** | **Mice** | **Neurons** | **Sessions** | **Mice** |
| Cortex | ACA | 429 | 13 | 4 | 317 | 15 | 4 |
| Thalamus | AD | 38 | 4 | 3 | 63 | 4 | 2 |
| Thalamus | AM | 36 | 4 | 3 | 46 | 2 | 1 |
| Cortex | AUD | 859 | 17 | 4 | 890 | 17 | 4 |
| Thalamus | AV | 96 | 4 | 3 | 333 | 2 | 2 |
| Hippocampus | CA1 | 391 | 16 | 4 | 483 | 16 | 4 |
| Hippocampus | CA2 | 27 | 6 | 2 | 47 | 9 | 3 |
| Hippocampus | CA3 | 829 | 18 | 4 | 690 | 19 | 4 |
| Thalamus | CL | 102 | 4 | 2 | 74 | 7 | 4 |
| Thalamus | CM | 43 | 3 | 2 | 29 | 3 | 2 |
| Striatum | CP | 651 | 17 | 4 | 522 | 14 | 4 |
| Hippocampus | DG | 284 | 15 | 4 | 136 | 16 | 4 |
| Cortex | DP | 412 | 13 | 4 | 147 | 8 | 4 |
| Cortex | IL | 390 | 11 | 3 | 551 | 12 | 4 |
| Thalamus | LD | 89 | 3 | 2 | 86 | 6 | 3 |
| Thalamus | LH | 22 | 2 | 2 | 18 | 3 | 3 |
| Thalamus | LP | 2 | 1 | 1 | 21 | 3 | 2 |
| Striatum | LS | 86 | 2 | 2 | 74 | 4 | 3 |
| Thalamus | MD | 394 | 7 | 3 | 473 | 11 | 4 |
| Thalamus | MG | 181 | 7 | 4 | 191 | 7 | 4 |
| Cortex | MOp | 693 | 15 | 4 | 883 | 16 | 4 |
| Cortex | MOs | 1659 | 21 | 4 | 1263 | 18 | 4 |
| Cortex | ORB | 874 | 10 | 3 | 113 | 3 | 3 |
| Thalamus | PCN | 17 | 3 | 2 | 43 | 6 | 3 |
| Cortex | PL | 1051 | 18 | 4 | 1048 | 16 | 4 |
| Thalamus | PO | 5 | 1 | 1 | 49 | 1 | 1 |
| Thalamus | PVT | 1 | 1 | 1 | 20 | 2 | 2 |
| Cortex | RSP | 232 | 6 | 2 | 122 | 6 | 3 |
| Thalamus | RT | 8 | 2 | 2 | 23 | 2 | 1 |
| Thalamus | SMT | 13 | 3 | 2 | 37 | 3 | 2 |
| Cortex | SSp | 289 | 9 | 3 | 353 | 11 | 4 |
| Striatum | STR | 73 | 4 | 3 | 63 | 4 | 3 |
| Thalamus | TH | 22 | 5 | 3 | 16 | 4 | 3 |
| Thalamus | VAL | 129 | 3 | 2 | 61 | 5 | 4 |
| Thalamus | VPM | 15 | 1 | 1 | 12 | 1 | 1 |
| Other | WM | 396 | 19 | 4 | 175 | 18 | 4 |
| Other | Other | 737 | 17 | 4 | 282 | 13 | 4 |

**Table S6. Neurons per region in manipulation experiments**

Number of neurons per region in *Grin2a*$^{+/-}$ animals across both manipulation types. Only regions that were recorded from in both manipulations included in this list. Region abbreviations in **Table S3**.

**Table S7. Statistics**

Statistical overview.

| Figure | Sample Size | Summary | Statistical Test | Values |
|---|---|---|---|---|
| 2c<br>Rate – All Areas | *Grin2a*$^{+/+}$: $n$ = 21,197<br>*Grin2a*$^{+/-}$: $n$ = 21,455 | | Two-Way Anova<br><br>Genotype*Brain Area | Area: $F(4, 42632) = 973.571$, $p < 0.001$<br>Genotype: $F(1, 42632) = 57.488$, $p$ = 3.47e-14<br>Interaction: $F(4, 42632) = 12.103$, $p$ = 7.84e-10 |
| 2e<br>Rate – All Regions | *Grin2a*$^{+/+}$: $n$ = 21,197<br>*Grin2a*$^{+/-}$: $n$ = 21,455 | | Two-Way Anova<br><br>Genotype*Brain Region | Region: $F(44, 42552) = 114.554$, $p < 0.001$<br>Genotype: $F(1, 42552) = 19.898$, $p$ = 8.19e-06<br>Interaction: $F(44, 42552) = 4.793$, $p$ = 1.42e-23 |
| 2h<br>Rate – Individual Regions | MD: *Grin2a*$^{+/+}$: $n$ = 799<br>*Grin2a*$^{+/-}$: $n$ = 636<br>ORB: *Grin2a*$^{+/+}$: $n$ = 133<br>*Grin2a*$^{+/-}$: $n$ = 320<br>MOs: *Grin2a*$^{+/+}$: $n$ = 3385<br>*Grin2a*$^{+/-}$: $n$ = 2848<br>LS: *Grin2a*$^{+/+}$: $n$ = 130<br>*Grin2a*$^{+/-}$: $n$ = 70<br>CM: *Grin2a*$^{+/+}$: $n$ = 28<br>*Grin2a*$^{+/-}$: $n$ = 38<br>Other: *Grin2a*$^{+/+}$: $n$ = 624<br>*Grin2a*$^{+/-}$: $n$ = 654<br>AM: *Grin2a*$^{+/+}$: $n$ = 46<br>*Grin2a*$^{+/-}$: $n$ = 38<br>IL: *Grin2a*$^{+/+}$: $n$ = 84<br>*Grin2a*$^{+/-}$: $n$ = 286<br>PCN: *Grin2a*$^{+/+}$: $n$ = 173<br>*Grin2a*$^{+/-}$: $n$ = 134<br>PO: *Grin2a*$^{+/+}$: $n$ = 456<br>*Grin2a*$^{+/-}$: $n$ = 240 | | Mann-Whitney $U$-Test | MD: $U = 618188$, $p$ = 5.36e-9<br>ORB: $U = 36405$, $p$ = 9.75e-07<br>MOs: $U = 10856426$, $p$ = 1.59e-05<br>LS: $U = 14664$, $p$ = 4.23e-05<br>CM: $U = 1238$, $p = 0.0001$<br>Other: $U = 421017$, $p = 0.000865$<br>AM: $U = 2310$, $p = 0.00144$<br>IL: $U = 121925$, $p = 0.00205$<br>PCN: $U = 28852$, $p = 0.00418$<br>PO: $U = 151967$, $p = 0.00585$ |

| | | | | |
|---|---|---|---|---|
| 3e<br>Co-Activity – All Regions | *Grin2a*$^{+/+}$: *n* = 21,197<br>*Grin2a*$^{+/-}$: *n* = 21,455 | | Two-Way Anova<br><br>Genotype*Brain Region | Region:<br>*F*(39, 2223804) = 1289.365, *p* = 0.001<br>Genotype:<br>*F*(1, 2223804) = 38.110, *p* = 6.69e-10<br>Interaction: *F*(39, 2223804) = 219.209, *p* = 0 |
| 3h<br>Co-Activity – Individual Regions | DG: *Grin2a*$^{+/+}$: *n* = 36185<br>*Grin2a*$^{+/-}$: *n* = 7426<br>MOs: *Grin2a*$^{+/+}$: *n* = 309614<br>*Grin2a*$^{+/-}$: *n* = 207581<br>PO: *Grin2a*$^{+/+}$: *n* = 26650<br>*Grin2a*$^{+/-}$: *n* = 7407<br>RT: *Grin2a*$^{+/+}$: *n* = 1540<br>*Grin2a*$^{+/-}$: *n* = 5914<br>CP: *Grin2a*$^{+/+}$: *n* = 164384<br>*Grin2a*$^{+/-}$: *n* = 165516<br>SSp: *Grin2a*$^{+/+}$: *n* = 29899<br>*Grin2a*$^{+/-}$: *n* = 20164<br>ACA: *Grin2a*$^{+/+}$: *n* = 4323<br>*Grin2a*$^{+/-}$: *n* = 2229<br>LP: *Grin2a*$^{+/+}$: *n* = 1567<br>*Grin2a*$^{+/-}$: *n* = 1148<br>CL: *Grin2a*$^{+/+}$: *n* = 6216<br>*Grin2a*$^{+/-}$: *n* = 19092<br>RSP: *Grin2a*$^{+/+}$: *n* = 14767<br>*Grin2a*$^{+/-}$: *n* = 24101 | | Unpaired, Two-tailed Student's *t*-test | DG: t(43609) = -33.992, p = 5.436e-250<br>MOs: t(517193) = -31.147, p = 8.901e-213<br>PO: t(34055) = -25.497, p = 4.626e-142<br>RT: t(7452) = 26.163, p = 2.005e-144<br>CP: t(329898) = -17.662, p = 8.787e-70<br>SSp: t(50061) = 15.595, p = 1.068e-54<br>ACA: t(6550) = -15.511, p = 2.59e-53<br>LP: t(2713) = -9.905, p = 9.611e-23<br>CL: t(25306) = -5.058, p = 4.277e-07<br>RSP: t(38866) = -5.214, p = 1.861e-07 |
| 4x<br>Rate Rescue – All Regions | | | | Region:<br>*F*(26, 22262) = 41.330, *p* = 9.22e-205<br>Genotype:<br>*F*(1, 22262) = |

| | | | | |
|---|---|---|---|---|
| | | | | 12.728, $p$ = 0.000361<br>Interaction: $F(26, 22262) = 9.652$, $p$ = 1.99e-38 |
| 4b<br>Injection Time | *Grin2a*$^{+/-}$ Dq: $n$ = 21<br>*Grin2a*$^{+/-}$mCh: $n$ = 20 | | Unpaired, two-tailed t-test | $t(39) = 0.568$, $p$ = 0.574 |
| 4d<br>HO Thalamus - Rates | *Grin2a*$^{+/+}$: $n$ = 1000<br>*Grin2a*$^{+/-}$: $n$ = 808<br>*Grin2a*$^{+/-}$ Dq: $n$ = 290<br>*Grin2a*$^{+/-}$ mCh: $n$ = 545 | | One-Way ANOVA | ANOVA: $F(3, 2639) = 28.882$, $p$ = 2.27e-18<br><br>xxx |
| 4e<br>Firing Rate | MOs:<br>*Grin2a*$^{+/+}$: $n$ = 3385<br>*Grin2a*$^{+/-}$: $n$ = 2848<br>*Grin2a*$^{+/-}$ Dq: $n$ = 1245 | | One-Way ANOVA | $F(3, 8386) = 4.231$, $p = 0.005$ |
| 4f<br>Firing Rate | ORB:<br>*Grin2a*$^{+/+}$: $n$ = 133<br>*Grin2a*$^{+/-}$: $n$ = 320<br>*Grin2a*$^{+/-}$ Dq: $n$ = 730 | | One-Way ANOVA | $F(3, 1250) = 5.692$, p = 0.00072 |
| 4g<br>Firing Rate | LS:<br>*Grin2a*$^{+/+}$: $n$ = 130<br>*Grin2a*$^{+/-}$: $n$ = 70<br>*Grin2a*$^{+/-}$ Dq: $n$ = 69 | | One-Way ANOVA | $F(3, 339) = 18.744$, p = 2.85e-11 |
| 4h<br>Firing Rate | As 4e-f | | Mann-Whitney $U$ test, uncorrected | MOs: 1, $p = 0.0087$<br>ORB: 0.4, $p$ = 0.0003<br>LS: 0, $p = 0.0021$<br>Other: 1, $p$ = 0.0001<br>IL: 0.72, $p = 0.019$<br>CA2: 1, $p = 0.0013$<br>MOp: 0.86, $p$ = 0.074<br>SSp: 1, $p = 0.0044$ |
| 4k<br>Co-activity | ORB<br>*Grin2a*$^{+/+}$: $n$ = 3868<br>*Grin2a*$^{+/-}$: $n$ = 21584<br>*Grin2a*$^{+/-}$ Dq: $n$ = 70459 pairs | | One-Way ANOVA | $F(3, 164948)$ = 43.841, $p$ = 2.61e-28 |
| 4l<br>Co-activity | DG<br>*Grin2a*$^{+/+}$: $n$ = 36185<br>*Grin2a*$^{+/-}$: $n$ = 7426 | | One-Way ANOVA | $F(3, 50509)$ = 400.199, $p$ = 6.04e-257 |

| | | | | |
|---|---|---|---|---|
| | *Grin2a*$^{+/-}$ Dq: $n$ = 3479 | | | |
| 4m<br>Co-activity | SSP<br>*Grin2a*$^{+/+}$: $n$ = 29899<br>*Grin2a*$^{+/-}$: $n$ = 20164<br>*Grin2a*$^{+/-}$ Dq: $n$ = 4769 | | One-Way ANOVA | $F$(3, 59597) = 169.025, $p$ = 4.07e-109 |
| S1i | *Grin2a*$^{+/+}$: $n$ = 8<br>*Grin2a*$^{+/-}$: $n$ = 8 | | Unpaired, two-tailed $t$-test | $t$(14) = 0.006, $p$ = 0.996 |
| S1j | *Grin2a*$^{+/+}$: $n$ = 47<br>*Grin2a*$^{+/-}$: $n$ = 46 | | Unpaired, two-tailed $t$-test | $t$(91) = 1.388, $p$ = 0.168 |
| S1k | *Grin2a*$^{+/+}$: $n$ = 45<br>*Grin2a*$^{+/-}$: $n$ = 45 | | Unpaired, two-tailed $t$-test | $t$(88) = 3.229, $p$ = 0.002** |
| S1l | *Grin2a*$^{+/+}$: $n$ = 13322<br>*Grin2a*$^{+/-}$: $n$ = 10383 | | Kolmogorov-Smirnov test | KS = 0.139, $p$ < 0.001 |
| S4c | *Grin2a*$^{+/+}$: $n$ = 21,197<br>Cortex: $n$ = 10,163<br>Hippocampus: $n$ = 2,563<br>Striatum: $n$ = 2,586<br>Thalamus: $n$ =3,399<br>Other: $n$ = 2,486 | | One-Way ANOVA | $F$(4, 21192) = 547.902, $p$ < 0.001 |
| S4d | As S4c | | One-Way ANOVA | $F$(4, 21192) = 589.931, $p$ < 0.001 |
| S4e | As S4c | | One-Way ANOVA | $F$(4, 21192) = 52.604, $p$ < 0.001 |
| S4f | As S4c | | One-Way ANOVA | $F$(4, 21192) = 427.136, $p$ < 0.001 |
| S4g | As S4c | | One-Way ANOVA | $F$(4, 21192) = 229.848, $p$ < 0.001 |
| S4h | As S4c | | One-Way ANOVA | $F$(4, 21192) = 283.445, $p$ < 0.001 |
| S4i | As S4c | | One-Way ANOVA | $F$(4, 21192) = 794.532, $p$ < 0.001 |
| S4j | As S4c | | One-Way ANOVA | $F$(4, 21192) = 429.971, $p$ < 0.001 |
| S5a | *Grin2a*$^{+/+}$: $n$ = 21,197 | | Two-Way Anova | Area: $F$(4, 42632) = 973.571, $p$ = 0 |

| | | | | |
|---|---|---|---|---|
| | *Grin2a*$^{+/-}$: $n$ = 21,455 | | Genotype*Brain Area | Genotype: $F(1, 42632) = 57.488$, $p = 3.47e-14$ Interaction: $F(4, 42632) = 12.103$, $p = 7.84e-10$ |
| S5b | *As S5a* | | Two-Way Anova<br><br>Genotype*Brain Area | Area: $F(4, 63{,}611) = 1639.788$, $p = 0$ Genotype: $F(1, 63611) = 84.551$, $p = 3.85e-20$ Interaction: $F(4, 63611) = 18.621$, $p = 2.62e-15$ |
| S5e | *As S5a* | Standard | Two-Way Anova<br><br>Genotype*Brain Area | Area: F(4, 61620) = 1198.179, p = 0 Genotype: F(2, 61620) = 28.299, p = 5.2e-13 Interaction: F(8, 61620) = 11.720, p = 8.27e-17 |
| S5f | *As S5a* | Probe | Two-Way Anova<br><br>Genotype*Brain Area | BrainArea main effect: F(4, 61620) = 1096.572, p = 0 Genotype main effect: F(2, 61620) = 28.075, p = 6.5e-13 BrainArea × Genotype interaction: F(8, 61620) = 9.047, p = 1.68e-12 |
| S5g | *As S5a* | Tone Only | Two-Way Anova<br><br>Genotype*Brain Area | Area: $F(4, 42632) = 1088.096$, $p = 0$ Genotype: $F(1, 42632) = 16.244$, $p = 8.86e-08$ Interaction: $F(4, 42632) = 5.822$, $p = 1.86e-07$ |
| S6c | *Grin2a*$^{+/+}$: $n$ = 21,197<br>*Grin2a*$^{+/-}$: $n$ = 21,455 | | Two-Way Anova<br><br>Genotype*Brain Area | Area: $F(4, 42632) = 609.5056$, $p < 0.001$ Genotype: $F(1, 42632) = 19.063$, $p = 1.27e-5$ Interaction: $F(4, 42632) = 21.678$, $p = 6.85e-18$ |

| | | | | |
|---|---|---|---|---|
| S6d | *As S6c* | | Two-Way Anova<br><br>Genotype*Brain Area | Area: $F(4, 42632) = 590.812$, $p < 0.001$<br>Genotype: $F(1, 42632) = 0.295$, $p = 0.587$<br>Interaction: $F(4, 42632) = 2.515$, $p = 0.039$ |
| S6e | *As S6c* | | Two-Way Anova<br><br>Genotype*Brain Area | Area: $F(4, 42632) = 1500.295$, $p < 0.001$<br>Genotype: $F(1, 42632) = 14.837$, $p < 0.001$<br>Interaction: $F(4, 42632) = 3.545$, $p = 0.007$ |
| S6f | *As S6c* | | Two-Way Anova<br><br>Genotype*Brain Area | Area: $F(4, 42632) = 809.508$, $p < 0.001$<br>Genotype: $F(1, 42632) = 3.162$, $p = 0.075$<br>Interaction: $F(4, 42632) = 5.286$, $p < 0.001$ |
| S10d | *Grin2a*$^{+/+}$: $n = 21{,}197$<br>*Grin2a*$^{+/-}$: $n = 21{,}455$ | | Two-Way Anova<br><br>Genotype*Brain Area | Area: $F(6, 42628) = 649.417$, $p < 0.001$<br>Genotype: $F(1, 42632) = 4.621$, $p = 0.031$<br>Interaction: $F(4, 42632) = 8.681$, $p = 1.81e-09$ |
| S11b | *Grin2a*$^{+/+}$: $n = 21{,}197$<br>*Grin2a*$^{+/-}$: $n = 21{,}455$ | CV | Two-Way Anova<br><br>Genotype*Brain Region | Region: F(38, 61431) = 22.939, $p = 4.58e-157$<br>Genotype: F(1, 61431) = 0.842, $p = 0.359$<br>Interaction: F(82, 61431) = 7.454, $p = 2.14e-81$ |
| S11d | *As S11b* | CV | Two-Way Anova<br><br>Genotype*Brain Area | Region: F(4, 61620) = 101.981, $p = 1.05e-86$<br>Genotype: F(2, 61620) = 2.238, $p = 0.107$ |

| | | | | |
|---|---|---|---|---|
| | | | | Interaction: F(8, 61620) = 4.926, *p* = 4.14e-06 |
| S11f | *As S11b* | CV2 | Two-Way Anova<br><br>Genotype*Brain Region | Region: F(38, 61431) = 136.831, *p* < 0.001<br>Genotype: F(1, 61431) = 5.894, *p* = 0.0152<br>Interaction: F(82, 61431) = 5.965, *p* = 6.06e-59 |
| S11h | *As S11b* | CV2 | Two-Way Anova<br><br>Genotype*Brain Area | Region: F(4, 61620) = 973.220, p = 0<br>Genotype: F(2, 61620) = 11.482, p = 1.03e-05<br>Interaction: F(8, 61620) = 17.198, p = 8.09e-26 |
| S12e | *Grin2a*$^{+/+}$: *n* = 21,197<br>*Grin2a*$^{+/-}$: *n* = 21,455 | Burst Firing | Two-Way Anova<br><br>Genotype*Brain Area | Area: *F*(4, 42632) = 984.111, *p* = 0<br>Genotype: *F*(1, 42632) = 1.483, *p* = 0.223<br>Interaction: *F*(4, 42632) = 16.534, *p* = 1.52e-13 |
| S12e | *As S12e* | | Two-Way Anova<br><br>Genotype*Brain Region | Area: *F*(44, 42552) = 100736, *p* = 0<br>Genotype: *F*(1, 42554) = 0.096, *p* = 0.756<br>Interaction: *F*(4, 42632) = 4.874 *p* = 3.37e-24 |
| S12g | RSP: *Grin2a*$^{+/+}$: *n* = 509 neurons<br>*Grin2a*$^{+/-}$: *n* = 564<br>ORB: *Grin2a*$^{+/+}$: *n* = 133 neurons<br>*Grin2a*$^{+/-}$: *n* = 320<br>MOp: *Grin2a*$^{+/+}$: *n* = 2767 neurons<br>*Grin2a*$^{+/-}$: *n* = 3318 neurons<br>ACA: *Grin2a*$^{+/+}$: *n* = 297 neurons<br>*Grin2a*$^{+/-}$: *n* = 197 neurons | | Mann-Whitney *U*-Test | RSP: *U* = 24317, *p* = 2.58e-9<br>ORB: *U* = 37480, *p* = 8.4e-09<br>MOp: *U* = 8061526, *p* = 1.47e-07<br>ACA: *U* = 65760, *p* = 5.91e-07<br>LS: *U* = 1.471750e+04, z = 4.233, p = 2.3e-05<br>VIS: *U* = 5782, z = 3.715, *p* = 0.000203 |

| | | | | |
|---|---|---|---|---|
| | LS: *Grin2a*$^{+/+}$: $n$ = 130 neurons<br>*Grin2a*$^{+/-}$: $n$ = 70 neurons<br>VIS: *Grin2a*$^{+/+}$: $n$ = 84 neurons<br>*Grin2a*$^{+/-}$: $n$ = 37 neurons<br>MOs: *Grin2a*$^{+/+}$: $n$ = 3385 neurons<br>*Grin2a*$^{+/-}$: $n$ = 2848 neurons<br>DP: *Grin2a*$^{+/+}$: $n$ = 141 neurons<br>*Grin2a*$^{+/-}$: $n$ = 81 neurons<br>AUD: *Grin2a*$^{+/+}$: $n$ = 1309 neurons<br>*Grin2a*$^{+/-}$: $n$ = 1201 neurons<br>IL: *Grin2a*$^{+/+}$: $n$ = 84 neurons<br>*Grin2a*$^{+/-}$: $n$ = 286 neurons<br>Other: *Grin2a*$^{+/+}$: $n$ = 624 neurons<br>*Grin2a*$^{+/-}$: $n$ = 654 neurons | | | MOs: $U$ = 10319018, z = -3.283, $p$ = 0.00103<br>DP: $U$ = 16747, z = 2.226, $p$ = 0.026<br>AUD: $U$ = 1605942, z = -2.070, p = 0.0385<br>IL: $U$ = 13860, z = -2.009, p = 0.0446<br>Other: $U$ = 386175, z = -1.953, $p$ = 0.050 |
| S21n | ORB<br>*Grin2a*$^{+/-}$: $n$ = 320 neurons<br>*Grin2a*$^{+/-}$ mCh: $n$ = 719 neurons | | Mann-Whitney *U*-Test | $U$ = 152198, z = -3.180, $p$ = 0.00147<br>+35.4% |
| | MOs<br>*Grin2a*$^{+/-}$: $n$ = 1235 neurons<br>*Grin2a*$^{+/-}$ mCh: $n$ = 1389 neurons | | Mann-Whitney *U*-Test | $U$ = 5932769, z = -1.952, $p$ = 0.0509<br>(Δ = +12.51%) |
| | LS<br>*Grin2a*$^{+/-}$: $n$ = 1515 neurons<br>*Grin2a*$^{+/-}$ mCh: $n$ = 1640 neurons | | Mann-Whitney *U*-Test | $U$ = 3576, z = -1.124, $p$ = 0.261<br>(Δ = 8.28%) |
| | Other<br>*Grin2a*$^{+/-}$: $n$ = 1613 neurons<br>*Grin2a*$^{+/-}$ mCh: $n$ = 3179 neurons | | Mann-Whitney *U*-Test | Mann–Whitney U = 366146, z = -5.672, p = 1.41e-08<br>(Δ = 97.12%) |

| | | | | |
|---|---|---|---|---|
| | IL<br>*Grin2a*$^{+/-}$: $n$ = 1266 neurons<br>*Grin2a*$^{+/-}$ mCh: $n$ = 1143 neurons | | Mann-Whitney *U*-Test | Mann–Whitney U = 65699, z = 0.150, p = 0.881<br>(Δ = -9.71%) |
| S23b | MOs | RS | Mann-Whitney *U*-Test | F(3, 7538) = 3.697, *p* = 0.012.<br><br>Mann–Whitney U = 4350782, p = 0.0282<br>(Δ = 15.5%) |
| | ORB | RS | Mann-Whitney *U*-Test | F(3, 1113) = 3.352, *p* = 0.018.<br><br>Mann–Whitney U = 114253, p = 0.004<br>(Δ = 30.5%) |
| | Other | RS | Mann-Whitney *U*-Test | F(3, 1550) = 9.011, *p* = 6.46e-06<br><br>Mann–Whitney U = 144826, p = 0.0001<br>(Δ = 87.5%) |
| | MOp | RS | Mann-Whitney *U*-Test | F(3, 6332) = 2.040, *p* = 0.106.<br><br>Mann–Whitney U = 4664068, p = 0.325<br>(Δ = 4%) |
| | IL | RS | Mann-Whitney *U*-Test | F(3, 1070) = 1.498, *p* = 0.213<br><br>Mann–Whitney *U* = 16710, p = 0.0133<br>(Δ = 58.8%) |
| S23d | ORB | FS | Mann-Whitney *U*-Test | F(3, 175) = 6.474, *p* = 0.000352<br><br>Mann–Whitney *U* = 2370, p = 0.0001<br>(Δ = 340.9%) |
| | AUD | FS | Mann-Whitney *U*-Test | F(3, 669) = 4.892, *p* = 0.00227 |

| | | | | |
|---|---|---|---|---|
| | | | | Mann–Whitney *U* = 31557, p = 0.1835 (Δ = 38.2%) |
| | DP | FS | Mann-Whitney *U*-Test | *F*(3, 79) = 1.212, *p* = 0.311<br><br>Mann–Whitney *U* = 549, *p* = 0.08 (Δ = 370.2%) |
| | RSP | FS | Mann-Whitney *U*-Test | F(3, 328) = 0.589, *p* = 0.311<br><br>Mann–Whitney *U* = 10976, *p* = 0.0081 (Δ = 96.1%) |
| | MOs | FS | Mann-Whitney *U*-Test | F(3, 1195) = 11.833, *p* = 1.22e-07<br><br>Mann–Whitney *U* = 92280, *p* = 0.0001 (Δ = 82.9%) |
| S24c | dCA3 | | One-Way ANOVA | F(3, 960) = 3.697, *p* = 0.012 |
| | dDG | | One-Way ANOVA | F(3, 705) = 0.941, *p* = 0.420 |
| | dCA1 | | One-Way ANOVA | F(3, 917) = 0.719, *p* = 0.541 |
| | dCA2 | | One-Way ANOVA | *F*(3, 117) = 1.606, *p* = 0.192 |
| S24e | vCA3 | | One-Way ANOVA | *F*(3, 1521) = 0.722, *p* = 0.539 |
| | vDG | | One-Way ANOVA | *F*(3, 953) = 0.521, *p* = 0.668 |
| | vCA1 | | One-Way ANOVA | *F*(3, 1633) = 0.319, *p* = 0.812 |
| | vCA2 | | One-Way ANOVA | *F*(3, 62) = 0.951, *p* = 0.422 |
| S24h | dCA3 | | One-Way ANOVA | F(3, 960) = 2.687, *p* = 0.045 |
| | dDG | | One-Way ANOVA | F(3, 705) = 4.436, *p* = 0.004 |
| | dCA1 | | One-Way ANOVA | F(3, 917) = 0.719, *p* = 0.541 |
| | dCA2 | | One-Way ANOVA | NA |

| | | | | |
|---|---|---|---|---|
| S24j | vCA3 | | One-Way ANOVA | $F(3, 1521) = 0.205$, $p = 0.893$ |
| | vDG | | One-Way ANOVA | $F(3, 953) = 4.715$, $p = 0.003$ |
| | vCA1 | | One-Way ANOVA | $F(3, 1633) = 0.319$, $p = 0.812$ |
| | vCA2 | | One-Way ANOVA | $F(3, 62) = 9.490$, $p < 0.001$ |
| S25b | ACA | | One-Way ANOVA | $F(3, 1133) = 4.824$, $p = 0.00242$ |
| S25c | RSP | | One-Way ANOVA | $F(3, 1359) = 14.812$, $p = 1.71e-09$ |
| S25d | MOp | | One-Way ANOVA | $F(3, 7435) = 6.597$, $p = 0.00019$ |
| S25e | RSP<br>ORB<br>Mop<br>ACA<br>LS<br>VIS<br>MOs<br>DP<br>AUD<br>IL<br>Other | | | RSP: 0.94, $p = 0.0068$<br>ORB: 0.09, $p = 0.255$<br>MOp: 0, $p = 0.0508$<br>ACA: 1, $p = 0.0000$<br>LS: 0, $p = 0.0038$<br>VIS: not rec.<br>MOs: 0, $p = 0.0022$<br>DP: 0.07, $p = 0.717$<br>AUD: 0.53, $p = 0.0374$<br>IL: 0.87, $p = 0.0691$<br>Other: 0, $p = 0.065$ |

Region abbreviations in **Table S3**.
